# Designed Ankyrin Repeat Proteins for detecting prostate-specific antigen expression *in vivo*

**DOI:** 10.1101/2023.01.24.525357

**Authors:** Melanie Gut, Birgit Dreier, Sven Furler, Jens Sobek, Andreas Plückthun, Jason P. Holland

**Author notes:** **Corresponding Author:** Prof. Dr Jason P. Holland, Website: www.hollandlab.org, Twitter: @HollandLab_.

## Abstract

Late-stage prostate cancer often acquires resistance to conventional chemotherapies and transforms into a hormone-refractory, drug-resistant, and non-curative disease. Developing non-invasive tools to detect the biochemical changes that correlate with drug efficacy and reveal the onset of drug resistance would have important ramifications in managing the treatment regimen for individual patients. Here, we report the selection of new Designed Ankyrin Repeat Proteins (DARPins) that show high affinity toward prostate-specific antigen (PSA), a biomarker used in clinical monitoring of prostate cancer. Ribosome display and *in vitro* screening tools were used to select PSA-binding DARPins based on their binding affinity, selectivity, and chemical constitution. Surface plasmon resonance measurements demonstrated that the four lead candidates bind to PSA with nanomolar affinity. DARPins were site-specifically functionalised at a unique *C*-terminal cysteine with the hexadentate *aza*-nonamacrocyclic chelate (NODAGA) for subsequent radiolabelling with the positron-emitting radionuclide ^68^Ga. [^68^Ga]GaNODAGA-DARPins showed high stability toward transchelation and were stable in human serum for >2 h. Radioactive binding assays using streptavidin-loaded magnetic beads confirmed that the functionalisation and radiolabelling did not compromise the specificity of [^68^Ga]GaNODAGA-DARPins toward PSA. Biodistribution experiments in athymic nude mice bearing subcutaneous prostate cancer xenografts derived from the LNCaP cell line revealed that three of the four [^68^Ga]GaNODAGA-DARPins displayed specific tumour-binding *in vivo*. For DARPin-**6**, tumour-uptake in the normal group reached 4.16 ± 0.58 %ID g^-1^ (*n* = 3; 2 h post-administration) and was reduced by ∼50% in the blocking group (2.47 ± 0.42 %ID g^-1^; *n* = 3; *P*-value = 0.018). Collectively, the experimental results support the future development of new PSA-specific imaging agents for potential use in monitoring the efficacy of androgen receptor (AR)-targeted therapies.

## Introduction

Prostate cancer (PCa) is the second most common cancer diagnosed in men world-wide.^1^ Cellular signalling events derived from activation of the androgen receptor (AR) pathway play a prominent role in PCa development and progression.^2,3^ Consequently, first-line treatment options for PCa patients in the clinic have focused on androgen deprivation therapy (ADT) which includes androgen ablation and AR inhibition.^3–6^ Ablation reduces androgen levels by using either surgical or chemical castration to interfere with androgen biosynthesis, whereas the objective of AR inhibition is to disrupt the signalling pathway and block the transcription of pro-oncogenic factors. First-generation AR inhibitors, including flutamide, nilandron, and bicalutamide,^5–8^ remain the mainstay of PCa chemotherapy but other drugs have emerged including the second-generation AR inhibitor MDV3100 (enzalutamide),^9,10^ and the CYP17A1 inhibitor abiraterone acetate (Zytiga),^11,12^ which suppresses androgen synthesis (**Figure 1**). Despite the many advances in chemotherapy, most PCa tumours develop resistance over time. The mechanisms that lead to drug resistance are not fully known but mutations that restore AR signalling can produce a hormone-refractory disease which is often referred to as castration-resistant prostate cancer (CRPC). Alternatively, other transformations associated with AR-independent signalling pathways have also been implicated in regulating the metastatic potential of PCa.^13–17^ Despite numerous second-generation drugs to treat CRPC, no curative treatment exists.^8,10,12,17,18^

**Figure 1.**
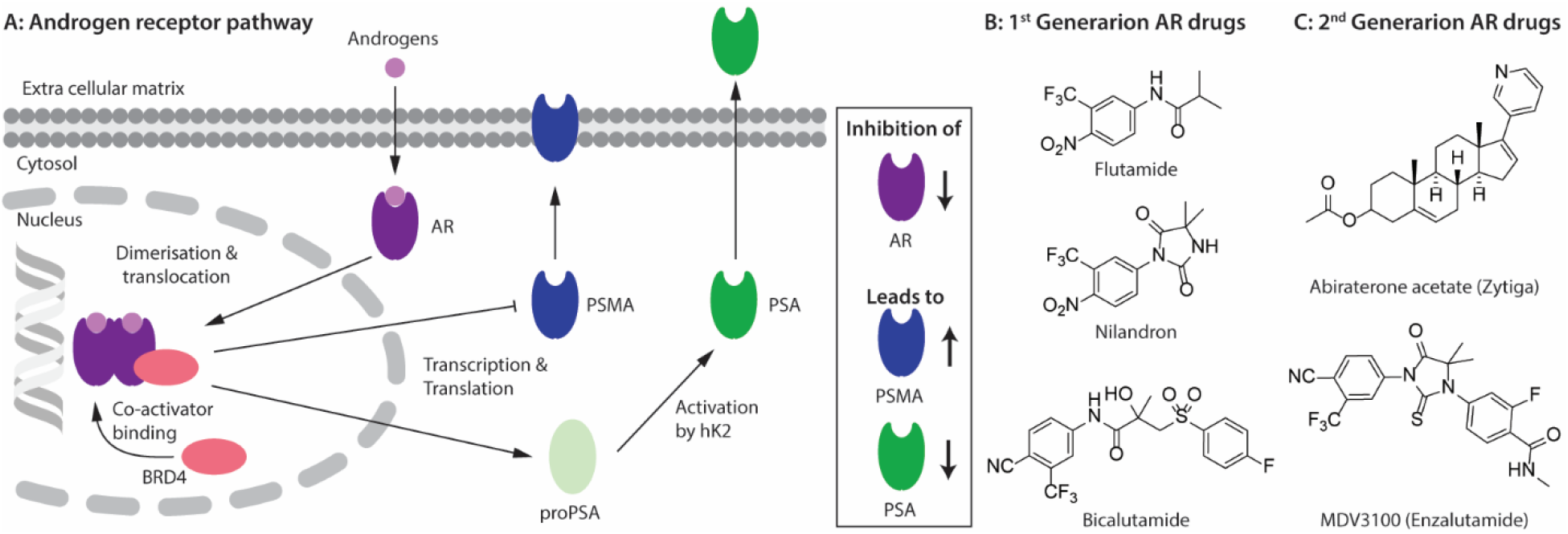
(A) Schematic showing the putative relationship between androgen-induced AR signalling and gene expression of the key PCa biomarkers, including PSMA and PSA.^2,3^ Chemical structures of prominent (B) 1^st^ generation, and (C) 2^nd^ generation PCa chemotherapeutics used in frontline clinical practice.^8^

The expression of several key protein biomarkers of PCa, including prostate-specific membrane antigen (PSMA) and prostate-specific antigen (PSA; also called gamma-seminoprotein or kallikrein 3 [KLK3]), have been shown to depend on AR activity (**Figure 1**). In some PCa phenotypes, AR acts as a transcription suppressor of PSMA, and ADT can lead to an increase in PSMA levels.^19–21^ In contrast, PSA levels correlate directly with AR transcription activity, where a decrease in the measured concentration of PSA provides a diagnostic readout of the efficacy of AR inhibition.^19–21^

Molecular imaging (MI) techniques such as positron emission tomography (PET) can visualise spatiotemporal distribution and quantify changes in biological processes at the tissue level, allowing disease progression or treatment efficacy to be monitored on a lesion-by-lesion basis.^22,23^ As PSMA and PSA have a strong correlation with the AR pathway^19–21^ they are promising targets for the use in molecular imaging and therapy. Numerous PET imaging agents that delineate PSMA expression have been tested in the clinic, and clever design of the radiotracers has facilitated the transition toward PSMA-targeted radionuclide therapy.^24,25^ For example, in March 2022, the culmination of an international effort led to the United States Federal Drug Administration (US-FDA) approval of two PSMA ligands, ^68^Ga-PSMA-11 (^68^Ga-gozetotide, Locametz®, Novartis) for diagnosis^26,27^ and ^177^Lu-PSMA-617 (Pluvicto™, Novartis) for targeted β^−^ therapy of PSMA-positive tumours.^28^ Whilst high expression levels in most PCa phenotypes mean that PSMA is the current biomarker of choice when developing radiotracers for imaging and therapy, several drawbacks remain. First, PSMA expression is not entirely PCa-specific and high levels have been identified in the salivary glands, kidneys, nerve endings, and other sites of neoangiogenesis.^29,30^ This off-target expression is less problematic for imaging where low chemical and radiochemical doses are administered, but can cause severe, dose-limiting side-effects when using targeted radionuclide therapy. Second, the high baseline levels of PSMA expression, and the relatively weak inverse relation between AR signalling and PSMA expression is sub-optimal in the context of companion diagnostics where PSMA-biomarker quantification is used to monitor chemotherapeutic efficacy.^25^ Due to the high baseline, the increase in PSMA concentration upon drug treatment is potentially too ambiguous to measure drug efficacy reliably using PET.

In contrast to the expression profile of PSMA, PSA is almost exclusively found in prostate epithelial cells.^31,32^ PSA catalyses the proteolysis of gel proteins^33^ and is mainly excreted into the seminal fluid where concentrations can range from 0.1 to 2.2 mg mL^-1^.^34^ In the absence of disease, only a small fraction of PSA leaks into the circulatory system (∼0.5 ng mL^-1^ of total PSA in males ≤50 years old),^34^ where high concentrations of protease inhibitors, like the serpins α1-antichymotrypsin (ACT) and α2-macroglobilin (A2M), form a complex with the catalytically active PSA (“free” PSA or fPSA). Serpin binding blocks the enzymatic site, inactivates PSA, and produces complexed PSA (cPSA).^31,35,36^ Measurements of both fPSA and total PSA (tPSA) in the blood are used in the clinic as both predictive and prognostic biomarkers to monitor PCa progression.^37–39^ However, serum measurements of PSA are considered an imperfect biomarker of PCa, because blood pool secretion does not always reflect local expression levels in the tumours^31^ and has been shown to vary with age, body mass index, and the presence of co-morbidities.^31,40^ The mismatch between the comparatively low levels of circulatory PSA and the true levels expressed in PCa lesions can lead to errors in patient diagnosis and management. Therefore, complementary diagnostic techniques are required to enable a more accurate assessment of PCa progression. Non-invasive PET imaging of fPSA expression in PCa lesions has been proven to be a potential route toward monitoring treatment response in patients treated with frontline AR inhibitors.^40–42^ However, a current limitation in utilising non-invasive detection of PSA as a companion diagnostic is that very few drugs^43–45^ or radiotacers^41,42^ have been developed to target PSA with high affinity and specificity.

A first step toward fPSA targeting for non-invasive detection and radioimmunotherapy was achieved by utilising the humanised monoclonal antibody (mAb) hu5A10 to image (with ^89^Zr) and treat (with ^90^Y or ^225^Ac) PCa tumours in mice, and in macaques.^41,42^ Hu5A19 binds selectively and with a high specificity and affinity to the catalytic cleft of fPSA.^46,47^ The ^89^Zr-DFO-hu5A10 radiotracer showed high and specific uptake in LNCaP xenografts (22.75 ± 8.2% ID g^-1^) which increased after treatment with testosterone (35.33 ± 13.1% ID g^-1^) and decreased in a dose-dependent fashion after treatment with the AR antagonist MDV3100 (3.19 ± 1.4% ID g^-1^; 80 mg kg^-1^).^41^ These results confirmed the biochemical relationship between AR-signalling and PSA expression in the LNCaP model (**Figure 1**). Therapeutic effects were observed for ^90^Y- or ^225^Ac-labelled hu5A10 with increased median survival rates of 32 days (*n* = 6) for the control group *versus* 64 days (*n* = 6) and 188 days (*n* = 14) for groups of mice treated with ^90^Y-hu5A10 or ^225^Ac-hu5A10, respectively.^42^

In this work, our experimental goal was to identify new Designed Ankyrin Repeat Proteins (DARPins) for selective, high-affinity binding of PSA. DARPins are small (11–20 kDa), highly stable synthetic consensus proteins.^48–50^ Ankyrin repeat proteins are composed of stacked ankyrin motifs, consisting of a β-turn followed by anti-parallel α-helices. The *C*-terminal and *N*-terminal capping repeats (caps) shield a more hydrophobic internal core of the protein.^48–51^ Targeted engagement occurs primarily *via* interactions with the internal repeats, which carry randomised residues, and the sequence can be selected to provide high affinity and specificity toward a given target epitope.^50,52^ Several groups have developed radiotracers based on this protein platform.^50,53^ To illustrate the potential for clinical application of DARPin-based radiotracers, Bragina *et al*. reported data from a phase I trial which confirmed the safety and excellent performance of a ^99m^Tc-labelled DARPin for imaging human epidermal growth factor 2 expression in breast cancer patients.^53–56^ Here, we demonstrate the feasibility of developing new PCa diagnostic imaging agents based on the selection of PSA-specific DARPins.

## Results and discussion

### Selection of PSA-binding DARPins

Ribosome display is an efficient technique to select high-affinity DARPin binders from a randomised library.^49,50,52^ PSA-binding DARPins were selected in four rounds of ribosome display. Initial selections produced a long-list of 380 binders, which were evaluated in a Förster Resonance Energy Transfer (FRET) based assay,^52^ from which 32 FLAG-tagged DARPins (abbreviated as DARPins) were chosen, their sequence was determined and 27 single clones were selected for further analysis (see Methods in the Supplementary Materials). The selected DARPins were assessed for their binding affinity and selectivity toward PSA by using ELISA methods (**Figure S1**). The chromatographic profile of each DARPin measured by size-exclusion chromatography (24 mL Superdex 200 10/30 column) was used as a second selection criterion to refine the pool to a short-list of 12 PSA-binding DARPins. This short-list contained DARPins with a varying number of internal ankyrin repeat units (N = 1, 2, or 3), leading to different sizes of approximately 12, 15, or 19 kDa, respectively. In addition to the 12 different PSA-DARPins (**1** – **12**), a non-binding DARPin (**13**, E3_5^48^) was used as a negative control. A summary of the exclusion criteria and associated experimental data for each of the 27 DARPins is presented in **Table S1**.

### Cloning and purification of DARPins with a *C*-terminal Cys

Cysteine functionalisation is one of the most commonly applied protein conjugation methods.^57^ The Michael-addition reaction of a Cys sulfhydryl group with a reagent bearing an electrophilic maleimido-group is a well-established, reliable, and biocompatible way to site-specifically functionalise proteins.^57^ The DARPin libraries were previously engineered to omit cysteines from their core structure.^48^ Here, to enable site-specific modifications by using maleimide reagents, the 12 PSA-binding DARPins (**1** – **12**) and control DARPin **13** were cloned into a vector encoding a unique Cys residue located at the *C*-terminus (Sequences of the 13 DAPRins and DAPRin-Cys are displayed in **Figure S2** and **S3**). The vectors also contained an *N*-terminal His_6_-tag to allow protein purification by immobilised metal ion (Ni) affinity chromatography (IMAC). After ligation, the vectors were cloned into *E. coli* and the modified proteins were expressed, extracted, and purified.

After isolating the 12 DARPins, an ELISA was used to evaluate the effect of introducing the Cys at the *C*-terminus on PSA-binding. Data revealed that when comparing the measured absorbance (A_405-540 nm_) from the ELISA for the 12 PSA-binding DARPins either with *C*-terminal Cys (**Figure 2**, DARPin-Cys, cyan) or the initial FLAG-tag (**Figure 2**, DARPin, light blue), the addition of the bioconjugation handle had no impact on PSA binding. All 12 PSA-binding DARPin-Cys (**1** – **12**) proteins displayed specificity toward PSA. However, both forms of DARPins **4, 5, 7**, and **11** gave significantly lower absorbance, suggesting lower affinity binding, and were excluded from further studies. We also noted that DARPin **2** showed a higher level of non-specific binding, and therefore, this construct was also excluded.

**Figure 2.**
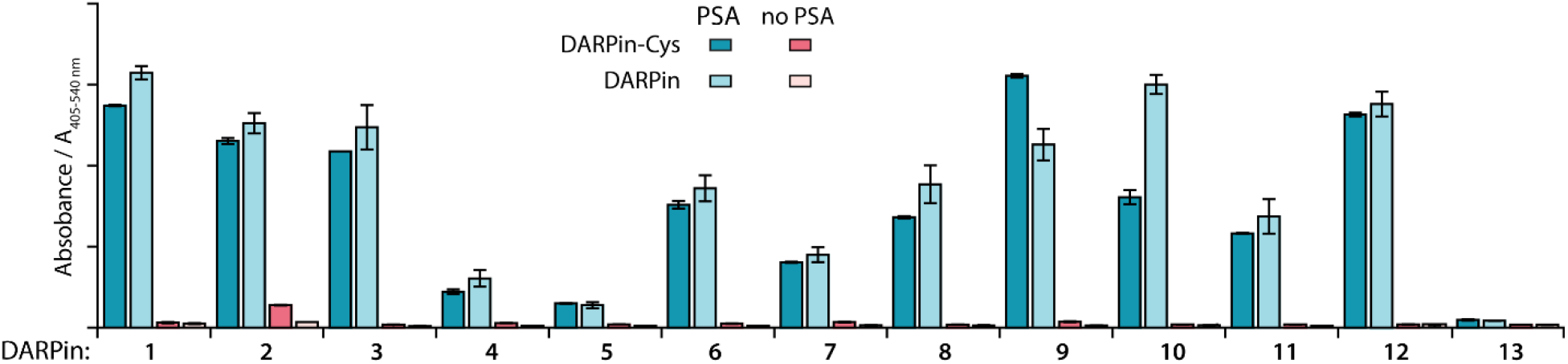
Bar chart with results of ELISA measurements showing the specific binding of the 12 selected DARPins towards PSA. Data presented show the absorbance (A_405-540 nm_) for PSA-binding DARPins (**1** – **12**) and the non-binding control (E3_5, **13**) either with *C*-terminal Cys modification (DARPin-Cys, cyan) or FLAG-tagged DARPins (light blue) a. Red bars show the non-specific binding control where no PSA was coated on the plate. Error bars represent one standard deviation about the mean (*n* = 2 measurements per sample).

In addition to the ELISA measurements, the DARPins-Cys constructs (**1** – **12**) were subjected to SDS-PAGE analysis (Supporting Information **Figure S4**) and size-exclusion chromatography coupled to a multi-angle light scattering (SEC-MALS) detector (Supporting Information **Figure S5**, and **Table S2**) to assess the molecular weight and purity of the protein samples. All DARPins-Cys proteins behaved as expected on the SDS-PAGE. Two constructs, DARPin-Cys **2** and **10**, gave a major band at the anticipated molecular weight but SDS-PAGE also revealed lower purity with the presence of several higher molecular weight proteins in the samples. SEC-MALS analysis revealed that DARPin-Cys **2** contained a significant fraction of dimeric (or aggregated) protein, and DARPin-Cys **4, 5** and **7** gave major peaks corresponding to low molecular weight species that were potentially protein fragments. Collectively, these experimental data supported the exclusion of DARPins **2, 4, 5, 7, 10**, and **11** from the pool.

### Functionalisation of DARPins-Cys proteins with a metal ion binding chelate

Of the remaining six PSA-binding DARPins-Cys proteins (**1, 3, 6, 8, 9**, and **12**), for practical and economic reasons, we made a further selection to reduce the number of constructs to four. DARPin-Cys **6** (19 kDa) and **9** (12 kDa) were chosen due to their different size arising from a different number of internal ankyrin repeats (N = 3, and 1 ankyrin repeats, respectively). In addition, we selected DARPins-Cys **3** and **12**, which are both of intermediate size (15 kDa with N = 2 ankyrin repeats) and showed high PSA-specific binding from the ELISA measurements. DARPins-Cys proteins **1** and **8** were randomly excluded from the study. The four PSA-binding DARPins-Cys proteins **3, 6, 9**, and **12**, and the non-binding control DARPin-Cys (**13**) were then site-specifically conjugated at the Cys residue by using a hexadentate *tris*-*aza*-nonamacrocyclic chelate (NODAGA)-maleimide (details are provided in Supporting Information **Table S3**). Prior to conjugation, potentially oxidised, dimeric DARPins-Cys species were reduced by incubating protein samples with an excess of dithiothreitol (DTT, 100 eq.), thereby ensuring that the Cys sulfhydryl groups were available for the Michael addition. After removal of the excess DTT by spin filtration, the DARPin-Cys samples (**3, 6, 9, 12**, and control **13**) were incubated with NODAGA-maleimide (2 eq.) in phosphate buffer (PBS, pH 7.9, 2 h, room temperature). Aliquots of the crude reaction mixtures were kept for radiochemical analysis. Excess or unreacted small-molecule reagents were removed by NAP-25 column purification, and subsequent spin concentration giving the purified chelate-functionalised NODAGA-DARPins in 21% to 56% yield (Supporting Information **Table S3**). We note that the variation in yields reflects the different behaviour of the NODAGA-DARPins on the purification column. Successful conjugation was confirmed by high-resolution mass spectrometry (HR-MS, Supporting Information **Table S4** and **Figures S6** – **S10**) and SDS-PAGE (**Figure 3**).

**Figure 3.**
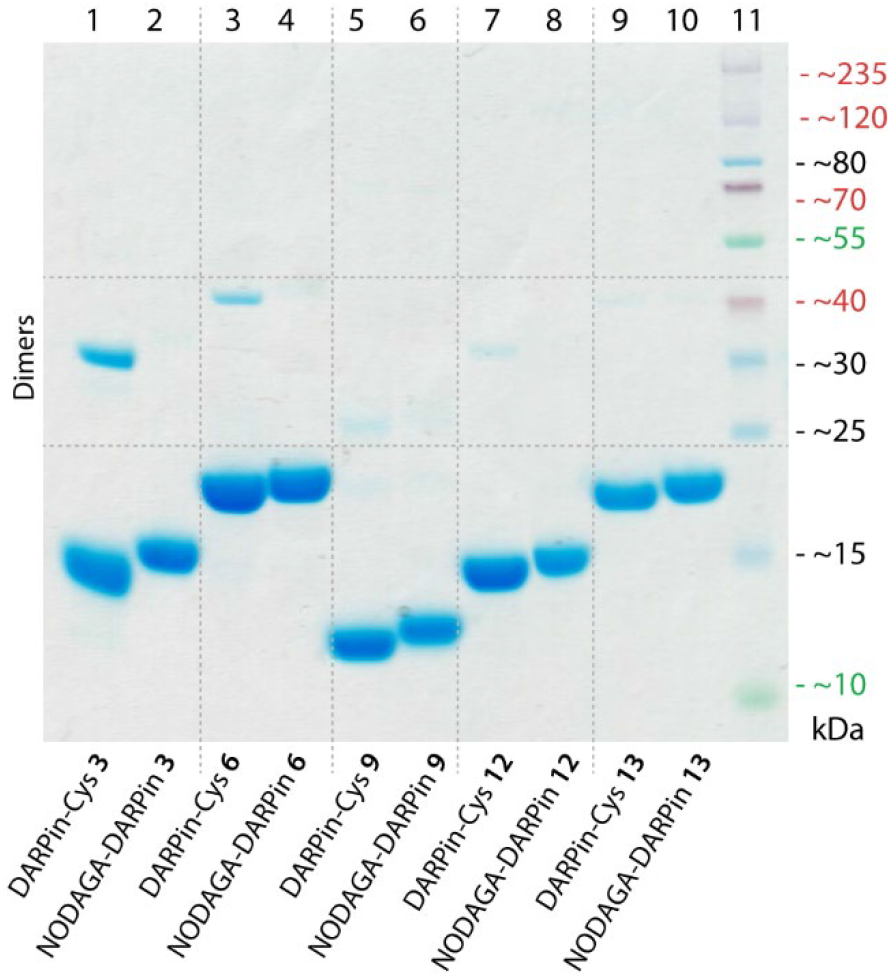
SDS-polyacrylamide gel showing DARPin-Cys **3, 6, 9, 12**, and **13** (lanes 1, 3, 5, 7, and 9, respectively) and the equivalent functionalised NODAGA-DARPins (lanes 2, 4, 6, 8 and 10, respectively). All NODAGA-modified constructs showed a small shift in retention due to the higher molecular weight. Unlike the DARPin-Cys protein samples, no dimers were detected for the NODAGA-DARPin samples even without addition of reductant in the loading buffer, indicating complete coupling. Note: Protein bands are stained with Coomassie blue, and the molecular weight ladder is shown in lane 11.

The chemical purity of all isolated PSA-binding NODAGA-DARPins (**3, 6, 9, 12**) was estimated to be >95% by SDS-PAGE and >98% by SEC (**Figure 3** and Supporting Information **Figure S12** light blue trace). From the SDS-PAGE data, shifts were observed in the band positions to slightly higher molecular weights for the NODAGA-DARPin products when compared with the non-functionalised DARPin-Cys constructs (**Figure 3**). SDS-PAGE also illustrated the different size of the NODAGA-DARPins (12, 15 and 19 kDa). Unlike the DARPin-Cys constructs, no bands associated with the protein dimerisation were observed in the SDS-PAGE of the NODAGA-DARPin samples, indicating that quantitative functionalisation of all available sulfhydryl residues was achieved. These observations were also supported by the HR-MS data from which only peaks associated with the correct isotopic distribution patterns of the functionalised NODAGA-DARPins were detected, with essentially no peaks present for the non-functionalised DARPin-Cys species. In all cases, covalent attachment of the NODAGA chelate to the proteins increased the molecular weight by *m*/*z* +497, concordant with the additional mass of the chelate and linker. Furthermore, upon addition of Ga(NO_3_)_3_ the expected shifts by *m*/*z* +66 (for the most abundant natural isotope: ^69^Ga, 60.11%) were measured in the HR-MS data of the ^nat^GaNODAGA-DARPin samples, confirming the successful metallation with ^nat^Ga^3+^ (Supporting Information **Table S4** and **Figure S6 – S10**).

### Qualitative assessment of NODAGA-DARPin PSA-binding by native PAGE and size exclusion chromatography

PSA-binding of the functionalised NODAGA-DARPins was assessed to confirm that the addition of the chelate did not interfere with target engagement. Aliquots of the NODAGA-DARPins (**3, 6, 9, 12**, and control **13**) were incubated with equimolar amounts of PSA at 24 °C for 30 min. For each of the NODAGA-DARPins (**3, 6, 9, 12**), native PAGE analysis showed a discrete shift in the protein band associated with formation of the PSA-DARPin complexes (**Figure 4**A). For the non-binding control NODAGA-DARPin **13**, no change was observed, and discrete bands were present for both free NODAGA-DARPin **13** and PSA.

**Figure 4.**
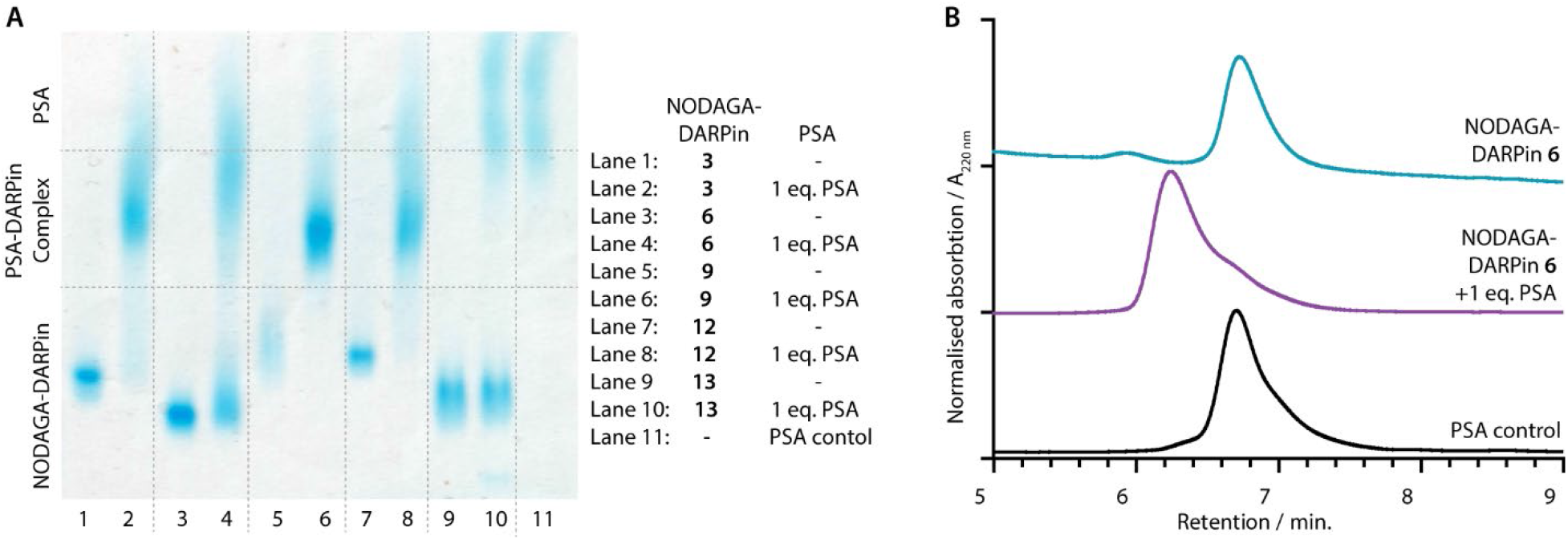
(A) Native PAGE analysis of PSA-binding: NODAGA-DARPins **3, 6, 9, 12** and **13** (lanes 1, 3, 5, 7 and 9, respectively) show bands in the lower third of the gel. NODAGA-DARPins **3, 6, 9**, and **12** incubated with 1 eq. PSA (lanes 2, 4, 6, and 8, respectively) form higher molecular weight NODAGA-DARPin-PSA complexes which appear in the centre of the gel with a broad band. The non-binding control NODAGA-DARPin **13** incubated with 1 eq. PSA (lane 10) does not form a complex and two distinct bands consistent with unbound proteins were detected. Note: Protein bands are stained with Coomassie blue, and PSA as a control is displayed in lane 11. (B) Analytical SEC analysis of PSA-binding: The NODAGA-DARPin **6** (cyan) was incubated with 1 eq. of PSA (black) giving a higher molecular weight complex with slightly shorter retention time (purple). Chromatographic profiles were measured by electronic absorption at 220 nm.

The same reaction mixtures were also analysed by SEC (**Figure 4**B and Supporting information **Figure S11**). As an example, NODAGA-DARPin **6** gave a major peak with a retention time, *R*_t_ of 6.66 min. (**Figure 4**B, cyan trace), which under the chromatography conditions, overlapped with the peak observed for the free PSA protein (**Figure 4**B, black trace, peak *R*_t_ = 6.67 min.). After incubating NODAGA-DARPin **6** with PSA, the peak observed in the SEC chromatogram (**Figure 4**B, purple trace) shifted to shorter retention time (*R*_t_ = 6.17 min.), which is characteristic of the formation of the higher molecular weight NODAGA-DARPin-PSA binary complex (total MW = ∼52 kDa). The SEC data also suggest that the NODAGA-DARPins bind to PSA with high affinity to produce complexes that are kinetically stable under the chromatographic conditions. For the non-binding control NODAGA-DARPin **13**, no shift in the *R*_t_ of the peaks was observed, which also provided support that the chromatographic shifts observed for NODAGA-DARPins (**3, 6, 9**, and **12**) reflect specific, high-affinity binding to PSA (Supporting information **Figure S11**).

### Measurement of PSA-binding affinities by surface plasmon resonance

The PSA-binding affinity for the various DARPin species was evaluated by using surface plasmon resonance (SPR) single-cycle kinetic measurements. SPR was used to estimate *k*_on_ (rate constant of association, M^-1^ s^-1^), *k*_off_ (rate constant of dissociation, s^-1^), and *K*_D_ (dissociation constant, M^-1^) of the DARPin-Cys and NODAGA-DARPins species with either PSA (**Table 1**), the PSA-α1-antichymotrypsin (PSA-ACT) complex, or ACT alone (Supporting information **Table S5, S6** and **Figure S12** – **S15**). The DARPin-Cys and NODAGA-DARPins were immobilised *via* their *N*-terminal His_6_ tags on a Ni-coated multidentate poly-nitrilotriacetic acid (NTA)-derivatized linear polycarboxylate chip surface (NiHC30M Chip). The analyte (PSA, PSA-ACT, or ACT) was passed through the chip at five different, increasing concentrations. All measurements were performed at 20 °C in HEPES buffer (HBS buffer). The non-binding DARPin-Cys **13** was immobilised in the reference flow cells 1 and 3. The use of DARPin-Cys **13** as a control stabilised the baseline and reduced non-specific binding of the analyte to the chip. When fitting the acquired data, we observed deviations from expected 1:1 binding kinetics. In most cases, modelling the experimental data with a sum of two exponentials (Sum2Exp) gave satisfactory fits. The human PSA analyte was obtained from natural sources and is known to contain at least 5 isoforms.^20,31,58^ Moreover, PSA carries a single *N*-linked glycosylation site at Asp69 that is known to be heterogeneous,^59^ as can also be seen in the diffuse band in **Figure 4**A. We suggest that the higher order kinetic profile observed in these measurements arises from these inhomogeneities of the analyte.

**Table 1.** Binding affinities measured in SPR experiments.^a^

| Probe | Analyte | Model | $k_{\text{on}} / 10^5 \text{ M}^{-1} \text{ s}^{-1}$ | $k_{\text{off}} / 10^{-3} \text{ s}^{-1}$ | $K_{\text{D}} / \text{nM}$ | $\text{RU}_{(\text{max})}$ | $\text{Chi}^2 / \text{RU}^2$ |
| --- | --- | --- | --- | --- | --- | --- | --- |
| DARPin-Cys **3** | PSA | Sum2Exp | $1.515 \pm 0.003$<br>$7.137 \pm 0.065$ | $2.440 \pm 0.001$<br>$12.92 \pm 0.005$ | $16.11 \pm 0.06$<br>$18.10 \pm 0.18$ | $117.1 \pm 0.3$<br>$55.92 \pm 0.37$ | 0.21 |
| NODAGA-DARPin **3** | PSA | Sum2Exp | $8.824 \pm 0.417$<br>$2.243 \pm 0.034$ | $0.8728 \pm 0.0073$<br>$4.046 \pm 0.043$ | $0.9891 \pm 0.009$<br>$18.04 \pm 0.33$ | $53.73 \pm 0.35$<br>$25.98 \pm 0.30$ | 0.72 |
| DARPin-Cys **6** | PSA | Sum2Exp | $8.646 \pm 0.038$<br>$1.201 \pm 0.003$ | $13.88 \pm 0.045$<br>$2.498 \pm 0.004$ | $16.06 \pm 0.09$<br>$20.79 \pm 0.06$ | $34.67 \pm 0.11$<br>$61.15 \pm 0.09$ | 0.066 |
| NODAGA-DARPin **6** | PSA | Sum2Exp | $5.399 \pm 0.027$<br>$1.432 \pm 0.041$ | $0.9626 \pm 0.0062$<br>$6.184 \pm 0.130$ | $1.783 \pm 0.015$<br>$43.18 \pm 1.54$ | $67.94 \pm 0.30$<br>$16.40 \pm 0.20$ | 0.62 |
| DARPin-Cys **9** | PSA | Sum2Exp | $1.104 \pm 0.006$<br>$8.924 \pm 0.632$ | $0.1460 \pm 0.0064$<br>$2.484 \pm 0.007$ | $1.322 \pm 0.059$<br>$2.783 \pm 0.021$ | $70.29 \pm 0.27$<br>$105.3 \pm 0.3$ | 0.44 |
| NODAGA-DARPin **9** | PSA | Sum2Exp | $46.95 \pm 0.39$<br>$2.428 \pm 0.010$ | $16.04 \pm 0.098$<br>$2.536 \pm 0.011$ | $3.416 \pm 0.036$<br>$10.44 \pm 0.06$ | $31.94 \pm 0.07$<br>$29.05 \pm 0.07$ | 0.295 |
| DARPin-Cys **12** | PSA | Sum2Exp | $9.484 \pm 0.063$<br>$0.8821 \pm 0.0043$ | $3.022 \pm 0.009$<br>$0.2977 \pm 0.0057$ | $3.186 \pm 0.023$<br>$3.375 \pm 0.067$ | $41.51 \pm 0.09$<br>$33.91 \pm 0.08$ | 0.069 |
| NODAGA-DARPin **12** | PSA | Sum2Exp | $16.05 \pm 0.08$<br>$1.683 \pm 0.006$ | $9.007 \pm 0.030$<br>$1.438 \pm 0.005$ | $5.613 \pm 0.039$<br>$8.543 \pm 0.045$ | $28.05 \pm 0.09$<br>$34.69 \pm 0.09$ | 0.21 |

<sup>a</sup> Summary of the results of single-cycle kinetic surface plasmon resonance (SPR) measurements of the DARPin-Cys and NODAGA-DARPins **3**, **6**, **9**, and **12** using PSA as analyte. (full data sets with PSA-ACT are given in Supporting Information **Tables S5** and **S6**). Measurements were performed on a NiHC30 chip at 20 °C in HBS buffer.

As summarised in **Table 1**, all DARPin-Cys and NODAGA-DARPins (**3, 6, 9**, and **12**) displayed comparable, nanomolar affinity for PSA and PSA-ACT. The measured *K*_D_ values ranged from 0.9 to 43 nM (full data are presented in **Tables S5** and **S6**; representative sensorgrams with fitted curves are presented in **Figure S12** – **S15**). The SPR data revealed that, in general, only minor differences were observed between the respective DARPin-Cys and NODAGA-DARPin pairs. In all cases, no significant differences were observed, and *K*_D_ values remained in the low nanomolar range, confirming that derivatisation did not impinge PSA binding.

SPR measurements were also performed using the PSA-ACT complex and ACT alone as the analytes (Supporting information **Table S5, S6** and **Figures S14**, and **S15**). No binding was observed for any of the DARPin constructs with the ACT protein (**Table S5** and **S6**). All DARPin-Cys proteins and two NODAGA-DARPins (**6** and **12**) displayed similar or higher affinity for the free PSA analyte *versus* the PSA-ACT complex, which suggest a small degree of selectivity for the catalytically active form of PSA. Conversely, for NODAGA-DARPins **3** and **9**, similar or slightly higher binding to the PSA-ACT complex were observed. However, the degree of selectivity displayed by our current pool of NODAGA-DAPRins (**3, 6, 9** and **12**) is likely insufficient to discriminate PSA from PSA-ACT using non-invasive imaging methods. Even though selective discrimination of free PSA and complexed (inactive) PSA-ACT could help stratify patient response to therapy, fPSA concentrations in cancerous prostate tissue are several orders of magnitude higher than total PSA levels found in circulation.^34^ This should lead to a sufficiently high contrast, even without selectivity for fPSA.

### Radiosynthesis of [^68^Ga]GaNODAGA-DARPins

Gallium-68 (half-life, *t*_1/2_ = 67.7 min.; positron emission intensity, *I*(β^+^) = 88.9%)^60^ radiolabelling of the NODAGA-DARPins was achieved by using standard radiolabelling conditions (**Figure 5**A).^61^ The positron-emitting radionuclide ^68^Ga was selected because the short half-life of this radionuclide provides a complementary match with the rapid tissue uptake and excretion profiles that have been reported for other radiolabelled DARPins *in vivo*.^54,56,62^ Briefly, an aliquot of [^68^Ga][Ga(H_2_O)_6_]Cl_3_ (3 – 4 MBq) was added to a buffered solution of NODAGA-DARPin (∼0.40 nmol) in NaOAc (0.1 M, pH 4.4), and the pH was adjusted to 5.3 – 5.5 by the addition of Na_2_CO_3_ (1 M). Successful radiometallation gave the [^68^Ga]GaNODAGA-DARPin species in <10 min. at room temperature. The [^68^Ga]GaNODAGA-DARPin radiotracers were obtained with high radiochemical purities (RCP) >99%, and molar activities (*A*_m_) ranging from 30.7 to 54.4 MBq nmol^-1^ of protein (Supporting Information **Figure S16** and **Table S7**).

**Figure 5.**
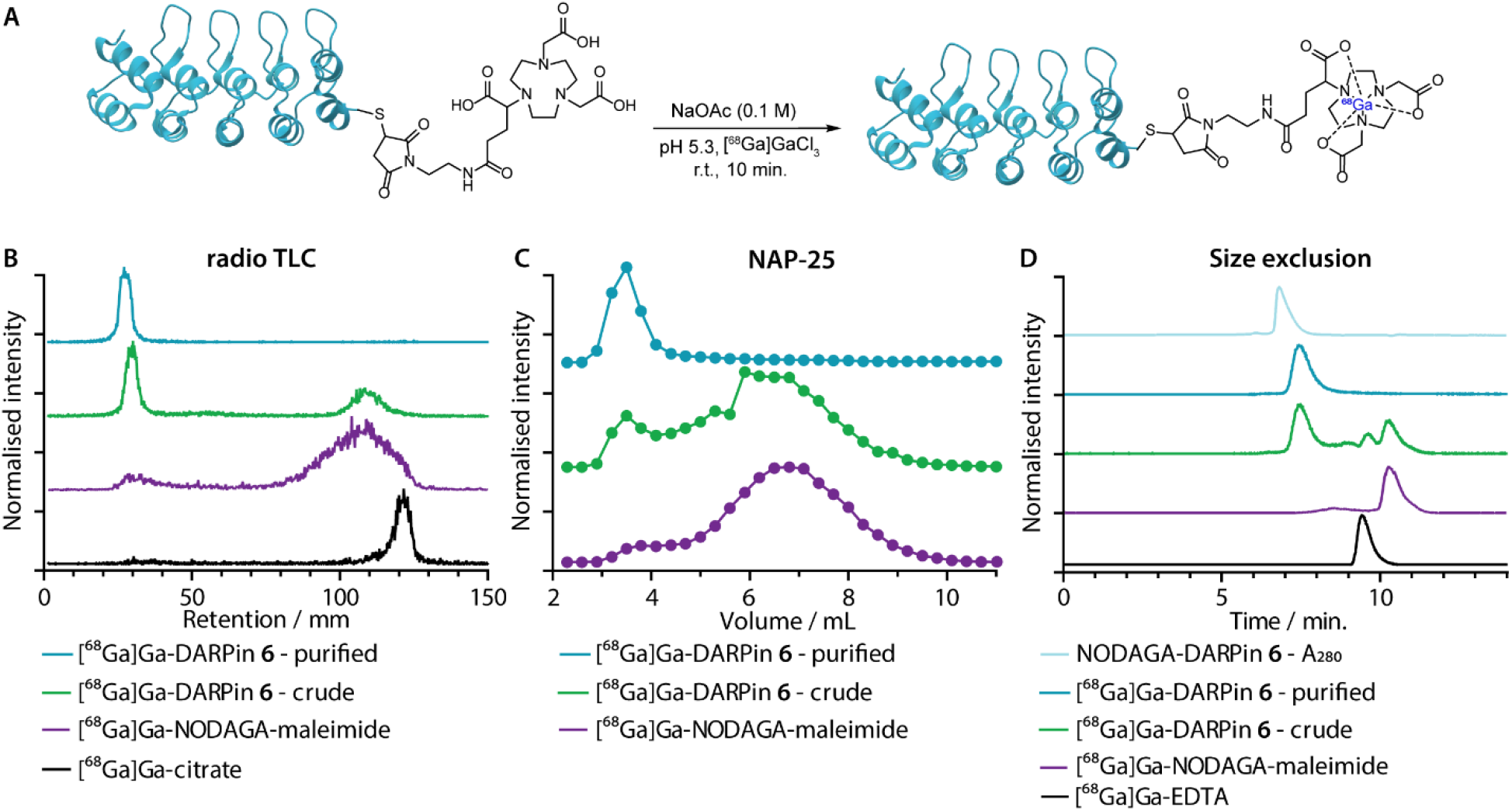
(A) Reaction scheme showing the one-step ^68^Ga-radiolabelling of the NODAGA-DARPins. (B) Representative radio-thin layer chromatography (radio-TLC; developed with 0.2 M, pH 4.4 aqueous sodium citrate as the mobile phase) data showing characterisation of the purified (cyan) and crude (green) samples of [^68^Ga]GaNODAGA-DARPin **6**. The purple and black traces show control data obtained with [^68^Ga]GaNODAGA-maleimide and [^68^Ga]Ga-citrate, respectively. (C) Manual SEC analysis of the purified (cyan) and the crude (green) construct as well as [^68^Ga]GaNODAGA-maleimide (purple) obtained by using NAP-25 columns eluted with PBS; (D) Automated SEC analysis showing a co-elution of the NODAGA-DARPin **6** UV trace (light blue) with the radiotrace of purified (cyan) and crude (green) samples of [^68^Ga]GaNODAGA-DARPin **6**. Chromatographic profiles are also presented for the control compounds [^68^Ga]GaNODAGA-maleimide (purple), and [^68^Ga]Ga-EDTA (black), both of which show distinct retention times.

Radiometal ion complexation of the non-purified crude reaction mixtures as well as the NAP-25 purified NODAGA-DARPins were analysed by using three separate chromatographic methods including radioactive instant thin-layer chromatography (radio-TLC), manual NAP-25 gel chromatography, and automated SEC coupled to an electronic absorption detector and a radioactivity flow detector in series. Representative data for ^68^Ga-radiolabelling of NODAGA-DARPin **6** are given in **Figures 5**B, C, and D, respectively. Full radiochemical data for [^68^Ga]GaNODAGA-DARPin radiotracers (**3, 6, 9, 12** and control **13**) are presented in Supporting Information **Figure S16**.

As an example, radio-TLC analysis (sodium citrate eluent, 0.2 M, pH 4.4) showed retention of the activity at the baseline (*R*_f_ = 0.0 – 0.1) for both the crude and the purified samples of [^68^Ga]GaNODAGA-DARPin-**6** (**Figure 6B**, green and cyan traces, respectively), indicative of radiolabelled protein. In the case of the crude sample, 35 – 45 % of the activity eluted at the solvent front and is assigned to the presence of an excess of [^68^Ga]GaNODAGA-maleimide. After the crude NODAGA-DARPin samples were purified, quantitative ^68^Ga-radiolabelling gave clean radio-TLC profiles whereby all the activity was retained at the baseline. For comparison, the elution profiles of the control samples of [^68^Ga]Ga-citrate (*R*_f_ = 0.95 – 1, **Figure 5**B, black trace) and [^68^Ga]GaNODAGA-maleimide (*R*_f_ = 0.85 – 1.0, **Figure 5**B, purple trace) are also presented, confirming that the presence of a major peak at the baseline corresponds to the ^68^Ga-labelled protein and that the small-molecule component present in the ^68^Ga-radiolabelling of the crude samples of NODAGA-DARPin **6** corresponds to [^68^Ga]GaNODAGA-maleimide.

**Figure 6.**
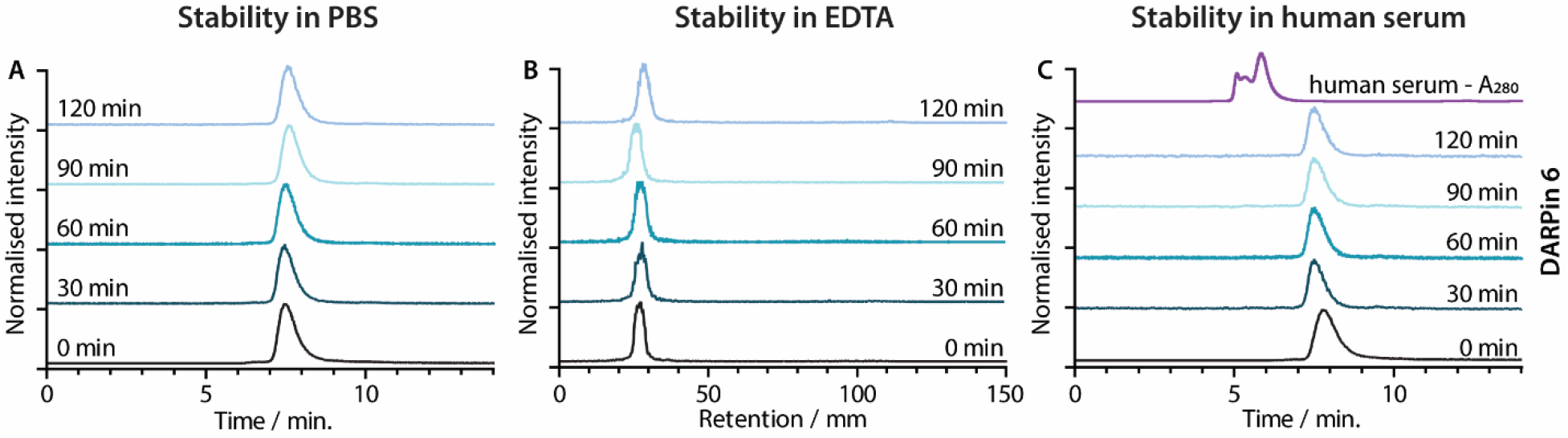
Stability of [^68^Ga]GaNODAGA-DARPin **6** in (A) the formulation buffer (PBS) at 37 °C assessed by radio-SEC, (B) excess EDTA at room temperature assessed by radio-*i*TLC and (C) human serum at 37 °C assessed by radio-SEC. The black traces show the baseline data obtained at *t* = 0 min. The fading blue traces indicate the analysis of aliquots taken at 30 min intervals from *t* = 30 – 120 min. The UV trace of human serum is shown in panel C in purple.

Analytical size-exclusion chromatography performed by using NAP-25 columns showed that the purified sample of [^68^Ga]GaNODAGA-DARPin-**6** (**Figure 5**C, cyan trace) gave a single peak with an elution maximum at ∼3.5 mL. These data also confirmed that the manual size-exclusion purification process successfully removed all small-molecule components from the NODAGA-DARPin conjugation step, as well as any non-specifically bound ^68^Ga in the radiolabelling step. Similarly, automated SEC chromatograms of the crude (**Figure 5**D, green trace) and purified samples of [^68^Ga]GaNODAGA-DARPin **6** (**Figure 5**D, cyan trace) indicated that the radiotracer was obtained in >99% RCP, where the peak for the product co-eluted with the electronic absorption chromatogram of NODAGA-DARPin **6** (**Figure 5**D, light blue trace, measured at 280 nm). Similarly, SEC chromatograms showed the absence of any non-specifically bound ‘free’ ^68^Ga-ions (represented by [^68^Ga]Ga-EDTA, (peak *R*_t_ = 9.3 min., **Figure 5**D, black trace) or non-conjugated [^68^Ga]GaNODAGA-maleimide reagent (peak *R*_t_ = 10.2 min., **Figure 5**D, purple trace). Overall, the radiochemical data confirmed that all the ^68^Ga-labelled NODAGA-DARPins could be synthesised in sufficient radiochemical yield, chemical and radiochemical purity, and with molar activities between 30.72 to 54.37 MBq nmol^-1^ (Supporting Information **Table S7**) suitable for applications in further biochemical studies.

### Radiotracer stability studies

Concerns have been raised over the stability of maleimide linkages *in vivo*,^63^ and a range of alternative bioconjugation methods have been developed.^62,64–66^ To evaluate if the stability profiles of the [^68^Ga]GaNODAGA-DARPin radiotracers were suitable for *in vivo* experiments, their radiochemical stability was measured *in vitro* over time in formulation buffer (PBS, pH 7.4), in chelate challenge studies using excess ethylenediamine tetraacetic acid (EDTA, 3700 eq., pH 7.1), and in human serum (37 °C, pH 7.4). Representative data for [^68^Ga]GaNODAGA-DARPin **6** are presented in **Figure 6**, and additional data for the other radiotracers are given in Supporting Information **Figure S17** and **Table S8** – **S10**. All the [^68^Ga]GaNODAGA-DARPin radiotracers were found to be stable in PBS with no change in radiochemical activity over 2 h (radio-SEC data, **Figure 5**A and **S17** and **Table S8**). Chelate challenge experiments using a large excess of EDTA also indicated that the [^68^Ga]GaNODAGA-DARPins complex remained intact with <9% transchelation observed after 2 h (radio-TLC data, **Figure 5**B and **S17**, and **Table S9**). Finally, human serum challenge experiments revealed that the [^68^Ga]GaNODAGA-DARPins did not bind to serum proteins and were also stable with respect to loss of the ^68^Ga metal ion, or the [^68^Ga]GaNODAGA chelate whereby measurements of the RCP over time showed that it remained >96% pure after 2 h (radio-SEC data, **Figure 5**C and **S17** and **Table S10**). Overall, the stability studies indicated that neither the metal ion is released from NODAGA, nor the [^68^Ga]Ga-NODAGA is cleaved from the proteins. Thus, our [^68^Ga]GaNODAGA-DARPins radiotracers were suitable for further evaluation in animal models.

### Radiotracer PSA-binding *in vitro*

To confirm that our ^68^Ga-radiolabelling conditions did not impair the specific binding of the [^68^Ga]GaNODAGA-DARPin radiotracers to PSA, we developed a radioactive PSA-binding assay by using streptavidin-coated magnetic beads. An illustration of the binding assay is presented in **Figure 7**A, and a summary of the data obtained for [^68^Ga]GaNODAGA-DARPin radiotracers (**3, 6, 9, 12**, and control **13**) are shown in **Figure 7**B, (with further details given in Supporting Information **Table S11**). Briefly, the [^68^Ga]GaNODAGA-DARPin radiotracers were first incubated with biotinylated PSA (10 eq.) to form the [^68^Ga]GaNODAGA-DARPin-PSA-biotin binary complex. Then, the mixture was incubated with streptavidin-coated magnetic beads to pull down the [^68^Ga]GaNODAGA-DARPin-PSA-biotin complex. After magnetic separation, the activity associated with the beads and with the supernatant was measured, and data were normalised with respect to the total activity loaded to determine the bound fraction of [^68^Ga]GaNODAGA-DARPins. As a control, blocking studies were performed whereby excess DARPin-Cys (250 eq. **Figure 7**B, block) was used to saturate the available PSA-biotin. In separate controls no PSA-biotin was added to the reaction mixture (**Figure 7**B, no PSA) to allow the measurement of non-specific binding to the magnetic beads. These binding assays confirmed that the [^68^Ga]GaNODAGA-DARPin radiotracers remained biochemically active with specific binding to PSA of between 37% – 48% of the total administered activity (*n* = 3 independent measurements per sample). In comparison, the non-binding control [^68^Ga]GaNODAGA-DARPin **13** showed only 4.8 ± 1.0% non-specific binding to the streptavidin magnetic beads (**Figure 7**B, red and Supporting Information **Table S11**). These data are consistent with the prior non-radioactive binding assays (*vide supra*) and indicate that the ^68^Ga-radiolabelling reactions did not compromise the biochemical viability or PSA-specificity of the [^68^Ga]GaNODAGA-DARPin constructs.

**Figure 7.**
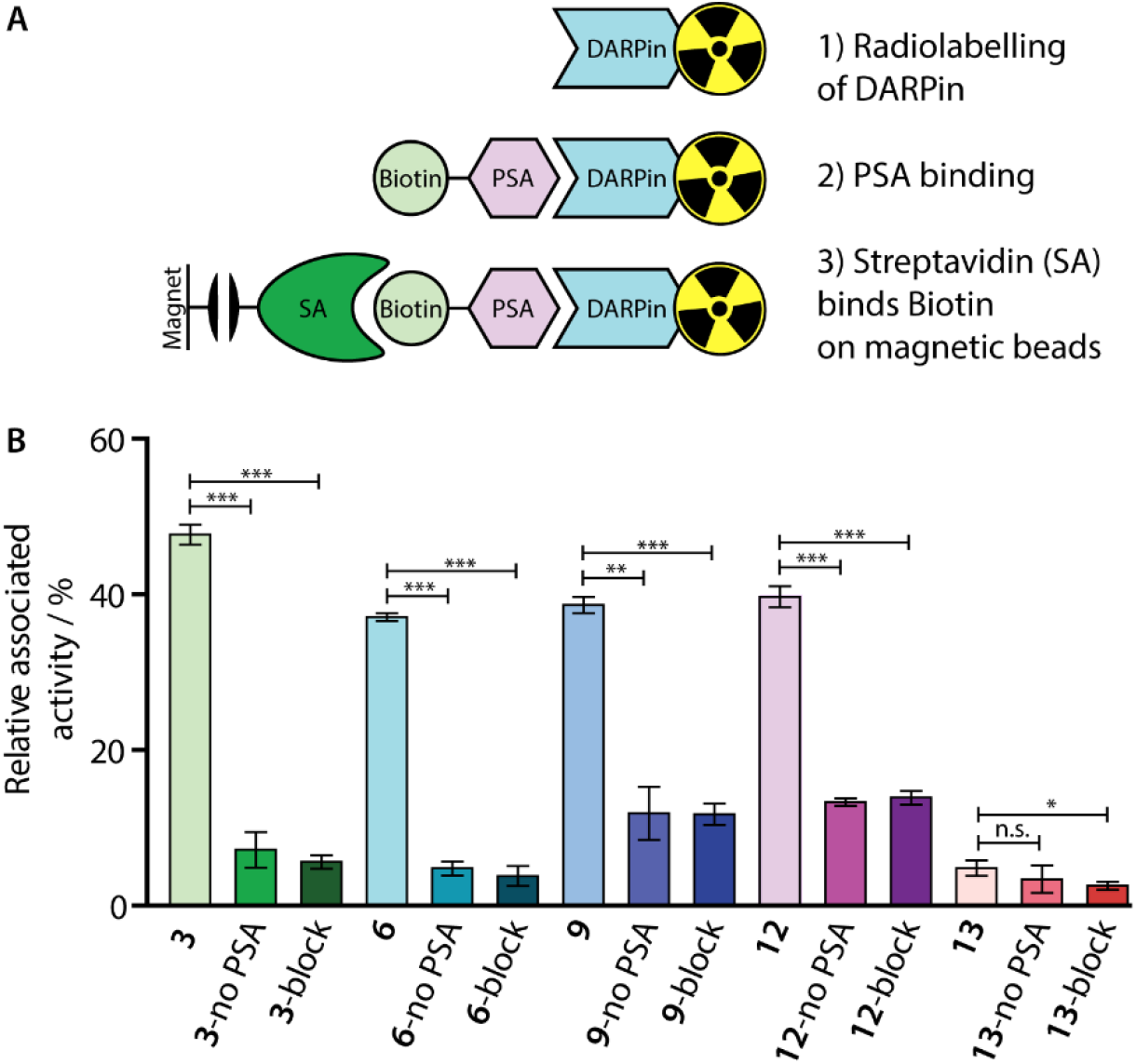
(A) Schematic representation of the *in vitro* PSA-binding assay developed to assess the biochemical viability of the [^68^Ga]GaNODAGA-DARPin radiotracers. (B) Bar chart showing the relative activities associated with the magnetic beads for the [^68^Ga]GaNODAGA-DARPin radiotracers with and without the addition of biotinylated PSA (no PSA), and in blocking studies containing excess non-radiolabelled DARPins (block).

### Pharmacokinetic studies *in vivo*

After demonstrating their high stability and biochemical specificity toward PSA-binding, the pharmacokinetic profiles and tumour specificity of the [^68^Ga]GaNODAGA-DARPins were evaluated *in vivo*. Athymic nude mice bearing subcutaneous LNCaP (PSA *+ve*) xenografts on the right flank (151.3 ± 78.5 mm^3^; *n* = 35) were randomised before the study with *n* = 3 or 4 animals per group. The [^68^Ga]GaNODAGA-DARPins radiotracers (**3, 6, 9, 12**, and control **13**) were purified manually by using size-exclusion NAP-5 gel chromatography to provide the purified radiotracers with decay-corrected radiochemical yields (d.c. RCYs) between 57 and 78% (*n* = 2, Supporting Information **Table S12**) in the formulation buffer (PBS, pH 7.4). [^68^Ga]GaNODAGA-DARPins **3, 6, 9, 12**, and **13** were administered by intravenous tail-vein injections (150 µL, 0.20 – 0.21 nmol of protein, 1.56 – 2.88 MBq per injection; see Supporting Information **Table S12**). In addition to the use of the non-binding control DARPin **13**, competitive (blocking) inhibition studies were performed to assess the tumour specificity of the PSA-binding radiotracers. For these blocking studies, doses of the [^68^Ga]GaNODAGA-DARPins radiotracers (**3, 6, 9**, and **12**) were prepared with the same amount of activity, but a reduced molar activity by co-administration of a 100-fold excess of the equivalent non-radiolabelled DARPin-Cys.

After radiotracer administration, the whole-body effective half-lives (*t*_1/2_(eff) / min.) of the different [^68^Ga]GaNODAGA-DARPin radiotracers were measured in each mouse for up to 2 h by using a dose calibrator (**Figure 8** and Supporting Information **Table S13**). Data indicated that the smallest radiotracer, [^68^Ga]GaNODAGA-DARPin **9**, had a *t*_1/2_(eff) value of 52.5 ± 3.6 min. The intermediate sized constructs, [^68^Ga]GaNODAGA-DARPins **3** and **12**, gave *t*_1/2_(eff) values of 61.3 ± 12.8 min. and 40.0 ± 5.6 min., respectively. The two largest constructs, [^68^Ga]GaNODAGA-DARPin **6** and the non-binding control **13** gave measured *t*_1/2_(eff) values of 39.2 ± 4.8 min. and 112.6 ± 32.3 min., respectively. In addition, changing the administered protein dose in the blocking groups also gave only small changes in the measured *t*_1/2_(eff) values. For example, *t*_1/2_(eff) values of [^68^Ga]GaNODAGA-DARPin **3**-block and **9**-block decreased (55.9 ± 22.0 min., and 38.67 ± 7.4 min., respectively), but measurements showed a slight increase for **6**-block (52.0 ± 17.0 min.), and **12**-block (64.6 ± 20.2 min.). In all cases, between 70% – 80% of the administered activity was eliminated from the mice by 2 h post-administration. These observations suggest that the elimination rates are all rather similar, and that other factors beyond radiotracer size and administered protein dose are likely to be involved in small differences in the tissue distribution and pharmacokinetic profiles of these [^68^Ga]GaNODAGA-DARPin radiotracers. Overall, these measurements of whole-body pharmacokinetics are consistent with previous published data which showed rapid clearance of DARPin-based radiotracers from the circulation and elimination *via* the renal pathway, indicating that proteins whose size is smaller than the glomerular filtration unit are not following a size-dependent elimination rate.^54,55,62^ Provided that the radiotracers also display rapid tumour uptake and specific retention of activity, the fast excretion *via* renal clearance is a desirable property for most molecular imaging agents, which helps to reduce background activity in healthy tissue, and increase target-to-background contrast.

**Figure 8.**
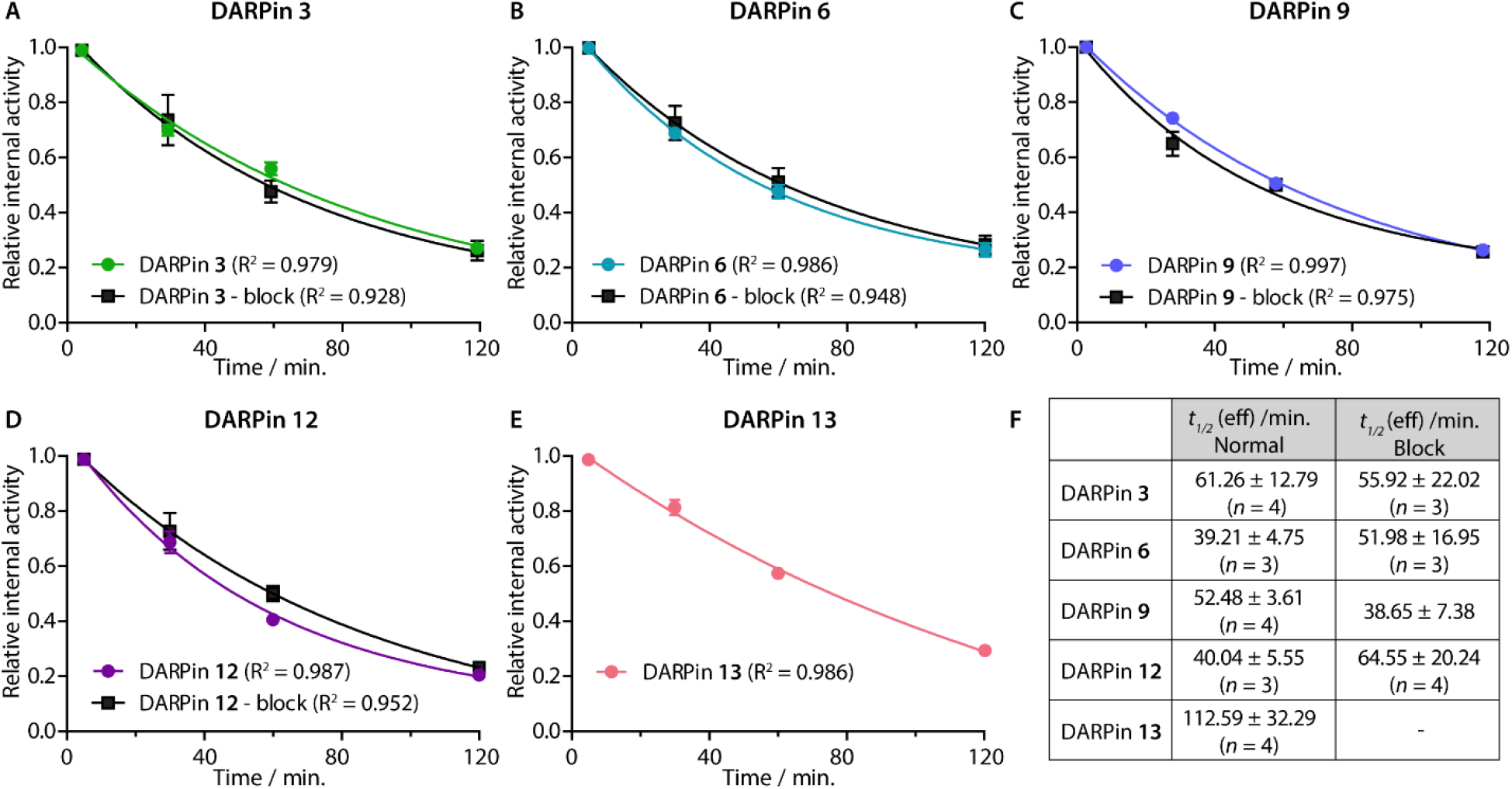
(Panels A – E) Relative internal activities plotted against time to determine *t*_1/2_ (eff) of the five [^68^Ga]GaNODAGA-DARPins in the normal and blocking groups of in athymic nude mice bearing *s*.*c*. LNCaP xenografts. (F) Table summarising the calculated *t*_1/2_ (eff) values obtained from a one-phase decay model. Note: Data were analysed in GraphPad Prism.

### Biodistribution studies

*Ex vivo* analysis of the tissue distribution of [^68^Ga]GaNODAGA-DARPin (**3, 6, 9, 12**, and control **13**) was performed at 2 h post-administration (**Figure 9**, and Supporting Information **Table S14)**. Tumour-associated activity varied between the four PSA-binding [^68^Ga]GaNODAGA-DARPins (**3, 6, 9**, and **12**). The smallest and largest radiotracers, [^68^Ga]GaNODAGA-DARPin **9** and [^68^Ga]GaNODAGA-DARPin **6**, showed the highest tumour uptake with values of 4.72 ± 1.07 %ID g^-1^ and 4.16 ± 0.58 %ID g^-1^, respectively. For both radiotracers, significantly higher tumour uptake was observed compared to the non-binding control radiotracer [^68^Ga]GaNODAGA-DARPin **13** (2.93 ± 0.55 %ID g^-1^, Student’s *t*-test *P*-value <0.05). The intermediate-sized radiotracers ([^68^Ga]GaNODAGA-DARPin **3** and **12**), which both feature two ankyrin repeat units, were found to have lower tumour uptake and retention (2.39 ± 0.43 %ID g^-1^ and 1.80 ± 0.59 %ID g^-1^, respectively). Blocking studies were used to measure the degree of PSA-specific binding in tumours *in vivo*. In the blocking studies, both [^68^Ga]GaNODAGA-DARPin **6** and **9** showed a significant decrease of ∼50% in tumour-associated uptake (2.47 ± 0.42 %ID g^-1^ and 2.90 ± 0.46 %ID g^-1^, respectively [both *P*-values <0.05 for the normal *versus* blocking groups]). [^68^Ga]GaNODAGA-DARPin **3** also showed a specific tumour-associated uptake where tumour-associated uptake in the blocking group decreased by ∼ 39% to 1.48 ± 0.19 %ID/g (*P*-value <0.05). In contrast, [^68^Ga]GaNODAGA-DARPin-**12** gave the lowest level of tumour-associated uptake which was found to be statistically the same as uptake observed in the blocking group (1.58 ± 0.28 %ID/g).

**Figure 9.**
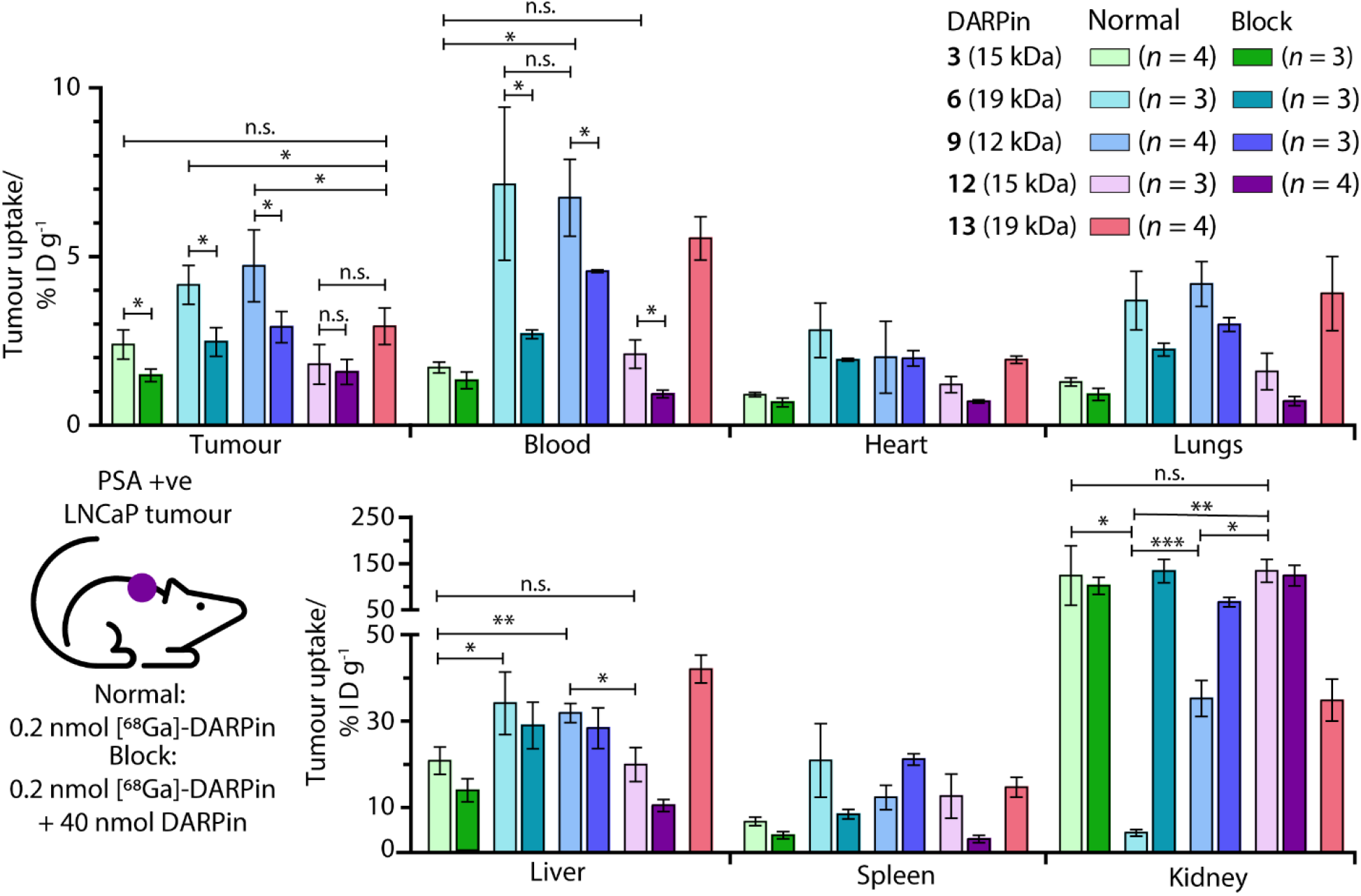
Bar charts showing the e*x vivo* biodistribution analysis in selected organs for the five [^68^Ga]GaNODAGA-DARPin radiotracers in mice bearing subcutaneous LNCaP (PSA *+ve*) tumours. Statistically significant changes are displayed with \**P* <0.05; \*\**P* <0.01. \*\*\**P* <0.001 and n.s. (not specific) >0.05

Unlike humans, mice do not carry functional PSA (KLK3) genes, and thus do not express PSA.^67,68^ Moreover, in humans, PSA concentrations at prostate tissue have been reported to be 10^6^-fold higher in comparison to concentrations in the blood serum.^34^ Therefore, background tissue uptake in organs due to PSA-related binding should be minimal in the reported experiments. We evaluated the radioactivity accumulation in background organs. Smaller proteins, like DARPins are known to be excreted *via* renal clearance. As expected, [^68^Ga]GaNODAGA-DARPin **3** and **12**, both ∼15 kDa in size, showed very similar organ distribution. One notable difference was found in the accumulation of the activity in bone, for which the uptake of [^68^Ga]GaNODAGA-DARPin **3** was significantly lower at 1.26 ± 0.17 %ID g^-1^ than for **12** (2.35 ± 0.39 %ID g^-1^, *P*-value <0.05). For [^68^Ga]GaNODAGA-DARPin **3** and **12**, retention of activity in the kidneys was high (139.3 ± 65.6 %ID g^-1^ and 149.6 ± 25.0 %ID g^-1^, respectively). In contrast, both the larger and the smaller [^68^Ga]GaNODAGA-DARPin radiotracers (**6** and **9**) showed considerably lower accumulation and retention of activity in the kidney (4.44 ± 0.68 %ID g^-1^ and 36.13 ± 4.28%ID g^-1^, respectively). The lowest kidney uptake, and the highest tumour-to-kidney contrast ratio (0.94 ± 0.19, **Table S15**) was observed for [^68^Ga]GaNODAGA-DARPin **6**. Interestingly, an inverse trend was observed for liver accumulation whereby [^68^Ga]GaNODAGA-DARPin **3** (21.31 ± 3.24 %ID g^-1^) and **12** (20.41 ± 3.99 %ID g^-1^), had lower liver retention than **6** (33.16 ± 7.93 %ID g^-1^), and **9** (32.65 ± 2.31 %ID g^-1^).

Blood pool activity was found to be considerably higher for [^68^Ga]GaNODAGA-DARPin **6** (7.13 ± 1.25 %ID g^-1^) and **9** (6.74 ± 1.14 %ID g^-1^) than either **3** (1.71 ± 0.16 %ID g^-1^), **12** (2.10 ± 0.42 %ID g^-1^), or the non-binding control **13** (5.53 ± 0.64 %ID g^-1^). Decreased clearance through the kidneys appears to correlate with prolonged circulation in the blood pool, and a concordant increase in clearance *via* the hepatobiliary pathway, contrasting DARPin **6** and **9** on the one hand with DARPins **3** and **12** on the other. All tumour-to-tissue ratios and calculated *P*-values are given in Supporting Information **Table S15** – **S18**.

From *in vitro* data alone, it was difficult to differentiate between the four candidate PSA-binding radiotracers. *In vivo* we could observe major differences in tumour-specific uptake and tissue distribution which might arise form sequence-specific differences between the four evaluated DARPins (Supporting Information **Figure S2** and **S3**). Prolonged circulation in the blood pool, specific tumour uptake with high contrast over most background tissues, and low levels of accumulation in the kidneys and bone, are attractive features when developing diagnostic radiotracers. In this regard, the *ex vivo* biodistribution analysis indicated that [^68^Ga]GaNODAGA-DARPin **6** and **9** are the most promising candidates for measuring specific changes in PSA-expression *in vivo*. Beyond the use of these radiotracers for non-invasive biomarker detection and quantification, these DARPin-based radiopharmaceuticals could be attractive compounds for developing molecularly targeted radiotherapeutics, provided the PSA-specific tumour-associated uptake can be further increased by a factor of ∼2 – 3 (as previously shown).^41,42^ Such DARPins could be attractive compounds for developing molecularly targeted radiotherapeutics using β– emitting radionuclides such as ^90^Y, ^177^Lu, or ^188^Re, or α-particle (or decay-chain) emitters such as ^211^At, ^212^Pb, or ^225^Ac. Future work will focus on fine-tuning the pharmacokinetic profile of DARPin-based radiotracers to facilitate diagnostic imaging and molecularly targeted radiotherapy.

## Conclusion

Here, we report the successful selection and evaluation of new PSA-targeting DARPins for use in the quantification of biomarker expression profiles *in vivo*. Four PSA-binding DARPins were selected from a pool of 380 hits. The DARPins were conjugated site-specifically to a metal binding chelate *via* a unique *C*-terminal Cys residue. *In vitro* studies, combined with radiolabelling experiments, demonstrated that all four chosen PSA-binding DARPins showed low nanomolar affinity and specific targeted binding toward PSA. Additional radiochemical studies *in vitro* confirmed the stability, the PSA-binding specificity, and the suitability of the ^68^Ga-radiolabelled probes for use in biomarker quantification. *In vivo* experiments using tumour-bearing mice confirmed that three radiotracers showed significant and specific tumour uptake. [^68^Ga]GaNODAGA-DARPin radiotracers **6** and **9** performed the best, showing promising tumour uptake with high contrast over background tissue, and desirable excretion profiles with low retention of the activity in the kidneys. Collectively, these results support the conclusion that new DARPin-based radiotracers can be developed for applications in non-invasive molecular imaging and radiotherapy of PCa *via* targeting the local tissue expression of PSA.

## Supporting information

ElectronicSupplementaryInformation

## Author contributions

M.G. performed most of the experiments including the synthesis, radiochemistry, *in vitro* and *in vivo* testing. B.D. performed the SEC-MALS analysis and coordinated DARPin screening and selection assays which were performed by S.F. and supported by co-workers mentioned in the acknowledgement. J.S. conducted the SPR measurements. J.P.H and A.P. supervised the project. M.G. wrote the first draft of the manuscript which was edited by B.D., A.P. and J.P.H. All authors read and approved the final version of the manuscript.

## Conflicts of interest

There are no conflicts to declare.

## Data availability statement

The data that support the findings of this study are available from the corresponding author upon reasonable request.

## Additional information

Supplementary information (PDF) is available and contains experimental details, high-resolution mass spectrometry data and HPLC chromatograms for all relevant compounds, as well as additional data from the radiochemistry and *in vivo* experiments.

## Acknowledgements

JPH is supported by the Swiss National Science Foundation (SNSF Professorship PP00P2_163683 and PP00P2_190093), the Swiss Cancer Research Foundation (KFS-4257-08-2017), and the University of Zurich (UZH) for financial support. We thank all members of the Holland group and Plückthun group for helpful discussions and continuous support. In particular, we thank Dr Rachael Fay and Dr Jonas V. Schaefer for their initial work on DARPin radiotracers, Dr José Esteban Flores for his contribution to the *in vivo* work, and Dr Sylvie Briand-Schumacher, Thomas Reinberg, Joana Marinho, Gabriela Nagy, and Christa Booy-Rutgers for contributions in the ribosome display, selection, and purification.

## Table of contents graphic

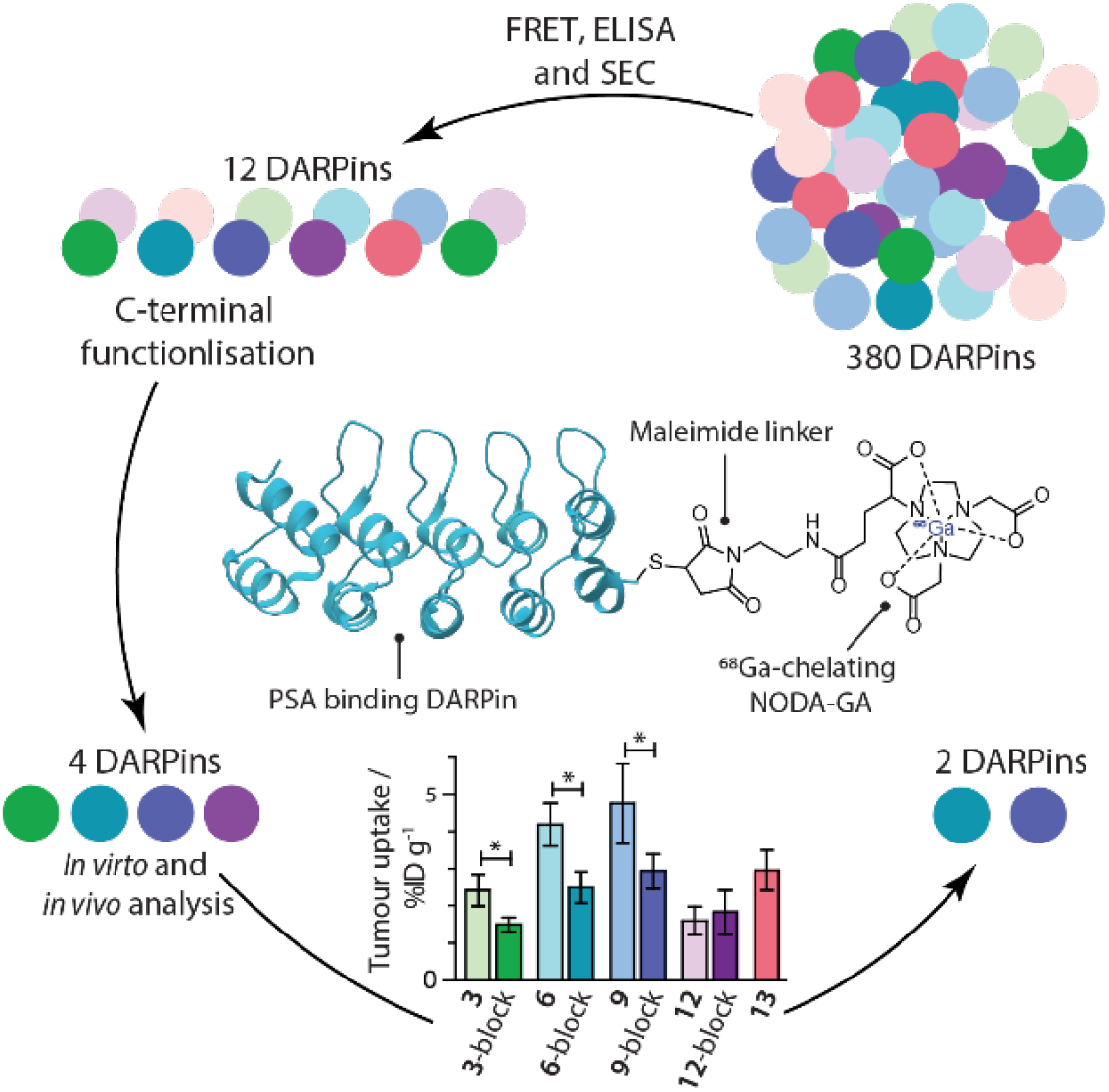

## Notes

### Competing Interest Statement

The authors have declared no competing interest.

