## Supplementary material for "Designed Ankyrin Repeat Proteins for detecting prostate-specific antigen expression *in vivo*": ElectronicSupplementaryInformation

**Electronic Supporting Information (ESI)**

*** Corresponding Author:**

Prof. Dr Jason P. Holland

**First author:**

Melanie Gut

#### General information

##### Chemicals

Unless otherwise stated, all other chemicals were of reagent grade and purchased from SigmaAldrich (St. Louis, MO), Merck (Darmstadt, Germany), Tokyo Chemical Industry (Eschborn, Germany), abcr (Karlsruhe, Germany) or CheMatech (Dijon, France). Water (>18.2 MΩ·cm at 25 ºC, Merck Millipore Milli-Q® Reference A+, with an additional 0.22 µm Millipack Express40 filter) was used without further purification for chemical and biological experiments, and additionally purified by aid of a Chelex® resin (100 sodium form, 100 – 200 mesh particle size, Sigma Aldrich) for radiochemical experiments.

##### Mass spectrometry

Liquid chromatography high resolution electrospray ionisation mass spectra (LC-HR-ESI-MS) were measured by the mass spectrometry service at the Department of Chemistry, University of Zurich.

Acquity UPLC (Waters, Milford, USA) connected to an Acquity eλ diode array detector and a Synapt G2 HR-ESI-QTOF-MS (Waters, Milford, USA); injection of 1 – 5 µL sample (concentration ca. 10-100 µg $mL^{-1}$ in the indicated solvent); UV spectra were recorded from 200 – 600 nm at 1.2 nm resolution and at a sampling rate of 20 points $s^{-1}$; ESI: positive ionisation mode, capillary voltage 3.0 kV, sampling cone 40 V, extraction cone 4 V, $N_2$ cone gas 4 L $h^{-1}$, $N_2$ desolvation gas 800 L $min^{-1}$, source temperature 120 °C; mass analyser in resolution mode: mass range 100 – 2,000 *m/z* with a scan rate of 1 Hz; mass calibration to <2 ppm within 50–2500 *m/z* with a 5 mM aqueous solution of $NaHCO_2$, lock-masses: *m/z* 195.0882 (caffein, 0.7 ng $mL^{-1}$) and 556.2771 (leucine-enkephalin, 2 ng $mL^{-1}$).

Acquity BEH C18 HPLC column (1.7 µm particle size, 2.1 × 50 mm, Waters) kept at 30 °C; elution at a flow rate of 400 µL $min^{-1}$ with solvent A: $H_2O$ + 0.02% TFA and solvent B: $CH_3CN$ + 0.02% TFA, linear gradient from 10 – 95% B within 3 min., then isocratic for 2 min. at 95% B.

Acquity BEH C8 HPLC column (1.7 µm particle size, 2.1 × 100 mm, Waters) kept at 30 °C; elution at a flow rate of 400 µL $min^{-1}$ with solvent A: $H_2O$ + 0.02% TFA and solvent B: $CH_3CN$ + 0.02% TFA, linear gradient from 10 – 70% B within 10 min.

##### Enzyme-linked immunosorbent assay (ELISA)

Biotinylated PSA was immobilised as follows: 384-well high binding microplates (MTP384 Greiner Bio-One GmbH, Frickenhausen, Germany) were coated with 20 µL of streptavidin (66 nM in PBS) overnight at 4 °C and blocked with 0.2% BSA (*w/v*) (45 µL PBS, supplemented with 0.1 [v/v] Tween20 and 0.2% [w/v] BSA [PBS-TB]) for 1 h at 4 °C with orbital shaking (Titramax 1000, at 900 rpm). After washing, biotinylated PSA (20 µL of 50 nM in PBS-TB supplemented with 1 mM DTT) was immobilised for 2 h, 4 °C, 900 rpm. IMAC-purified DARPin constructs (expressed and selected as described below; 20 µL, 100 nM in PBS supplemented with 0.1 [v/v] Tween20 [PBS-T] and 1 mM DTT) were applied to wells with or without immobilised PSA for 1.5 h, 4 °C, 900 rpm. After extensive washing, binding was detected by incubation with anti-FLAG antibody (FLAG-tagged DARPins; 20 µL, 1:5'000 diluted in PBS-TB; mouse anti-FLAG IgG, F3165, Sigma-Aldrich, Buchs, Switzerland) or anti-RGS-His antibody (DARPin-Cys; 20 µL, 1:5'000 diluted in PBS-TB; mouse anti RGA $(His)_4$ IgG1, No.

34650, Qiagen, GmbH, Hilden, Germany, which specifically discriminates between the *N*-terminal RGS-His6 tags of the DARPins and the *N*-terminal $His_6$ tag of the antigen) for 1 h, 4 °C, 900 rpm. After washing, anti-mouse IgG-alkaline phosphatase conjugate (20 µL, 1:10,000 diluted in PBS-TB; Pierce, Thermo Fisher Scientific) was applied to wells and the plate was incubated for 1 h, 4 °C, 900 rpm. After extensive washing *p*-nitrophenylphosphate (20 µL of 50 µM solution containing 1 M $MgCl_2$ and 1 M $NaHCO_3$; Sigma-Aldrich, Buchs, Switzerland) was added and the plate was incubated for 30 min., 37 °C, 900 rpm, and the $OD_{405\text{-}540nm}$ was measured (Micro Plate Reader, BIO-TEK Synergy). All washing steps were performed with 110 µL PBS-T per well on an ELx405 microplate washer (BioTek, Winooski, USA). Dispensing steps were performed on a MicroFlo Select liquid dispenser (BioTek). DARPin constructs were applied with a Liquidator96 system (Mettler-Toledo GmbH, Greifensee, Switzerland).

**Size-exclusion chromatography**

Automated size-exclusion chromatography was performed using an Agilent AdvanceBio SEC 130 Å column (Agilent Technologies, AdvanceBio SEC 130 Å, 2.7 µm 4.6 × 300 mm) connected to a Hitachi Chromaster Ultra Rs system equipped with a UV/visible detector (electronic absorption measured at 220 and 280 nm) as well as a radioactivity detector (FlowStar$^2$ LB 514, Berthold Technologies, Zug, Switzerland). Note that the UV/Vis detector and radioactivity detector were arranged serially with an offset time of approximately 0.10 min. The identity of the radiolabelled $^{68}$Ga-compound was confirmed by co-injection with an authenticated sample of $^{nat}$Ga-compound. Dependent on the system, different elution methods were used:

*Method A*: Isocratic elution with phosphate buffered saline (PBS, pH 7.4) and 20 mM arginine, flow rate: 0.35 mL min$^{-1}$. Electronic absorption was measured at 220 nm.

*Method B*: Isocratic elution with phosphate buffered saline (PBS, pH 7.4), 20 mM arginine and 10 mM EDTA, flow rate: 0.35 mL min$^{-1}$. Electronic absorption was measured at 280 nm.

**Spectroscopy**

Electronic absorption spectra were recorded using a Nanodrop$^{TM}$ One$^C$ Microvolume UV-Vis Spectrophotometer (ThermoFisher Scientific, supplied by Witec AG, Sursee, Switzerland). The same system was used to evaluate protein concentrations in accordance with the manufacturer's protocol.

**Size-exclusion chromatography coupled to multi-angle light scattering (SEC-MALS) detector**

The absolute mass of protein samples was determined by using a liquid chromatography system (Agilent LC1100 Agilent Technologies, Santa Clara, USA) coupled to an Optilab rEX refractometer (Wyatt Technology, Santa Barbara, USA) and a miniDAWN three-angle light-scattering detector (Wyatt Technology). For protein separation, a 24 mL Superdex 200 10/30 column (GE Healthcare Biosciences) was run in Tris-buffered saline (TBS150, pH 7.5) at 0.5 mL min$^{-1}$. Analysis was performed by using the ASTRA software (version 5.2.3.15; Wyatt Technology).

**Sodium dodecyl sulfate - polyacrylamide gel electrophoresis (SDS-PAGE)**

Samples (15 – 20 µL) were prepared for SDS-PAGE by mixing single proteins or reaction mixtures (1 – 3 µg of protein, or 20 – 30 mg of total protein for cell lysates), sample buffer (4 × lithium dodecyl sulfate (LDS)-Buffer for NuPage, supplemented with 0.05 M DTT unless otherwise stated) and phosphate buffered saline (PBS, pH 7.4). The resulting solution was heated at 80 °C for 10 min. (Labnet Accu Therm™). Samples (10 – 15 µL) including a protein ladder (Spectra™ Multicolor Broad Range Protein Ladder, ThermoFisher Scientific) were applied to a SDS-polyacrylamide gel (4 – 12% Bis-Tris, unless otherwise stated). Unless otherwise stated, gels were run in NuPage MES-SDS buffer (ThermoFisher Scientific), at 150 V for 60 min. Gels were washed in deionised water before staining or transfer procedures.

##### Native-PAGE analysis of proteins

Samples (15 – 20 µL) were prepared for native-PAGE by mixing single proteins or reaction mixtures with sample buffer (Native-TrisGly Sample Buffer 2×, NOVEX Life Technologies) and phosphate buffered saline (PBS, pH 7.4). The resulting solutions were mixed and samples (10 – 15 µL) including a protein ladder (Spectra™ Multicolor Broad Range Protein Ladder, ThermoFisher Scientific) were applied to a Bis-Tris Gel (Native-PAGE 4 – 16%, ThermoFisher Scientific). The gel was run in MES running buffer (0.05 M MES, pH 6) at 150 V for 60 min. The gel was washed with deionised water before the staining.

#### Radiochemical Methods

All instruments for measuring radioactivity were calibrated and maintained in accordance with previously reported routine quality control procedures.[1] For analysis appropriate background and decay corrections were applied as necessary.

##### Gallium-68

$[^{68}Ga][Ga(H_2O)_6]Cl_3$ (aq.) was obtained from $^{68}Ge/^{68}Ga$-generators (Eckert&Ziegler, Model IGG100 Gallium-68 Generator), and eluted with 0.1 M HCl (aq.). The eluted $^{68}Ga$ activity was trapped and purified by using a strong cation exchange column (Strata-XC, [SCX], Eckert&Ziegler). $[^{68}Ga][Ga(H_2O)_6]Cl_3$ (aq.) was eluted from the SCX cartridge by using a solution containing 0.13 M HCl (aq.) and approx. 5 M NaCl (aq.) (SCX eluent). For radiolabelling experiments, the $^{68}Ga$ stock solution (in ~0.1 M HCl, ca. 5 – 20 MBq) was typically added as the limiting reagent to an aqueous solution of unconjugated compounds (ca. $10^{-8}$ mol for PSMA-11; ca. $4 \times 10^{-9}$ mol for DARPins) buffered with NaOAc (0.1 M, pH 4.4). The reaction was monitored by using radio-iTLC and characterised by analytical size-exclusion chromatography.

##### Radio-TLC

Radioactive reactions were monitored by using instant thin-layer chromatography (iTLC). Glass-fibre iTLC plates impregnated with silica-gel (iTLC-SG, Agilent Technologies) were developed in citrate buffer (0.2 M, pH 4.5, >18.2 MΩ·cm $H_2O$) and analysed on a radio-TLC detector (SCAN-RAM, LabLogic Systems Ltd, Sheffield, United Kingdom). Radiochemical conversion (RCC) was determined by integrating the data obtained by the radio-TLC plate reader and determining both the percentage of radiolabelled product (retention factor $R_f = 0.0 – 0.1$) and 'free' $^{68}Ga$ ($R_f = 0.8 – 1.0$). Integration and data analysis were performed by using the software Laura version 5.0.4.29 (LabLogic).

##### Radioactive size-exclusion chromatography (radio-SEC)

Samples were analysed by using two methods of SEC. First, radiochemical purities (RCPs) of labelled protein samples were determined by using automated size-exclusion chromatography (Agilent AdvanceBio SEC 130 Å column, method B) as outlined above. Radiochemical purity (RCP) was determined by integrating the data obtained by the radio-SEC and determining both the percentage of radiolabelled product and 'free' $^{68}$Ga after appropriate background and decay corrections. Integration and data analysis were performed by using the software RadioStar version 5.0. As a second method, manual SEC procedures were used involving PD-10 (Sephadex G-25 resin, 85-260 μm, 14.5 mm ID × 50 mm, >30 kDa, GE Healthcare) or NAP$^{TM}$-25 columns (Cytiva) desalting columns. For analytical procedures, PD-10 and NAP-25 columns were eluted with sterile PBS. A total of 20 × 200 μL (PD-10), or 30 × 300 μL (NAP-25) fractions were collected up to a final elution volume of 6.5 mL (PD-10) or 11.5 mL NAP-25). The loading/dead-volume of the PD-10 and NAP-25 columns is precisely 2.50 mL which was discarded prior to aliquot collection. PD-10 and NAP-25 SEC columns were also used for preparative purification and reformulation of radiolabelled products by collecting a fraction of the eluate corresponding to the high molecular weight protein (>30 kDa fraction) eluted in ranges indicated for each experiment.

##### Quantification of radioactivity

For quantification of radioactivity, each fraction was measured on a gamma counter (HIDEX Automatic Gamma Counter, Hidex AMG, Turku, Finland) by using a counting time of 30 s and an energy window between 480 – 558 keV for $^{68}$Ga (511 keV emission), and 15 – 2047 keV for $^{111}$In. Appropriate background and decay corrections were applied throughout.

#### Cell culture

##### General methods in cell culture

All cells were cultured at 37 °C in a humidified 5% $CO_2$ atmosphere. All cell media was supplemented with fetal bovine serum (FBS, 10% [*v/v*], ThermoFisher Scientific) and penicillin/streptomycin (P/S, 1% [*v/v*] of penicillin 10,000 U mL$^{-1}$ and streptomycin 10 mg mL$^{-1}$). Cells were pelleted (100 g, 5 min.) and resuspended in media after trypsinisation.

##### LNCaP cells

The human prostate cancer cell line LNCaP was obtained from the American Type Culture Collection (ATCC-CRL-1740$^{TM}$, Manassas, VA). Cells were cultured at 37 °C in a humidified 5% $CO_2$ atmosphere in RPMI-1640 (without phenol-red) containing [+]-L-glutamine (2.5 mM). Cells were grown by serial passage and were harvested by using trypsin (0.1%).

##### Animals and xenograft models

All experiments involving mice were conducted in accordance with an animal experimentation license approved by the Zurich Canton Veterinary Office, Switzerland (Jason P. Holland). Experimental procedures also complied with guidelines issued in the *Guide for the Care and Use of Laboratory Animals.*[2] Female athymic nude mice (Crl:NU(NCr)-*Foxn1$^{nu}$*, 17 – 30 g, 6 – 8 weeks old) were obtained from Charles River Laboratories Inc. (Freiburg im Breisgau, Germany), and were allowed to acclimatise at the University of Zurich Laboratory Animal Services

Center vivarium for at least 1 week prior to implanting tumour cells. Mice were provided with food and water *ad libitum*. Tumours were induced on the right shoulder or flank by subcutaneous (s.c.) injection of LNCaP ($5.5 \times 10^6$) cells. The cells were injected in a 150-200 µL suspension of a 1:1 *v/v* mixture of cell medium (RPMI-1640) and reconstituted basement membrane (Corning® Matrigel® Basement Membrane Matrix, obtained from VWR International).[3] Tumours developed after a period of between 3 – 8 weeks. Tumour volume ($V$ / mm$^3$) was estimated by external Vernier caliper measurements of the longest axis, $a$ / mm, and the axis perpendicular to the longest axis, $b$ / mm. The tumours were assumed to be spheroidal and the volume was calculated in accordance with **Equation S1**. All mice injected with cancer cells developed tumours and the average volume of the LNCaP tumours was 624.56 ± 714.01 mm$^3$ (151.25 ± 78.46 mm$^3$, $n$ = 35 mice).

**Equation S1**. Approximation of tumour volume.

$$V = \frac{4\pi}{3} \cdot \left(\frac{a}{2}\right)^2 \cdot \left(\frac{b}{2}\right)$$

##### *Ex vivo* biodistribution studies

Biodistribution studies were conducted at the specified time points after completing the final measurements for estimating the effective half-live $t_{1/2}$(eff). Animals ($n$ = 3 – 4 mice per group) were anaesthetised individually by isoflurane and euthanised by isoflurane asphyxiation followed by terminal exsanguination. A total of 15 tissues (including the tumour) were removed, rinsed in water (except the fat, skin, muscle, and bone), dried in air for approx. 1 min., weighed and counted on a calibrated gamma counter (HIDEX Automated AMG Gamma Counter) for accumulation of activity. The mass of radiotracer formulation injected into each animal was measured and used to determine the total number of counts per minute (cpm) injected into each mouse by comparison to a standard syringe of known activity and mass. Count data were background- and decay-corrected, and the tissue uptake for each sample (determined in units of percentage injected dose per gram [%ID g$^{-1}$]) was calculated by normalisation to the total amount of activity injected for each individual animal.

##### Effective half-life measurement

The effective half-live $t_{1/2}$(eff) was calculated from the measurement of total internal radioactivity in the mouse models over time by using a dose calibrator (ISOMED 2010, NuviaTech Healthcare).

##### Statistical analysis

Data were analysed using GraphPad Prism and Excel. Unpaired, two-tailed Student's *t*-test were used with confidence levels of $p < 0.05$, $p < 0.01$, and $p < 0.001$ suggesting low, medium, and high statistical significance differences.

#### DARPin expression and selection

##### Biotinylation of PSA

PSA (250 µL, 1.5 mg mL$^{-1}$ , obtained from BBI™ Solutions) was re-buffered (Amicon 0.5, 10 kDa MWCO, 3,000 g, 15 min.) in PBS buffer (pH 7.4), and the concentration was measured to yield purified PSA (156.5 µL, 2.33 mg mL$^{-1}$ ). Biotin-NHS (6.2 µL, 62 nM 5 eq., 10 mM in DMSO) was added to purified PSA (156.5 µL, 368 µg, 12.58 nmol). The reaction was mixed by gentle pipetting and incubated at room temperature (30 min., 300 rpm). The reaction mixture was purified, to remove excess biotin, by spin filtration (Amicon 0.5, 10 kDa MWCO, 3,000 *g*,

15 min., 3 × 0.5 mL) into PBS + 0.1% w/v $NaN_3$. The purified PSA-biotin conjugate was then stored at 4 °C until further use.

**Ribosome display and FRET analysis**

To generate PSA-specific DARPin binders, biotinylated PSA was immobilized on either MyOne T1 streptavidin-coated beads (Pierce) or Sera-Mag neutravidin-coated beads (GE), depending on the selection round, and these beads were alternated. Ribosome display selections were performed similar to those described earlier,[4] using a semi-automatic KingFisher Flex MTP96 well platform.

The library includes N3C-DARPins with the original randomization strategy as reported[5] but includes a stabilized C-cap.[6–8] Additionally, the library is a mixture of DARPins with randomized and non-randomized N- and C-terminal caps, respectively[7,9] and successively enriched pools were cloned as intermediates in a ribosome display-specific vector.[9] Selections were performed over four rounds with decreasing target concentration and increasing washing steps to enrich for binders with high affinities.

The final enriched pool was cloned as fusion construct into a bacterial pQE30 derivative vector with an *N*-terminal MRGS$(H)_8$ tag and *C*-terminal FLAG tag *via* unique BamHI and HindIII sites containing *lacIq* for expression control. After transformation of *E. coli* XL1-blue, 380 single DARPin clones selected to bind to PSA were expressed in 96-well format and lysed by addition of B-Per Direct detergent plus Lysozyme and Nuclease (Pierce). These bacterial crude extracts of single DARPin clones were subsequently used in a Homogeneous Time Resolved Fluorescence (HTRF)-based screen to identify potential binders. Binding of the FLAG-tagged DARPins to streptavidin-immobilized biotinylated PSA was measured using FRET (donor: Streptavidin-Tb cryptate (610SATLB, Cisbio), acceptor: mAb anti FLAG M2-d2 (61FG2DLB, Cisbio). Further HTRF measurement against 'No Target' allowed for discrimination of PSA-specific hits. Experiments were performed at room temperature in white 384-well Optiplate plates (PerkinElmer) using the Taglite assay buffer (Cisbio) at a final volume of 20 µl per well. FRET signals were recorded after an incubation time of 30 minutes using a Varioskan LUX Multimode Microplate (Thermo Scientific). HTRF ratios were obtained by dividing the acceptor signal (665 nm) by the donor signal (620 nm) and multiplying this value by 10,000 to derive the 665/620 ratio. The background signal was determined by using reagents in the absence of DARPins.

From the initial hits of the 380 analysed DARPin clones, 32 were chosen for further analysis and their sequence determined. 27 were identified as true single clones. They were expressed in a 96-well format and IMAC purified. The purified DARPins were used in a hit validation to bind immobilized PSA by ELISA using an anti-FLAG antibody and SEC (see sections below).

#### ELISA of 27 DARPin constructs

ELISA binding of the 27 FLAG-tagged DARPins (abbreviated as DARPin) was evaluated by using the standard protocol (using anti-FLAG antibody; see above).

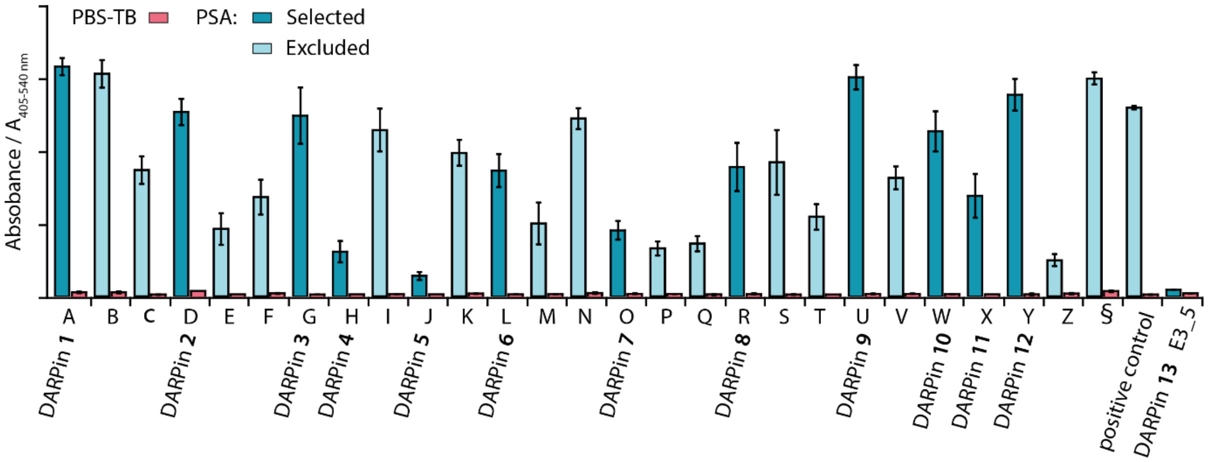

**Figure S1**. Bar chart showing the results from ELISA measurements. The specific binding of the 27 constructs towards PSA was tested. The 12 selected DARPins are highlighted in dark cyan. Data presented show the relative absorbance for PSA-binding DARPins (A – §) and the non-binding control (DARPin **13**, E3_5). Red bars show the non-specific binding control where no PSA was coated on the plate. As a positive control DARPin Off7 was added on a Maltose-Binding Protein (MBP) coated well. Error bars represent one standard deviation around the mean ($n$ = 2 measurements per sample).

##### Other exclusion criteria including SEC-MALS

SEC-MALS measurements of the 27 single clones were conducted in accordance with the standard procedure described above.

**Table S1.** Table of the 27 constructs showing a summary of the selection criteria: ELISA binding, SEC profile and sequence alignment. The 12 selected DARPins are highlighted in green, the exclusion reasons are highlighted in red.

| Initial DARPins | ELISA binding | SEC | Selected DARPins |
|---|---|---|---|
| A | high | monomeric | DARPin **1** |
| B | high | not monomeric | |
| C | low | broad | |
| D | high | monomeric | DAPRin **2** |
| E | low | monomeric | |
| F | low | monomeric | |
| G | high | monomeric | DARPin **3** |
| H | low | monomeric | DARPin **4** |
| I | high | not monomeric | |
| J | low | monomeric | DARPin **5** |
| K | medium | not monomeric | |
| L | medium | monomeric | DARPin **6** |
| M | low | monomeric | |
| N | high | not monomeric | |
| O | low | monomeric | DARPin **7** |
| P | low | monomeric | |
| Q | low | monomeric | |
| R | medium | monomeric | DARPin **8** |
| S | medium | not monomeric | |
| T | low | monomeric | |
| U | high | monomeric | DARPin **9** |
| V | medium | not monomeric | |
| W | high | monomeric | DARPin **10** |
| X | medium | monomeric | DARPin **11** |
| Y | high | monomeric | DARPin **12** |
| Z | low | monomeric | |
| § | high | not monomeric | |
| E3_5 | negative control | | DARPin **13** |

Based on ELISA and SEC-MALS results 12 PSA binding DARPins and one negative control (E_5, DARPin **13**) were selected. The corresponding amino acid sequences are shown in **Figure S2**.

| DARPin | N-cap | 1st repeat |
|---|---|---|
| DARPin | MRGSHHHHHHHHGSDLGKKLLEAAXXGQDDEVRILMANGADVNA | XDXXGYTPLHLAAXXGHLEIVEVLLKXGADVNA |
| 1 | MRGSHHHHHHHHGSDLGKKLLEAADNGQDDEVRILMANGADVNA | |
| 2 | MRGSHHHHHHHHGSDLGKKLLEAAETGQDDEVRILMANGADVNA | |
| 3 | MRGSHHHHHHHHGSDLGKKLLEAARAGQDDEVRILMANGADVNA | |
| 4 | MRGSHHHHHHHHGSDLGKKLLEAAWFGHLDEVRILMANGADVNA | |
| 5 | MRGSHHHHHHHHGSDLGKKLLEAAWAGQLDEVRILMANGADVNA | |
| 6 | MRGSHHHHHHHHGSDLGKKLLEAARAGQDDEVRILMANGADVNA | SDYVGYTPLHLAADYGHLEIVEVLLKTGADVNA |
| 7 | MRGSHHHHHHHHGSDLGKKLLEAATMGHLDEVRILMANGADVNA | |
| 8 | MRGSHHHHHHHHGSDLGKKLLEAARAGQDDEVRILMANGADVNA | |
| 9 | MRGSHHHHHHHHGSDLGKKLLEAAMRGHLDEVRILMANGADVNA | |
| 10 | MRGSHHHHHHHHGSDLGKKLLEAARAGQDDEVRILMANGADVNA | |
| 11 | MRGSHHHHHHHHGSDLGKKLLEAARAGQDDEVRILMANGADVNA | |
| 12 | MRGSHHHHHHHHGSDLGKKLLEAARAGQDDEVRILMANGADVNA | |
| 13 | MRGSHHHHHHHHGSDLGKKLLEAARAGQDDEVRILMANGADVNA | TDNDGYTPLHLAASNGHLEIVEVLLKNGADVNA |

| DARPin | 2nd repeat | 3rd repeat |
|---|---|---|
| DARPin | XDXXGXTPLHLAAXXGHLEIVEVLLKXGADVNA | XDXXGXTPLHLAAXXGHLEIVLLKHGADVNA |
| 1 | KDHIGSTPLHLAAIEGHLEIVEVLLKAGADVNA | TDWFGWTPLHLAADWGHLEIVLLKHGADVNA |
| 2 | WDVMGQTPLHLAALEGHLEIVEVLLKTGADVNA | VDAFGWTPLHLAAVDGHLEIVLLKHGADVNA |
| 3 | RDRLGQTPLHLAAIDGHLEIVEVLLKAGADVNA | TDQFGFTPLHLAAQEGHLEIVLLKHGADVNA |
| 4 | DDLLGQTPLHLAAMHGHLEIVEVLLKAGADVNA | RDYFGYTPLHLAAIWGHLEIVLLKHGADVNA |
| 5 | WDFIGQTPLHLAAYRGHLEIVEVLLKTGADVNA | QDYIGWTPLHLAAMQGHLEIVLLKHGADVNA |
| 6 | LDRIGQTPLHLAAWQGHLEIVEVLLKTGADVNA | IDYFGWTPLHLAASEGHLEIVLLKHGADVNA |
| 7 | QDMLGRTPLHLAAHNGHLEIVEVLLKTGADVNA | SDWFGWTPLHLAAFDGHLEIVLLKHGADVNA |
| 8 | HDYIGQTPLHLAASYGHLEIVEVLLKAGADVNA | QDFMGLTPLHLAAEWGHLEIVLLKHGADVNA |
| 9 | | QDQTGWTPLHLAAVYGHLEIVLLKAGADVNA |
| 10 | HDRLGQTPLHLAAMLGHLEIVEVLLKTGADVNA | TDYFGWTPLHLAAFDGHLEIVLLKTGADVNA |
| 11 | WDIIGNTPLHLAAMQGHLEIVEVLLKTGADVNA | ADVFGWTPLHLAAEYGHLEIVLLKHGADVNA |
| 12 | WDWLGQTPLHLAALEGHLEIVEVLLKTGADVNA | KDRFGWTPLHLAATDGHLEIVLLKHGADVNA |
| 13 | SDLTGITPLHLAAATGHLEIVEVLLKHGADVNA | YDNDGHTPLHLAAKYGHLEIVLLKHGADVNA |

| DARPin | C-cap | FLAG-tag |
|---|---|---|
| DARPin | QDKFGKTXFDLAIDNGNEDIAEVLQKAAKLN | DYKDDDDK |
| 1 | QDKFGKTAFDLAIDNGNEDIAEVLQKAAKLN | DYKDDDDK |
| 2 | QDVWGETAFDLAAWWGNEDIAEVLQKAAKLN | DYKDDDDK |
| 3 | QDKFGKTAFDLAIDNGNEDIAEVLQKAAKLN | DYKDDDDK |
| 4 | QDKFGKTAFDLAIDNGNEDIAEVLQKAAKLN | DYKDDDDK |
| 5 | QDKFGKTAFDLAIDNGNEDIAEVLQKAAKLN | DYKDDDDK |
| 6 | QDKFGKTAFDLAIDNGNEDIAEVLQKAAKLN | DYKDDDDK |
| 7 | QDKFGKTAFDISIDNGNEDIAEVLQKAAKLN | DYKDDDDK |
| 8 | QDKFGKTPFDLAIDNGNEDIAEVLQKAAKLN | DYKDDDDK |
| 9 | QDWYGETPFDLAAWEGNEDIAEVLQKAAKLN | DYKDDDDK |
| 10 | QDKFGKTAFDISIDNGNEDIAEVLQKAAKLN | DYKDDDDK |
| 11 | QDKFGKTPFDLAIDNGNEDIAEVLQKAAKLN | DYKDDDDK |
| 12 | QDKFGKTPFDLAIDNGNQDIAEVLQKAAKLN | DYKDDDDK |
| 13 | QDKFGKTAFDISIDNGNEDLAEILQK---LN | DYKDDDDK |

**Figure S2**. Amino acid sequence (single letter representation) of FLAG-tagged DARPins **1** – **13** showing the different repeats. The randomized regions are highlighted in blue. Note: The N-cap sequence of the FLAG-tagged DARPins features a His8-tag.

#### Cloning of PSA-DARPins with a Cys C-terminus

For the expression of Cys-*C*-terminally modified PSA-DARPins (DARPin-Cys) the cDNA encoding for PSA-DARPin was cloned into the *E. coli* expression plasmid pQI_E3_5_GGC (Containing DARPin E3_5 from the unselected library previously described[5]) *via* a *BamH*I/*Hind*III under the control of a T5 lac promoter to yield the expression vectors MRGS(H)$_6$ tag_PSA_DARPin_GGC (general amino acid sequence shown in **Figure S3**, with the modified regions highlighted in green).[10–12]

| | N-cap | 1st repeat |
|---|---|---|
| DARPin-Cys | MRGSHHHHHH--GSDLGKKLLEAAXXGQDDEVRILMANGADVNA | XDXXGYTPLHLAAXXGHLEIVEVLLKXGADVNA |
| | **2nd repeat** | **3rd repeat** |
| DARPin-Cys | XDXXGXTPLHLAAXXGHLEIVEVLLKXGADVNA | XDXXGXTPLHLAAXXGHLEIVLLKHGADVNA |
| | **C-cap** | **Cys-tag** |
| DARPin-Cys | QDKFGKTXFDXXIDNGNEDIAEVLQKAAKLN | GGC----- |

**Figure S3**. Amino acid sequence (single letter representation) of DARPin-Cys **1 – 13** showing the different repeats. The randomized regions are highlighted in blue, with the sequence being equivalent to FLAG-tagged DARPins **1** – **13** (**Figure S2**); the modified region is highlighted in green. Note: The *N*-cap sequence of the DARPin-Cys clones features a His$_6$-tag.

#### DARPin expression

For the expression of the DARPin-Cys constructs, the *E. coli* strain XL1-Blue (Stratagene, USA) was transformed with the plasmids described above. After overnight expression single clones were picked and used for the inoculation of 2YT medium (15 or 30 mL) supplemented with ampicillin (100 µg mL$^{-1}$) and glucose (1%). The inoculated medium was incubated overnight at 200 rpm and 37 ºC. These cultures were used to inoculate 200 mL or 500 mL cultures (2YT media supplemented with 100 µg $mL^{-1}$ ampicillin and 1% glucose) at an $OD_{600}$ of 0.1. Expression was induced with isopropyl β-D-thiogalactopyranoside (1 mM) at $OD_{600}$ of 0.8 – 1. After 4 h incubation at 200 rpm and 37 °C, the cultures were harvested by centrifugation for 15 min. (5,000 g, 4 °C). The pellet was washed with ice-cold PBS (30 mL, pH 7.4), centrifuged for 15 min (4,000 g, 4 °C) and stored at -20 °C until further purification.

**DARPin purification**

All DARPin-Cys proteins were purified by their $His_6$-tag using Ni-NTA immobilised metal-ion affinity chromatographic (IMAC) methods. All steps were performed at 4 °C. DARPins expressed in 200 mL cultures were thawed and resuspended in ice cold Ni-Lysis buffer (3.6 mL, 20 mM $Na_2HPO_4/NaH_2PO_4$, 500 mM NaCl, 1 mM DTT, pH 7.4) supplemented with 25 U $mL^{-1}$ Pierce™ Universal Nuclease for Cell Lysis (ThermoFischer Scientific). The cells were lysed by sonication and centrifuged for 30 min. (20,000 g, 4 °C). The lysates were supplemented with imidazole (20 mM) and glycerol (10% *v/v*). The proteins were purified *via* automated IMAC using a HisTrap FF crude (GE Healthcare) column on an ÄKTA Pure system (GE Healthcare). Columns were equilibrated with washing buffer (20 mM $Na_2HPO_4/NaH_2PO_4$, 500 mM NaCl, 20 mM imidazole, 10% *v/v* glycerol, 1 mM DTT, pH 7.4). Lysates were applied to the columns, washed with 5 column volumes (CV) of low-salt buffer (20 mM $Na_2HPO_4/NaH_2PO_4$, 137 mM NaCl, 20 mM imidazole, 10% *v/v* glycerol, 1 mM DTT, pH 7.4), 5 CV of high-salt buffer (20 mM $Na_2HPO_4/NaH_2PO_4$, 1 M NaCl, 20 mM imidazole, 10 % glycerol, 1 mM DTT, pH 7.4) and 5 CV of washing buffer. The proteins were eluted with 5 CV elution buffer (20 mM $Na_2HPO_4/NaH_2PO_4$, 500 mM NaCl, 250 mM imidazole, 10 % *v/v* glycerol, 1 mM DTT, pH 7.4). The proteins were buffer-exchanged using a HiTrap 26/10 Desalting (GE Healthcare) column. Protein-containing fractions were evaluated with a NanoDrop spectrometer, pooled, aliquoted, flash-frozen in liquid nitrogen and stored at -80 °C until further use. Protein purity and monomeric behaviour were determined to be >90% by Coomassie-stained SDS-PAGE and by MALS-SEC.

Bacterial pellets from 500 mL cultures were thawed and resuspended in ice-cold $TBS_{400}$ (15 − 20 mL, 50 mM Tris-HCl, 400 mM NaCl, pH 7.4) supplemented with 20 mM imidazole, glycerol (10% *v/v*), and 5 mM DTT. Lysozyme (1 g $L^{-1}$) was added before the cells were lysed by sonication and centrifuged for 20 min. (15,000 g, 4 °C). DARPins were purified *via* manual IMAC. Ni-NTA Superflow Resin (Quiagen) was packed in 1 – 2 mL benchtop columns (PD-10). Columns were equilibrated with lysis buffer (50 mM Tris-HCl, 400 mM NaCl, 20 mM imidazole, 1 mM DTT, 10% *v/v* glycerol, pH 7.4). Lysates were applied to the columns, washed with 2 × 10 CV washing buffer (50 mM Tris-HCl, 400 mM NaCl, 20 mM imidazole, 1 mM DTT, 10 % glycerol, pH 7.4), 10 CV of low-salt buffer (50 mM Tris-HCl, 20 mM NaCl, 20 mM imidazole, 10% *v/v* glycerol, 1 mM DTT, pH 7.4), 10 CV of high-salt buffer (50 mM Tris-HCl, 1 M NaCl, 20 mM imidazole, 1 mM DTT, 10% *v/v* glycerol, pH 7.4) and 10 CV of washing buffer (50 mM Tris-HCl, 400 mM NaCl, 20 mM imidazole, 1 mM DTT, 10% *v/v* glycerol, pH 7.4). The proteins were eluted with 2.5 CV elution buffer (50 mM Tris-HCl, 400 mM NaCl, 250 mM imidazole, 1 mM DTT, 10% *v/v* glycerol, pH 7.4). The eluate was dialysed overnight against PBS (pH 7.4, supplemented with 1 mM DTT). DARPins were evaluated for the purity by SDS-PAGE (**Figure S4**), aliquoted, flash-frozen in liquid nitrogen and stored at -80 °C until further use.

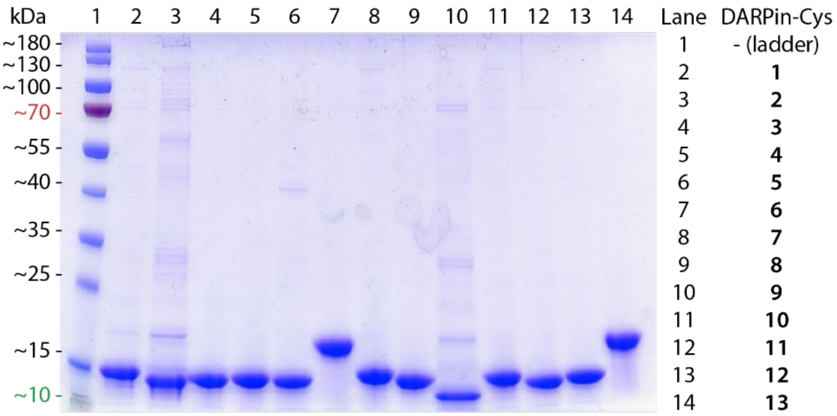

**Figure S4.** SDS PAGE analysis of PSA-DARPin-Cys where: (lane 1) molecular weight marker, (lane 2-14) PSA-DARPin-Cys **1** – **13**. Protein bands are stained with Coomassie blue.

**Size-exclusion chromatography coupled to multi-angle light scattering (SEC-MALS) detector**

The absolute mass of protein samples was determined by using a liquid chromatography system (Agilent LC1100 Agilent Technologies, Santa Clara, USA) coupled to an Optilab rEX refractometer (Wyatt Technology, Santa Barbara, USA) and a miniDAWN three-angle light-scattering detector (Wyatt Technology). For protein separation, a 24 mL Superdex 200 10/30 column (GE Healthcare Biosciences) was run in TBS150 (pH 7.5) at 0.5 mL min$^{-1}$. Analysis was performed by using the ASTRA software (version 5.2.3.15; Wyatt Technology). Note: For some of the smaller DARPin constructs a S75 column might have improved results of these measurements.

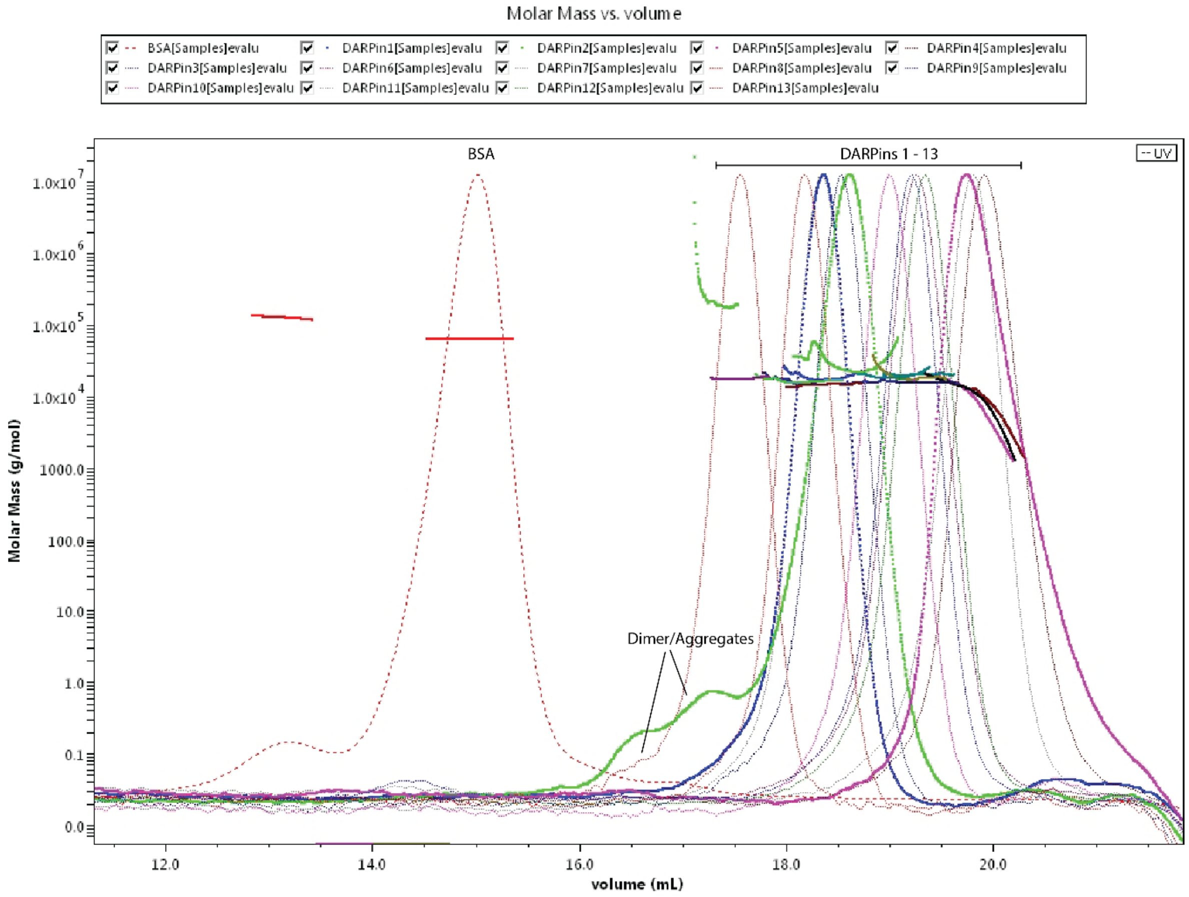

**Figure S5.** SEC-MALS traces showing the elution of BSA (red dotted line) as a standard and the 12 selected DARPin-Cys constructs with elution times between 17 and 20 min. DARPin-Cys **2** shows higher molecular weight aggregates, and DARPin-Cys **4**, **5** and **7** show peaks corresponding to a lower molecular weight species.

**Table S2.** Summary of the SEC-MALS analysis. The determined size and calculated mass are displayed for each DARPin-Cys construct. BSA was used as a standard.

| | | Monomer (Peak 1) | | | | Dimer (Peak 2) | | | |
|---|---|---|---|---|---|---|---|---|---|
| | Expected Mw (kDa) | Mw (kDa) | Uncertainty | Calculated mass (µg) | Mass fraction (%) | Mw (kDa) | Uncertainty | Calculated mass (µg) | Mass fraction (%) |
| BSA | 66.4 | 64.3 | 0.10% | 97.71 | 93.5 | 128.2 | 0.30% | 6.74 | 6.5 |
| DARPin-Cys **1** | 15.3 | 18.6 | 5.30% | 6.38 | 100 | | | | |
| DARPin-Cys **2** | 15.2 | 28.4 | 8.50% | 3.49 | 109.9 | 223.9 | 17.00% | -0.31 | -9.9 |
| DARPin-Cys **3** | 15.3 | 15.4 | 7.00% | 9 | 100 | | | | |
| DARPin-Cys **4** | 15.4 | 6.1 | 4.50% | 20.25 | 100 | | | | |
| DARPin-Cys **5** | 15.4 | 5.7 | 4.80% | 17.84 | 103.4 | | | -0.59 | -3.4 |
| DARPin-Cys **6** | 18.8 | 20.3 | 2.40% | 7.85 | 100 | | | | |
| DARPin-Cys **7** | 15.4 | 7.4 | 3.50% | 17.09 | 100 | | | | |
| DARPin-Cys **8** | 15.2 | 16.4 | 3.70% | 7.7 | 100 | | | | |
| DARPin-Cys **9** | 11.8 | 18.8 | 7.10% | 3.43 | 172 | | | -1.43 | -72 |
| DARPin-Cys **10** | 15.2 | 19.3 | 2.80% | 5.47 | 100 | | | | |
| DARPin-Cys **11** | 15.3 | 14.6 | 4.10% | 7.33 | 100 | | | | |
| DARPin-Cys **12** | 15.3 | 15.7 | 2.60% | 6.62 | 100 | | | | |
| DARPin-Cys **13** | 18.3 | 18 | 1.50% | 8.15 | 100 | | | | |
| Average | | 19.2 | | 15.59 | 105.6 | 176.1 | | 1.1 | -19.7 |
| Standard deviation | | 14.3 | | 24.22 | 19.4 | 67.7 | | 3.79 | 35.5 |
| % Standard deviation | | 74.6 | | 155.3 | 18.4 | 38.5 | | 344.38 | -180.1 |
| Minimum | | 5.7 | | 3.43 | 93.5 | 128.2 | | -1.43 | -72 |
| Maximum | | 64.3 | | 97.71 | 172 | 223.9 | | 6.74 | 6.5 |

### Bioconjugation of DARPin-Cys proteins with a NODAGA-maleimide

##### NODAGA-maleimide bioconjugation

The DARPin-Cys proteins (40 nmol) were reduced by incubation with an excess of DTT (8 µL, 0.5 M stock in water, 4 µmol, 100 eq. see **Table S3**A) for 30 min. (24 °C, 300 rpm). The excess DTT was removed by two subsequent spin filtrations (Zeba desalting column, 7 kDa MWCO, 0.5 mL). The DARPin-Cys proteins (21 to 39 nmol, see **Table S3**B) were then incubated with NODAGA-maleimide (2.14 to 3.92 µL, 2 eq., 20 mM in DMSO, see **Table S3**B) in PBS at pH 7.9 for 2 h (24 °C, 300 rpm). To remove excess NODAGA-maleimide, the reactions were purified with NAP-25 columns (Cytiva) pre-equilibrated with the running buffer (PBS, pH 7.4). The dead volume (2.6 mL) was discarded, and the high molecular weight fraction (1.6 mL) was collected and rebuffered into PBS and concentrated by spin-filtration (Amicon ultra 0.5 mL, 3 kDa MWCO, 5,000 g, 3 × 10 min.). Protein concentrations of the purified NODAGA-DARPin conjugates were measured by using a NanoDrop spectrometer, to estimate the recovered protein fractions (21 – 56% yield, see **Table S3**C). Aliquots of the crude reaction mixtures were retained for further radiochemical analysis.

**Table S3.** Table of (A) the utilised reaction conditions for DTT reduction, (B) maleimide conjugation reactions for DARPin-Cys **3**, **6**, **9**, **12**, and **13**; and (C) The protein concentrations and recovery yields for each NODAGA-DARPin construct.

| | DARPin **3** | DARPin **6** | DARPin **9** | DARPin **12** | DARPin **13** |
|---|---|---|---|---|---|
| (A) DTT treatment | | | | | |
| DARPin conc. / mg mL$^{-1}$ | 9.76 | 6.12 | 5.43 | 8.50 | 3.93 |
| DARPin conc. / µM | 640.00 | 324.88 | 458.42 | 553.94 | 214.77 |
| DARPin amount / µL | 62.50 | 123.12 | 87.26 | 72.21 | 186.25 |
| DARPin amount / nmol | 40.00 | 40.00 | 40.00 | 40.00 | 40.00 |
| DTT (0.5 M) / µL | 8.00 | 8.00 | 8.00 | 8.00 | 8.00 |
| DTT (0.5 M) / nmol | 4000.00 | 4000.00 | 4000.00 | 4000.00 | 4000.00 |
| (B) NODAGA-maleimide conjugation after rebuffering | | | | | |
| DARPin conc. / mg mL$^{-1}$ | 6.05 | 4.54 | 3.18 | 5.91 | 4.95 |
| DARPin conc. / µM | 396.36 | 241.16 | 268.48 | 384.85 | 270.69 |
| DARPin amount / µL | 54.00 | 120.00 | 90.00 | 74.00 | 145.00 |
| DARPin amount / nmol | 21.40 | 28.94 | 24.16 | 28.48 | 39.25 |
| NODAGA-maleimide (20uM in DMSO) /µL | 2.14 | 2.89 | 2.42 | 2.85 | 3.92 |
| NODAGA-maleimide (20uM in DMSO) /nmol | 42.81 | 57.88 | 48.33 | 56.96 | 78.50 |
| PBS / µL | 68.86 | 2.11 | 32.58 | 48.15 | 0.00 |
| Conc. DMSO / % | 1.71 | 2.32 | 1.93 | 2.28 | 2.64 |
| (C) After purification | | | | | |
| DARPin conc. / mg mL$^{-1}$ | 2.05 | 3.59 | 1.73 | 2.42 | 3.15 |
| DARPin conc. / µM | 134.55 | 190.45 | 145.79 | 157.58 | 172.26 |
| DARPin amount / µL | 92.57 | 116.80 | 108.87 | 54.53 | 91.40 |
| DARPin amount / nmol | 12.45 | 22.24 | 15.87 | 8.59 | 15.74 |
| yield recovered DARPin / % | 31.14 | 55.61 | 39.68 | 21.48 | 39.35 |

##### SDS-PAGE gel with DARPin-Cys constructs

Purified DARPin-Cys constructs were analysed by SDS-PAGE using the standard protocol. The gel (Bis-Tris, 12%) was run in MES-SDS buffer (see above). DTT was omitted from the loading buffer to analyse the extent of dimerisation *via* the formation of disulfide bridge.

##### Synthesis of $^{nat}$Ga-NODAGA-DARPin complexes

NODAGA-DARPins (~0.02 mg, ~1 nmol) were diluted in $H_2O$ (50 µL). $Ga(NO_3)_3$ (130 mM in $H_2O$, 1 µL, 130 eq.) was added. After incubation of the reaction for 15 min. at room temperature, the reaction was analysed by analytical HR-MS. The mass of the non-complexed conjugates was no longer be observed.

The different DARPin-Cys, NODAGA-DARPins and $^{nat}$Ga-NODAGA-DARPin complexes were analysed by liquid chromatography high resolution electrospray ionization mass spectrometry (LC-HR-ESI-MS). The obtained results are summarised in **Table S4** and shown in **Figure S6 – Figure S10**.

**Table S4.** High-resolution mass spectrometry analysis results with the calculated and observed *m/z* of the five DARPin-Cys, NODAGA-DARPin and $^{nat}$Ga-NODAGA-DARPin. Showing a shift in molecular mass of 597 *m/z* up on NODAGA conjugation and 66 *m/z* up on complexation with $^{nat}$Ga. Note: No peaks associated with non-conjugated DARPin-Cys were observed in the measurements for NODAGA-DARPins.

| | DARPin-Cys | | NODAGA-DARPin | | $^{nat}$GaNODAGA-DARPin | |
|---|---|---|---|---|---|---|
| | Calculated / *m/z* | Observed / *m/z* | Calculated / *m/z* | Observed / *m/z* | Calculated / *m/z* | Observed / *m/z* |
| DARPin **3** | 15255.13 | 15254.79 | 15752.34 | 15752.93 | 15818.24 | 15820.91 |
| DARPin **6** | 18842.18 | 18841.65 | 19339.39 | 19338.83 | 19405.29 | 19404.48 |
| DARPin **9** | 11854.27 | 11855.83 | 12351.48 | 12351.01 | 12417.38 | 12419.13 |
| DARPin **12** | 15353.28 | 15351.82 | 15850.49 | 15850.99 | 15916.39 | 15915.91 |
| DARPin **13** | 18285.34 | 18287.33 | 18782.55 | 18784.44 | 18848.45 | 18848.32 |

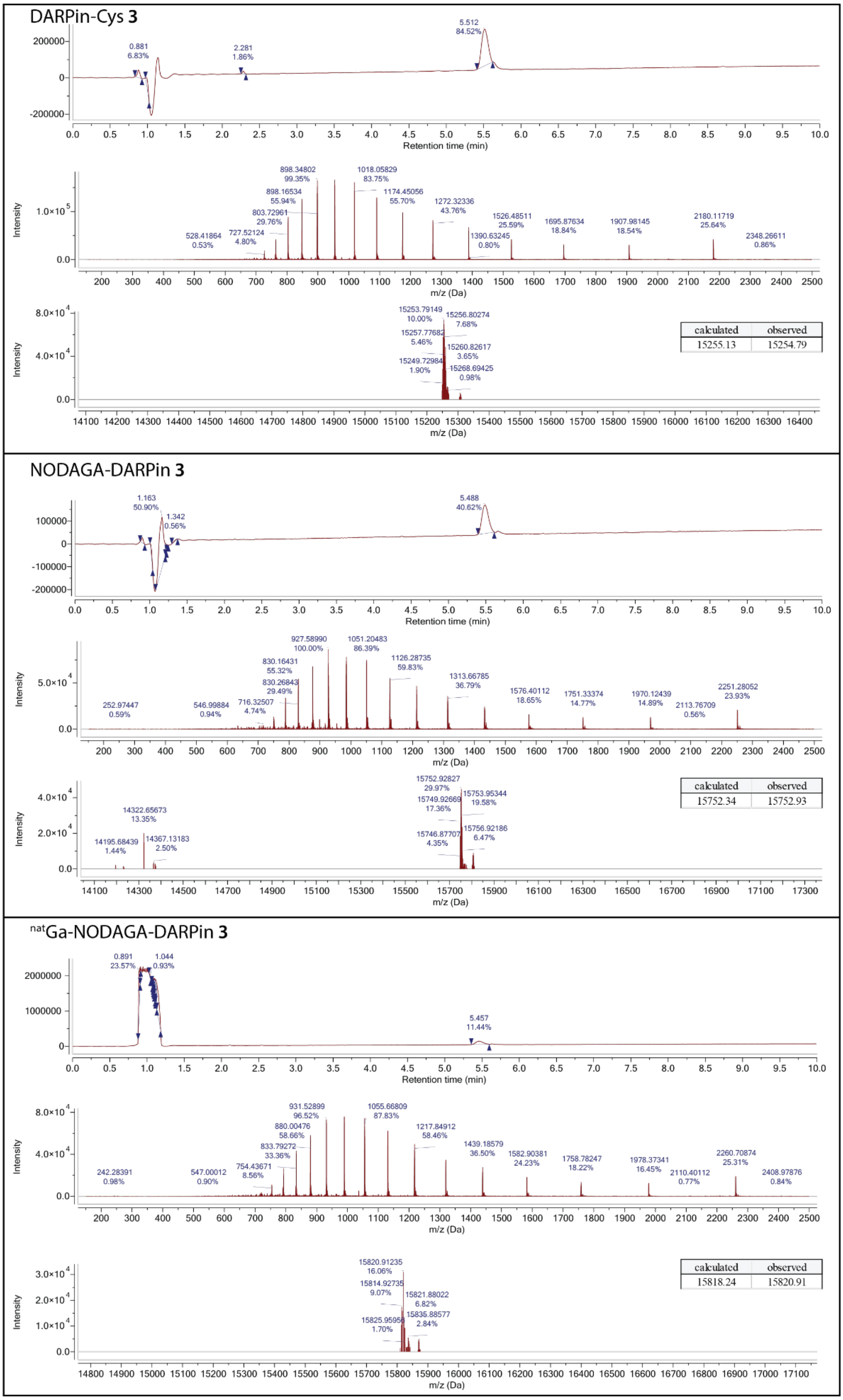

**Figure S6.** Mass-spectrum trace convoluted and de-convoluted HR-MS spectra of DARPin-Cys **3**, NODAGA-DARPin **3** and $^{nat}$Ga-NODAGA-DARPin **3**. Showing a shift in molecular mass of 597 *m*/*z* upon NODAGA conjugation and 66 *m*/*z* upon complexation with $^{nat}$Ga.

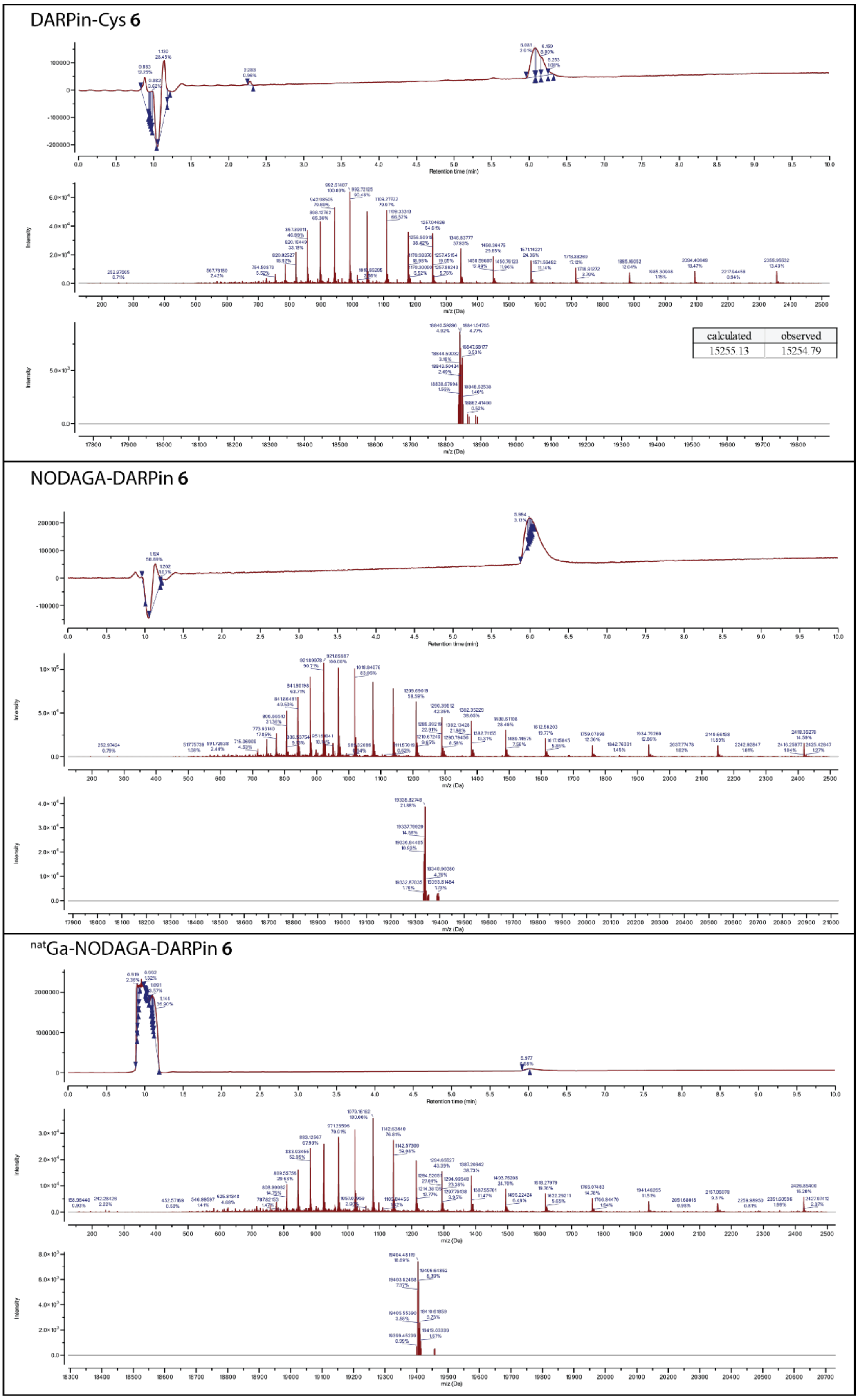

**Figure S7.** Mass-spectrum trace convoluted and de-convoluted HR-MS spectra of DARPin-Cys **6**, NODAGA-DARPin **6** and $^{nat}$Ga-NODAGA-DARPin **6**. Showing a shift in molecular mass of 597 *m*/*z* upon NODAGA conjugation and 66 *m*/*z* upon complexation with $^{nat}$Ga.

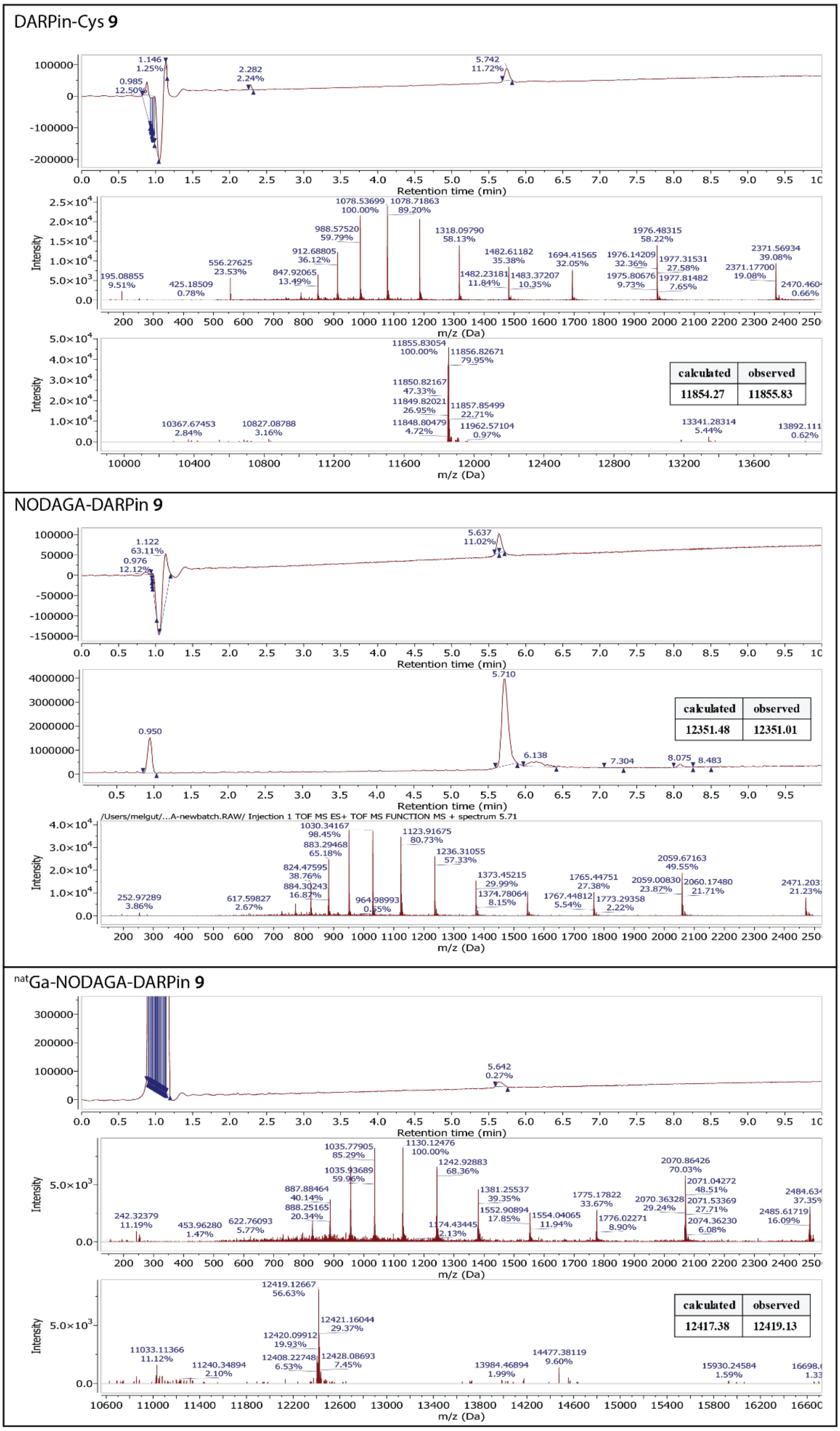

**Figure S8.** Mass-spectrum trace convoluted and de-convoluted HR-MS spectra of DARPin-Cys **9**, NODAGA-DARPin **9** and $^{nat}$Ga-NODAGA-DARPin **9**. Showing a shift in molecular mass of 597 *m*/*z* upon NODAGA conjugation and 66 *m*/*z* upon complexation with $^{nat}$Ga.

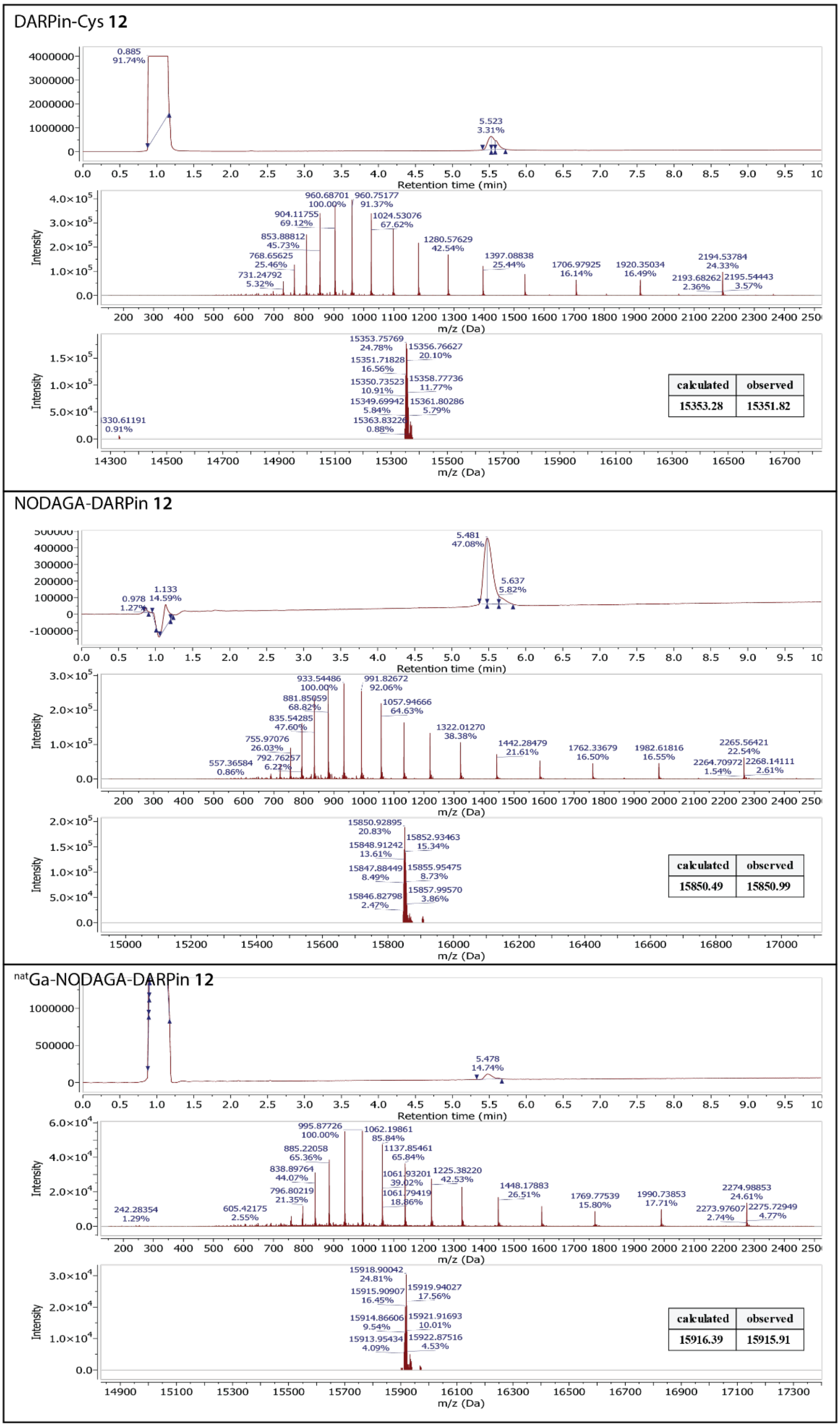

**Figure S9.** Mass-spectrum trace convoluted and de-convoluted HR-MS spectra of DARPin-Cys **12**, NODAGA-DARPin **12** and $^{nat}$Ga-NODAGA-DARPin **12**. Showing a shift in molecular mass of 597 *m*/*z* upon NODAGA conjugation and 66 *m*/*z* upon complexation with $^{nat}$Ga.

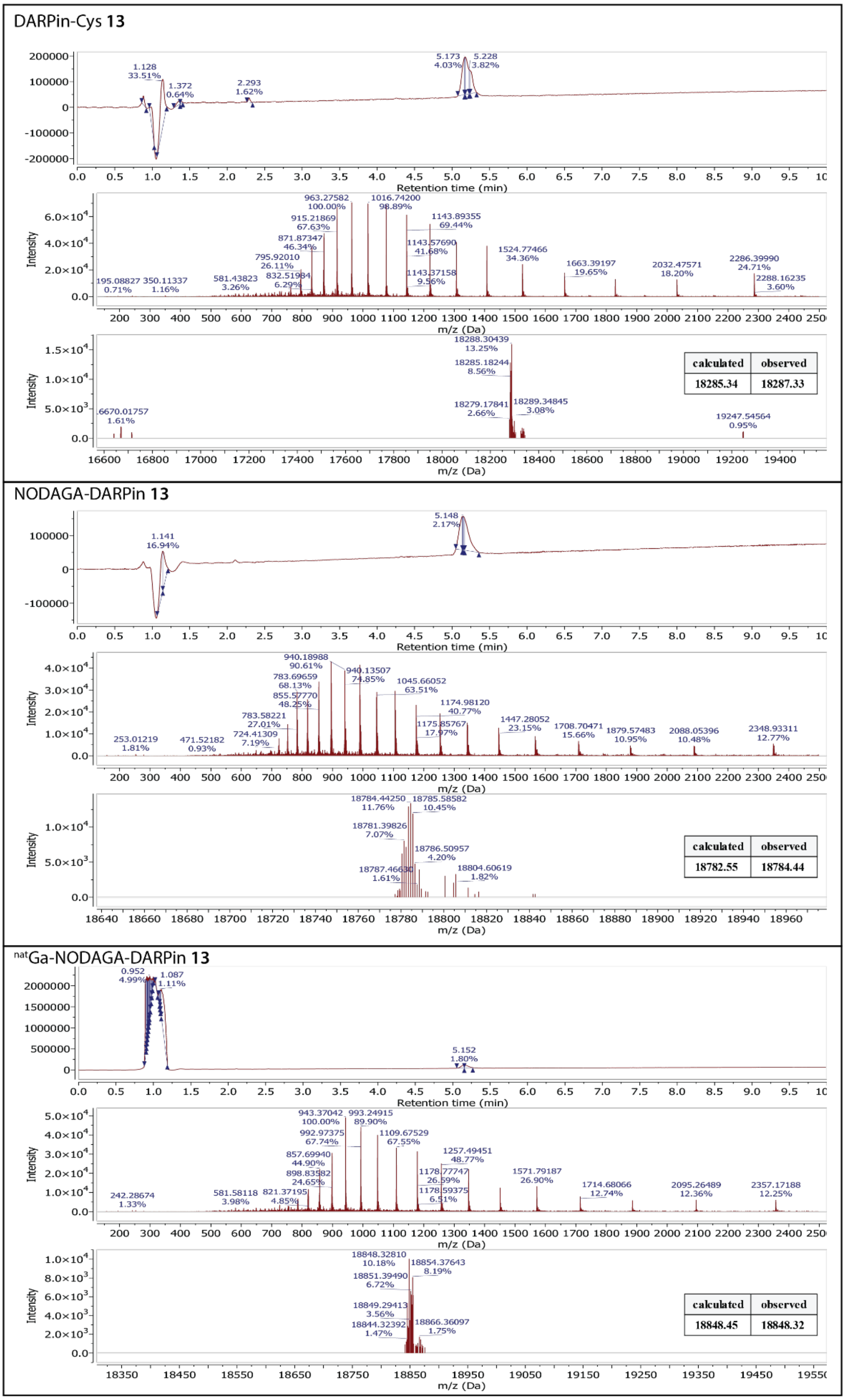

**Figure S10.** Mass-spectrum trace convoluted and de-convoluted HR-MS spectra of DARPin-Cys **13**, NODAGA-DARPin **13** and $^{nat}$Ga-NODAGA-DARPin **13**. Showing a shift in molecular mass of 597 *m*/*z* upon NODAGA conjugation and 66 *m*/*z* upon complexation with $^{nat}$Ga.

#### Qualitative assessment of PSA binding of NODAGA-DARPin constructs

*Evaluation of PSA binding of DARPin constructs after NODAGA modification by automated SEC chromatography*

The purified NODAGA-DARPin conjugates (7.5 µL, 125 µM, 937.5 pmol) were incubated with PSA (8 µL, 937.5 pmol, 1 eq.) at 24 °C (30 min., 400 rpm). These reaction mixtures (10 µL) were then analysed by using analytical SEC (method A). In the chromatograms, higher molecular weight complexes were observed at shorter retention times for all four PSA binding DARPins (**3**, **6**, **9**, and **12**), but not for the negative control (DARPin **13**).

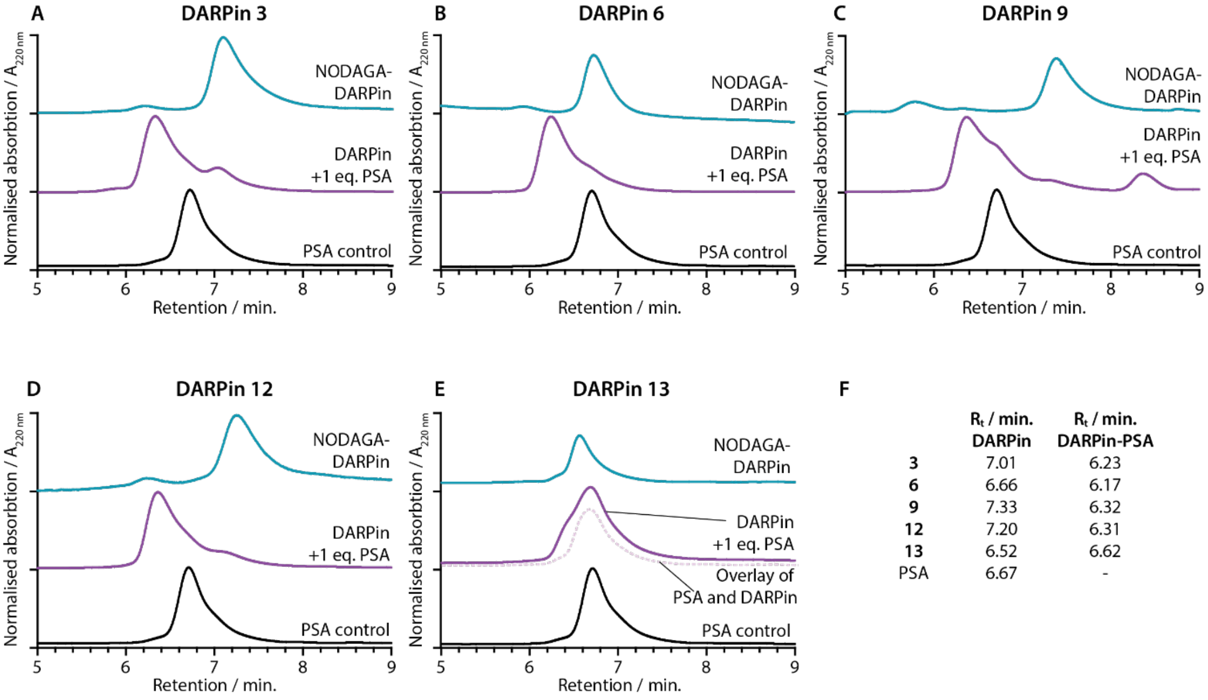

| | $R_t$ / min. DARPin | $R_t$ / min. DARPin-PSA |
|---|---|---|
| **3** | 7.01 | 6.23 |
| **6** | 6.66 | 6.17 |
| **9** | 7.33 | 6.32 |
| **12** | 7.20 | 6.31 |
| **13** | 6.52 | 6.62 |
| PSA | 6.67 | - |

**Figure S11.** SEC analysis of PSA binding. The NODAGA-DARPins **3**, **6**, **9**, **12**, and **13** (panels A – E, cyan) were incubated with 1 eq. of PSA (panels A – E, black) leading to the formation of a higher molecular weight complex with slightly shorter retention times (panels A – E, purple), except for the non-binding control DARPin **13**, where only a mixture of both proteins was detected and no complex formation is expected (theoretical overlay of PSA and DARPin **13** trace in pink dotted line). (F) Table of retention times of the NODAGA-DARPins and the NODAGA-DARPin-PSA complexes. Note: The absorption was measured at 220 nm.

*Native-PAGE gel analysis of PSA-DARPin-Cys complexes*

The purified NODAGA-DARPin conjugates (1.5 µL, 125 µM, 187.5 pmol) were incubated with PSA (1.6 µL, 187.5 pmol, 1 eq.) at 24 °C (30 min., 400 rpm). These reaction mixtures were then mixed with native loading buffer (Native-TrisGly Sample Buffer 2×, NOVEX Life Technologies), and the binding of the NODAGA-DARPins to PSA was evaluated by using native PAGE, as described above.

S

**Surface plasmon resonance measurements for quantitative assessment of PSA binding of DARPin-Cys and NODAGA-DARPins**

*The measurements and analyses were performed at the Functional Genomics Center Zürich (FGCZ)*

SPR measurements were performed at 20 °C in HBS buffer (10 mM HEPES, 150 mM NaCl, 0.005% Tween-20, 50 µM EDTA, pH 7.6) using a Biacore T200/S200 instrument (GE Healthcare/Cytiva). Proteins were immobilised on a multidentate poly-nitrilotriacetic acid (NTA)-derivatised linear polycarboxylate hydrogel NiHC30M chip, according to a protocol recommended by the chip manufacturer (Xantec, Düsseldorf, Germany). Firstly, a 0.35 M EDTA solution (pH 8.5) was injected for 300 s, followed by a 120 s buffer injection, a surface activation step with 5 mM $NiCl_2$ (in HBS, 20 µM EDTA) for 60 – 120 s, and a 40 – 60 s injection of DARPin-Cys (200 nM) or NODAGA-DARPins (125 nM), respectively, both containing a $His_6$ tag. This was followed by a 120 s injection of immobilization buffer. All immobilization steps were performed in immobilization buffer (10 mM HEPES, 150 mM NaCl, 0.005% Tween-20, and 50 µM EDTA) at a flow rate of 5 µL $min^{-1}$. This results in a surface density of 35 – 140 RU.

In the double-referenced binding experiments, a two-fold dilution series of five concentrations of PSA, PSA-ACT and ACT were injected for 120 s, at a flow rate of 30 µL $min^{-1}$ in HBS buffer. PSA concentrations were in the range of 69 – 142 nM, PSA-ACT concentrations in the range 1 – 16 nM, and ACT (from human plasma, Sigma-Aldrich) concentration in the range 1 – 16 nM, respectively. The chip surface was regenerated by injection of 0.35 M EDTA for 300 s. A concentration of 0.4 mg $mL^{-1}$ for the PSA-ACT complex stock solution (average of the concentration provided by the supplier: 0.3 – 0.5 mg $mL^{-1}$, Prostate Specific Antigen–α1-Antichymotrypsin Complex human, Sigma-Aldrich) and a molecular weight of 60 kDa were assumed.

The DARPin 13 coated flow cell 1 and 3 of the sensor chip were used as a reference. Data were evaluated by using Biacore software version 2.0.3. Due to unsatisfactory fits with a 1:1 model, most sensorgrams were fitted by using a sum of two exponentials (Sum2Exp, in BiaEvaluate v2.03 called heterogeneous ligand) kinetic fit. All obtained values are displayed in **Table S5** and **Table S6**; all measured sensorgrams are displayed in **Figure S12** – **Figure S15**.

**Table S5.** Results of single-cycle kinetic surface plasmon resonance (SPR) measurements of the DARPins-Cys **3**, **6**, **9**, and **12**. Determined rate constant of association ($k_{on}$ / $M^{-1}$ $s^{-1}$), rate constant of dissociation ($k_{off}$ / $s^{-1}$) and the dissociation constant ($K_D$ / $M^{-1}$) are given for each construct. For all ACT measurements no binding was detected (not detected, *n.d.*)

| Probe | Probe Density | Temp / °C | Buffer | Analyte | Model | $k_{on}$ / $10^5$ $M^{-1}$ $s^{-1}$ | $k_{off}$ / $10^{-3}$ s-1 | $K_D$ / nM | RU(max) | $Chi^2$ / $RU^2$ |
|---|---|---|---|---|---|---|---|---|---|---|
| DARPin-Cys **3** | 70 | 20 | HBS | PSA | Sum2Exp | 1.515 ± 0.003 | 2.440 ± 0.001 | 16.11 ± 0.06 | 117.1 ± 0.3 | 0.207 |
| | | | | | | 7.137 ± 0.065 | 12.92 ± 0.005 | 18.10 ± 0.18 | 55.92 ± 0.37 | |
| | | | | PSA-ACT | 1:1 | 6.966 ± 0.132 | 44.37 ± 0.30 | 63.69 ± 1.28 | 8.509 ± 0.115 | 0.0259 |
| | | | | ACT | *n.d.* | | | | | |
| DARPin-Cys **6** | 112 | 20 | HBS | PSA | Sum2Exp | 8.646 ± 0.038 | 13.88 ± 0.045 | 16.06 ± 0.09 | 34.67 ± 0.11 | 0.0659 |
| | | | | | | 1.201 ± 0.003 | 2.498 ± 0.004 | 20.79 ± 0.06 | 61.15 ± 0.09 | |
| | | | | PSA-ACT | 1:1 | 6.711 ± 0.083 | 18.27 ± 0.238 | 27.22 ± 0.49 | 13.43 ± 0.05 | 0.0387 |
| | | | | ACT | *n.d.* | | | | | |
| DARPin-Cys **9** | 110 | 20 | HBS | PSA | Sum2Exp | 1.104 ± 0.006 | 0.1460 ± 0.0064 | 1.322 ± 0.059 | 70.29 ± 0.27 | 0.440 |
| | | | | | | 8.924 ± 0.632 | 2.484 ± 0.007 | 2.783 ± 0.021 | 105.3 ± 0.3 | |
| | | | | PSA-ACT | Sum2Exp | 6.308 ± 0.096 | 4.400 ± 0.047 | 6.976 ± 0.130 | 1.556 ± 0.016 | 0.0302 |
| | | | | | | 10.57 ± 0.070 | 53.00 ± 0.26 | 50.13 ± 0.41 | 11.15 ± 0.04 | |
| | | | | ACT | *n.d.* | | | | | |
| DARPin-Cys **12** | 90 | 20 | HBS | PSA | Sum2Exp | 9.484 ± 0.063 | 3.022 ± 0.009 | 3.186 ± 0.023 | 41.51 ± 0.09 | 0.0690 |
| | | | | | | 0.8821 ± 0.0043 | 0.2977 ± 0.0057 | 3.375 ± 0.067 | 33.91 ± 0.08 | |
| | | | | PSA-ACT | Sum2Exp | 7.961 ± 0.029 | 2.281 ± 0.013 | 2.865 ± 0.020 | 25.27 ± 0.15 | 0.345 |
| | | | | | | 4.084 ± 0.035 | 12.61 ± 0.13 | 30.87 ± 0.41 | 20.38 ± 0.12 | |
| | | | | ACT | *n.d.* | | | | | |

**Table S6.** Results of single-cycle kinetic surface plasmon resonance (SPR) measurements of the NODAGA-DARPins **3**, **6**, **9**, and **12**. Determined rate constant of association ($k_{on}$ / $M^{-1}$ $s^{-1}$), rate constant of dissociation ($k_{off}$ / $s^{-1}$) and the dissociation constant ($K_D$ / $M^{-1}$) are given for each construct. For all ACT measurements no binding was detected (not detected, *n.d.*)

| Probe | Probe Density | Temp / °C | Buffer | Analyte | Model | $k_{on}$ / $10^5$ $M^{-1}$ $s^{-1}$ | $k_{off}$ / $10^{-3}$ s-1 | $K_D$ / nM | RU(max) | $Chi^2$ / $RU^2$ |
|---|---|---|---|---|---|---|---|---|---|---|
| NODAGA-DARPin **3** | 49 | 20 | HBS | PSA | Sum2Exp | 8.824 ± 0.417 | 0.8728 ± 0.0073 | 0.9891 ± 0.009 | 53.73 ± 0.35 | 0.715 |
| | | | | | | 2.243 ± 0.034 | 4.046 ± 0.043 | 18.04 ± 0.33 | 25.98 ± 0.30 | |
| | | | | PSA-ACT | Sum2Exp | 0.2488 ± 0.0089 | 0.01391 ± 0.00079 | 0.5691 ± 0.0037 | 61.21 ± 1.93 | 2.53 |
| | | | | | | 22.50 ± 0.18 | 2.923 ± 0.029 | 1.299 ± 0.014 | 108.8 ± 0.1 | |
| | | | | ACT | *n.d.* | | | | | |
| NODAGA-DARPin **6** | 54 | 20 | HBS | PSA | Sum2Exp | 5.399 ± 0.027 | 0.9626 ± 0.0062 | 1.783 ± 0.015 | 67.94 ± 0.30 | 0.618 |
| | | | | | | 1.432 ± 0.041 | 6.184 ± 0.130 | 43.18 ± 1.54 | 16.40 ± 0.20 | |
| | | | | PSA-ACT | Sum2Exp | 10.26 ± 0.076 | 0.9818 ± 0.0039 | 0.9566 ± 0.0080 | 90.69 ± 0.2378 | 1.05 |
| | | | | | | 1.033 ± 0.037 | 6.542 ± 0.089 | 63.33 ± 2.40 | 46.38 ± 0.95 | |
| | | | | ACT | *n.d.* | | | | | |
| NODAGA-DARPin **9** | 44 | 20 | HBS | PSA | Sum2Exp | 46.95 ± 0.39 | 16.04 ± 0.098 | 3.416 ± 0.036 | 31.94 ± 0.07 | 0.295 |
| | | | | | | 2.428 ± 0.010 | 2.536 ± 0.011 | 10.44 ± 0.06 | 29.05 ± 0.07 | |
| | | | | PSA-ACT | Sum2Exp | 2.028 ± 0.006 | 0.07143 ± 0.00173 | 0.3523 ± 0.0086 | 133.1 ± 0.1 | 0.0515 |
| | | | | | | 20.21 ± 0.15 | 2.049 ± 0.005 | 1.014 ± 0.008 | 60.64 ± 0.10 | |
| | | | | ACT | *n.d.* | | | | | |
| NODAGA-DARPin **12** | 62 | 20 | HBS | PSA | Sum2Exp | 16.05 ± 0.08 | 9.007 ± 0.030 | 5.613 ± 0.039 | 28.05 ± 0.09 | 0.214 |
| | | | | | | 1.683 ± 0.006 | 1.438 ± 0.005 | 8.543 ± 0.045 | 34.69 ± 0.09 | |
| | | | | PSA-ACT | Sum2Exp | 0.8394 ± 0.0080 | 0.6479 ± 0.0018 | 7.706 ± 0.077 | 129.2 ± 0.8 | 0.0580 |
| | | | | | | 5.804 ± 0.174 | 12.35 ± 0.11 | 21.29 ± 0.66 | 50.83 ± 0.29 | |
| | | | | ACT | *n.d.* | | | | | |

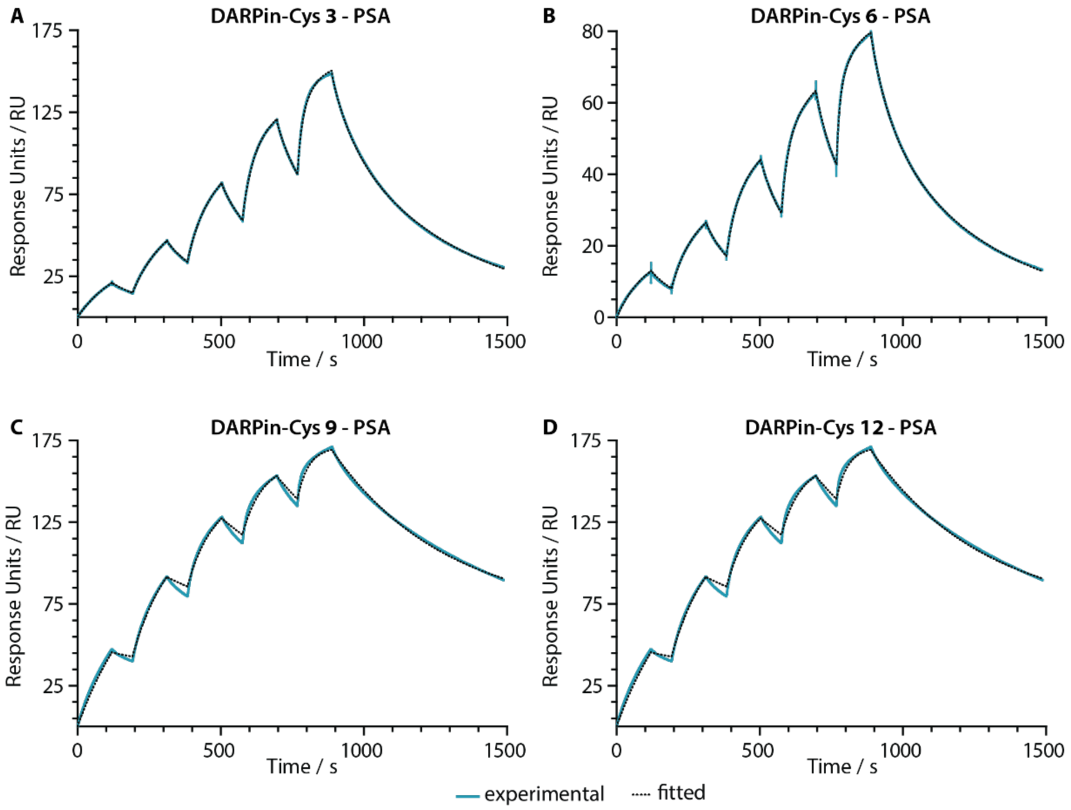

**Figure S12.** Surface plasmon resonance (SPR) sensorgrams of DARPin-Cys **3**, **6**, **9**, **12** indicating binding to the analyte PSA. Experimental data are displayed in cyan; the black dashed line shows Sum2Exp fit.

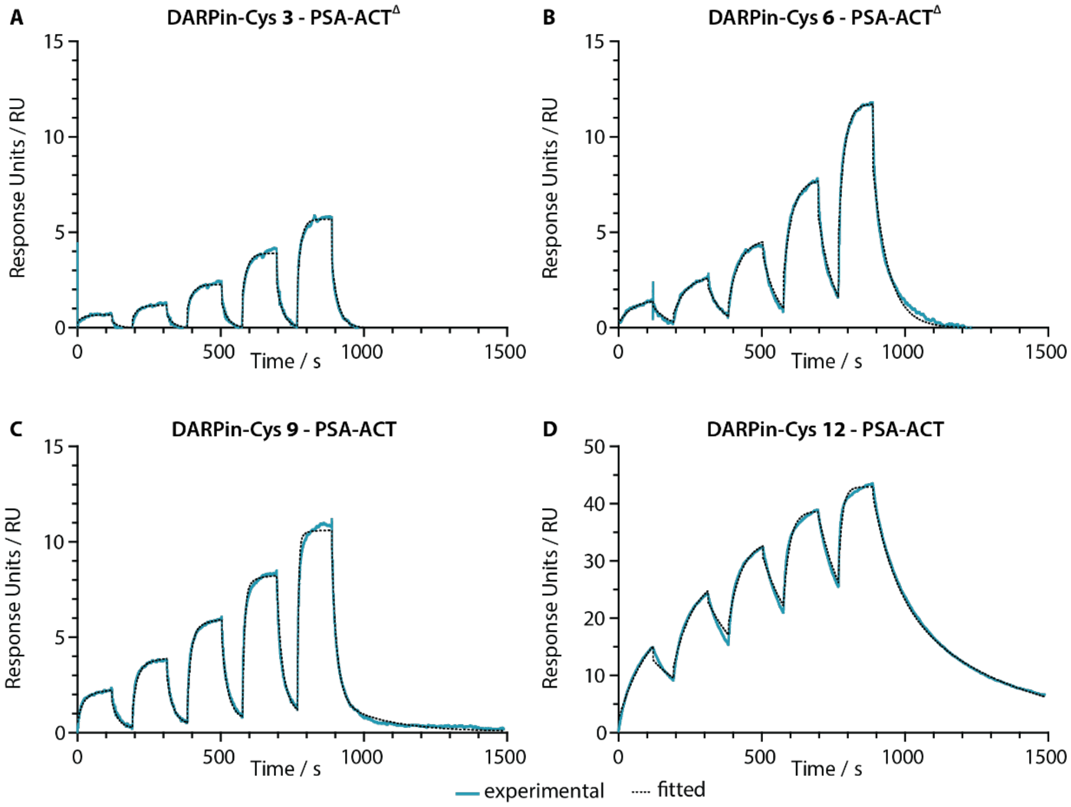

**Figure S13.** Surface plasmon resonance (SPR) sensorgrams of DARPin-Cys **3**, **6**, **9**, **12** indicating binding to the analyte PSA-ACT. Experimental data are displayed in cyan; the black dashed line shows Sum2Exp fit. $^{\Delta}$ Data were fitted with a 1:1 model.

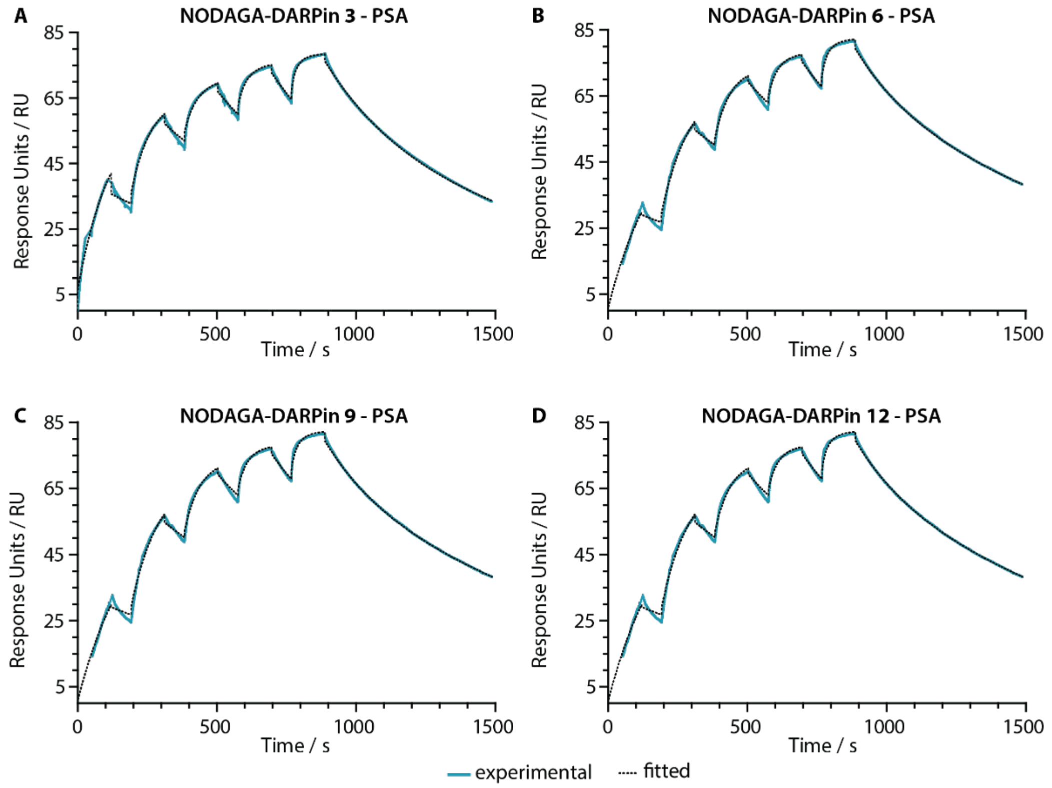

**Figure S14.** Surface plasmon resonance (SPR) sensorgrams of NODAGA-DARPin **3**, **6**, **9**, **12** indicating binding to the analyte PSA. Experimental data are displayed in cyan; the black dashed line shows Sum2Exp fit.

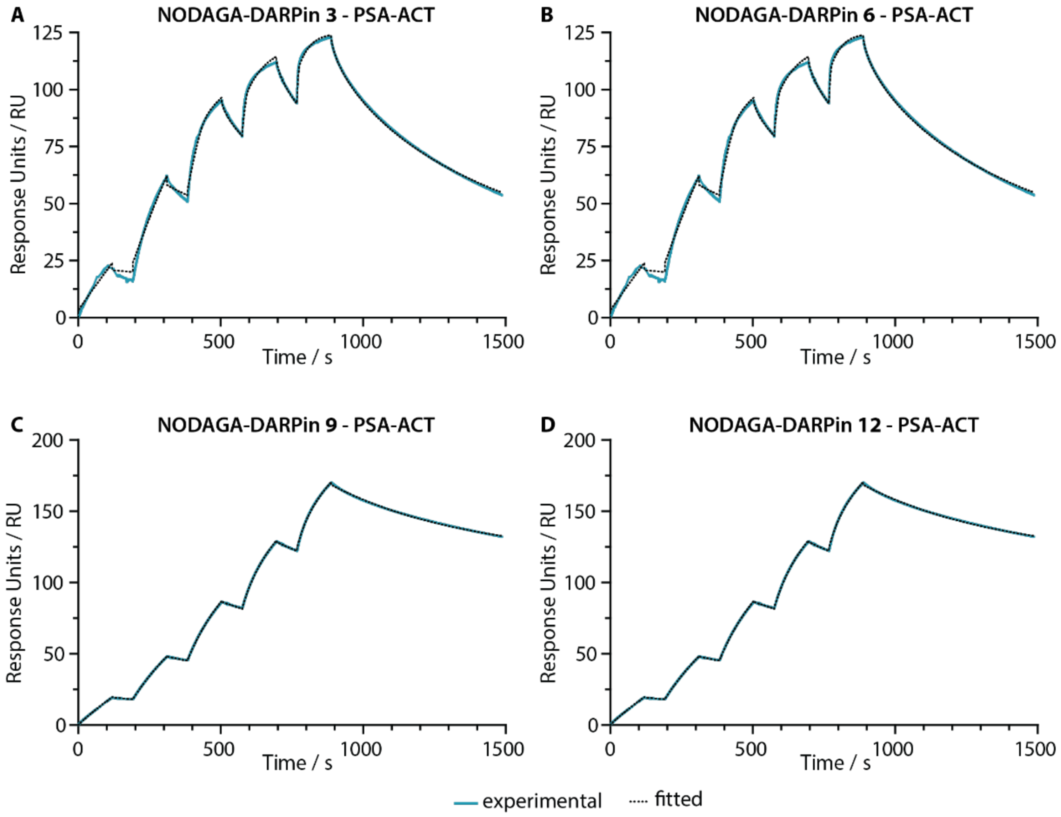

**Figure S15.** Surface plasmon resonance (SPR) sensorgrams of NODAGA-DARPin **3**, **6**, **9**, **12** indicating a binding to the analyte PSA-ACT. Experimental data are displayed in cyan; the black dashed line shows Sum2Exp fit.

#### Radiolabelling of NODAGA-DARPins

##### $^{68}$Ga-radiolabelling of NODAGA-DARPins

NODAGA-DARPins (3.0 µL, ~6 µg, ~0.40 nmol) were buffered in NaOAc (50 µL, 0.1 M, pH 4.4). Radiolabelling was achieved by addition of an aliquot of [$^{68}$Ga][Ga($H_2O$)$_6$]$Cl_3$ (8.5 µL, ~4 MBq) and adjustment of the pH to 5.3 – 5.5 with $Na_2CO_3$ (1.5 µL, 1 M). After incubation of the reaction for 10 min. at room temperature, the [$^{68}$Ga]GaNODAGA-DARPin complexes were analysed by radio-TLC, manual NAP-25 and radio-SEC (**Figure S16**).

##### Molar activities of $^{68}$Ga-radiollabelling of [$^{68}$Ga]GaNODAGA-DARPins

Varying amounts of NODAGA-DARPins were radiolabelled with the same amounts of radioactivity by using the standard procedure described above. After incubation of the reaction mixtures for 10 min. at room temperature, the reaction was analysed by radio-SEC (method B) and the RCPs were determined by integration of the peaks. The molar activities were determined to be 30.72 to 54.37 MBq nmol$^{-1}$ of protein (**Table S7**).

**Table S7.** Summary of molar activity values in MBq nmol$^{-1}$ obtained by titrations of different amounts of the five NODAGA-DARPins with a constant amount of [$^{68}$Ga][Ga($H_2O$)$_6$]$Cl_3$.

| [$^{68}$Ga]GaNODAGA-DARPin | **3** | **6** | **9** | **12** | **13** |
|---|---|---|---|---|---|
| Specific activity / MBq nmol$^{-1}$ | 36.21 | 37.16 | 30.72 | 54.37 | 39.82 |

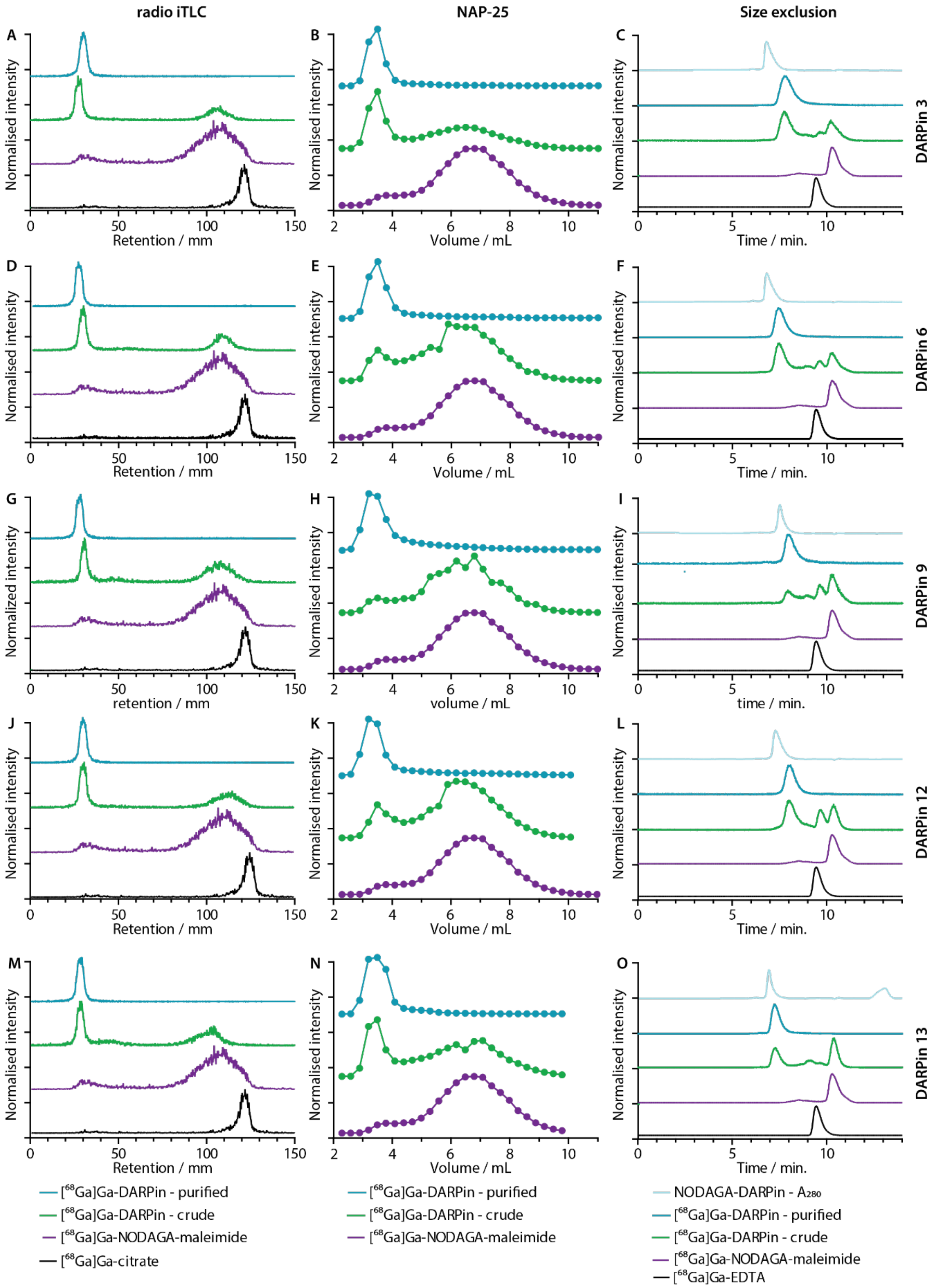

**Figure S16.** (Panels A, D, G, J, and M) Radio-TLC analysis data showing characterisation of [$^{68}$Ga]GaNODAGA-DARPins **3**, **6**, **9**, **12**, and **13** after purification (cyan), as a crude (green) in citrate buffer. The purple and black traces show controls with [$^{68}$Ga]Ga-NODAGA-maleimide, and [$^{68}$Ga]Ga-citrate. (Panels B, E, H, K, and N) Radio-NAP-25 SEC analysis of the purified (cyan) and the crude (green) constructs, and [$^{68}$Ga]Ga-NODAGA-maleimide (purple). (Panels C, F, I, L, and O) Radio-SEC analysis showing co-elution of the NODAGA-DARPins **3**, **6**, **9**, **12**, and **13** UV trace (light blue) with the [$^{68}$Ga]GaNODAGA-DARPins **3**, **6**, **9**, **12**, and **13** radiotrace (cyan). The crude (green) construct, [$^{68}$Ga]Ga-NODAGA-maleimide (purple), and [$^{68}$Ga]Ga-EDTA control complex (black).

**Stability of [$^{68}$Ga]GaNODAGA-DARPins**

The stability of the five [$^{68}$Ga]GaNODAGA-DARPins was investigated with respect to change in radiochemical purity (RCP). The results are summarised below (**Figure S17**)

*Incubation in PBS*

Radiolabelled [$^{68}$Ga]GaNODAGA-PSA-DARPins (30 µL, ~3 µg, ~0.20 nmol, ~2 MBq) were incubated with PBS (pH 7.4, 60 µL) at 37 °C. Aliquots of the reactions were analysed by automated Radio-SEC (method B) at time intervals of 0, 30, 60, 90, and 120 min. The RCP are summarised in **Table S8**.

**Table S8.** Table showing the RCP of [$^{68}$Ga]GaNODAGA-DARPins over time after incubation in PBS at 37 °C as determined by automated radio-SEC.

| Time / min. | [$^{68}$Ga]GaNODAGA-DARPin **3** RCP / % | [$^{68}$Ga]GaNODAGA-DARPin **6** RCP / % | [$^{68}$Ga]GaNODAGA-DARPin **9** RCP / % | [$^{68}$Ga]GaNODAGA-DARPin **12** RCP / % | [$^{68}$Ga]GaNODAGA-DARPin **13** RCP / % |
|---|---|---|---|---|---|
| 0 | 100 | 100 | 100 | 100 | 100 |
| 30 | 100 | 100 | 100 | 100 | 100 |
| 60 | 99 | 100 | 100 | 98 | 100 |
| 90 | 99 | 99 | 100 | 97 | 100 |
| 120 | 98 | 99 | 100 | 99 | 100 |

*Incubation with excess EDTA*

Radiolabelled [$^{68}$Ga]GaNODAGA-PSA-DARPins (30 µL, ~3 µg, ~0.20 nmol, ~2 MBq) were incubated with EDTA (pH 7.1, 30 µL, 50 mM, 1.5 µmol, 3,700 fold excess) at room temperature, and the RCP was monitored by radio-*i*TLC at 0, 30, 60, 90, and 120 min. post EDTA addition. The RCPs are summarised in **Table S9**.

**Table S9.** Table showing the RCP of [$^{68}$Ga]GaNODAGA-DARPins over time after EDTA addition (evaluated by radio-TLC).

| Time / min. | [$^{68}$Ga]GaNODAGA-DARPin **3** RCP / % | [$^{68}$Ga]GaNODAGA-DARPin **6** RCP / % | [$^{68}$Ga]GaNODAGA-DARPin **9** RCP / % | [$^{68}$Ga]GaNODAGA-DARPin **12** RCP / % | [$^{68}$Ga]GaNODAGA-DARPin **13** RCP / % |
|---|---|---|---|---|---|
| 0 | 100 | 100 | 100 | 100 | 100 |
| 30 | 100 | 98 | 97 | 99 | 98 |
| 60 | 97 | 98 | 95 | 97 | 99 |
| 90 | 95 | 98 | 92 | 97 | 100 |
| 120 | 95 | 96 | 92 | 96 | 99 |

*Incubation in human serum*

Radiolabelled [$^{68}$Ga]GaNODAGA-PSA-DARPins (40 µL, ~4 µg, ~0.27 nmol, ~2.7 MBq) were incubated with human serum (80 µL) at 37 °C, pH 7.4. Aliquots of the reactions were analysed by automated radio-SEC (method B) at time intervals of 0, 30, 60, 90, and 120 min. The RCP are summarised in **Table S10**.

**Table S10.** Table showing the RCP of [$^{68}$Ga]GaNODAGA-DARPins over time after incubation with human serum at 37 °C , (evaluated by automated Radio-SEC).

| Time / min. | [$^{68}$Ga]GaNODAGA-DARPin **3** RCP / % | [$^{68}$Ga]GaNODAGA-DARPin **6** RCP / % | [$^{68}$Ga]GaNODAGA-DARPin **9** RCP / % | [$^{68}$Ga]GaNODAGA-DARPin **12** RCP / % | [$^{68}$Ga]GaNODAGA-DARPin **13** RCP / % |
|---|---|---|---|---|---|
| 0 | 100 | 100 | 100 | 100 | 100 |
| 30 | 96 | 98 | 97 | 99 | 100 |
| 60 | 97 | 98 | 97 | 99 | 94 |
| 90 | 99 | 99 | 97 | 100 | 99 |
| 120 | 99 | 99 | 97 | 100 | 99 |

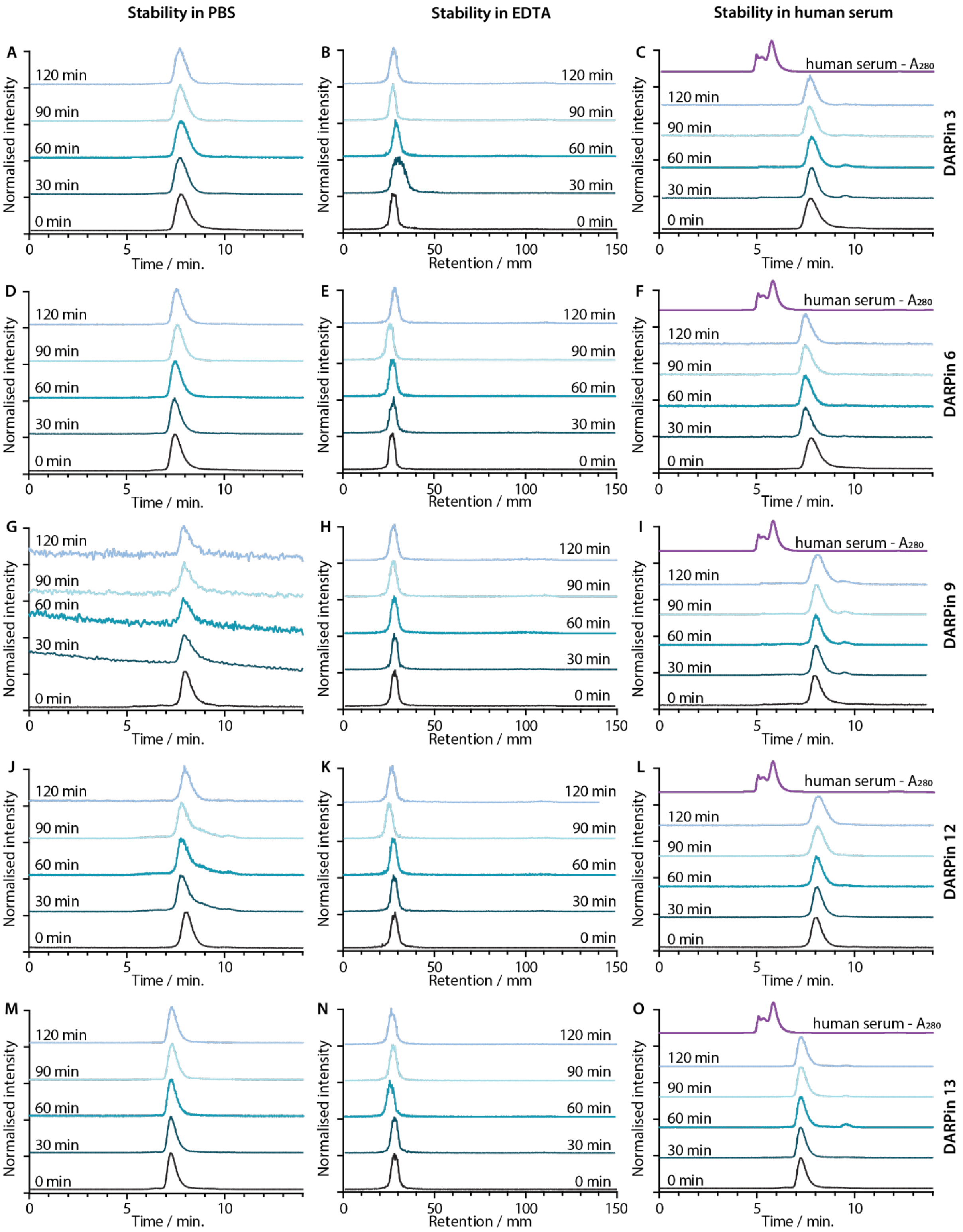

**Figure S17**. (Panels A, D, G, J and M) Radio-SEC chromatograms following the change in RCP of [$^{68}$Ga]GaNODAGA-DARPins over time after PBS (pH 7.4) addition at 37 °C. (Panels B, E, H, K and N) Radio-TLC chromatograms developed with citrate eluent (0.2 M, pH 4.5) following the change in RCP of [$^{68}$Ga]GaNODAGA-DARPins over time after EDTA addition at 24 °C, pH 7.1. (Panels C, F, I, L and O) Radio-SEC following the change in RCP of [$^{68}$Ga]GaNODAGA-DARPins over time after human serum addition at 37 °C, pH 7.4.

**Radioactive PSA-binding assay**

Since PSA is a secreted protein, the immunoreactivity could not be determined by using a cellular-based procedure adapted from Lindmo and co-workers.[13] Instead, a radioactive immune capture-type assay was developed to confirm binding.

[$^{68}$Ga]GaNODAGA-PSA-DARPins were synthesised by using the standard procedure stated above (reaction volume 60 µL). The reaction was diluted (1 mL, PBS + 0.3% *w/v* BSA), and the diluted radiotracer [$^{68}$Ga]GaNODAGA-PSA-DARPin (6 – 9 µL, 2 pmol, ~26 kBq) was pre-incubated with biotinylated-PSA (85 µg mL$^{-1}$, 2.9 µM, 6.8 µL, 20 pmol, 10 eq.) at 4 °C (300 rpm, 30 min.). For blocking experiments, non-radiolabelled DARPin-Cys (250 eq. 1 – 2 µL, 500 pmol) were added to biotin-PSA 15 min before the addition of the radiotracer. As a second control, biotin-PSA was omitted from the assay and substituted with PBS. Three samples of the same specified quantity of radiotracer were also prepared to serve as standards for total activity added to each sample. Then streptavidin-coated magnetic beads (5 µL, washed 2 × with PBS + *w/v* 0.3% BSA prior to use) were added and the reactions were incubated at 4 °C (10 min., 800 rpm). The magnetic beads were then pulled down and washed 3 times (0.5 mL, PBS + *w/v* 0.3% BSA), with re-pulling down and the supernatant was removed between each wash. Bead-associated radioactivity of the washed pull down was determined by using a gamma counter and the relative bead-bound fraction was determined relative to the loading standard. All reactions were performed in triplicate and the mean was determined. Results are summarised in **Table S11**.

**Table S11.** Relative associated activities (RAA) in % of the standard loading dose for all the evaluated [$^{68}$Ga]GaNODAGA-DARPins.

| | [$^{68}$Ga]GaNODAGA-DARPin **3** RAA/ % | [$^{68}$Ga]GaNODAGA-DARPin **6** RAA/ % | [$^{68}$Ga]GaNODAGA-DARPin **9** RAA/ % | [$^{68}$Ga]GaNODAGA-DARPin **12** RAA/ % | [$^{68}$Ga]GaNODAGA-DARPin **13** RAA/ % |
|---|---|---|---|---|---|
| binding | 47.61 ± 1.27 | 37.03 ± 0.52 | 38.60 ± 1.05 | 39.64 ± 1.35 | 4.75 ± 0.99 |
| no PSA | 7.10 ± 2.26 | 4.72 ± 0.88 | 11.81 ± 3.42 | 13.23 ± 0.51 | 3.33 ± 1.77 |
| block | 5.56 ± 0.88 | 3.73 ± 1.28 | 11.67 ± 1.36 | 13.82 ± 0.87 | 2.51 ± 0.52 |

**Tumor uptake studies and *ex vivo* biodistribution studies of [$^{68}$Ga]GaNODAGA-DARPin radiotracers in LNCaP xenografts**

NODAGA-DARPins (~1.30 nmol) were buffered in NaOAc (100 µL, 0.1 M, pH 4.4). Radiolabelling was achieved by addition of an aliquot of [$^{68}$Ga][Ga(H$_2$O)$_6$]Cl$_3$ (50 µL) and adjustment of the pH to >5.3 with Na$_2$CO$_3$ (10 µL, 1M). After incubation of the reaction for 10 min. at room temperature, the [$^{68}$Ga]GaNODAGA-DARPin complexes were analysed by radio-TLC and radio-SEC. The crude reactions were purified *via* NAP-5 SEC chromatography to obtain the purified radiotracer in >99% RCP in PBS (pH 7.4) with decay-corrected RCYs of 57 – 78% ($n$ = 2). Tumour uptake studies were conducted in mouse models with LNCaP xenografts. Tumour-bearing mice were randomised before the study with $n$ = 3 – 4, animals per group. Doses of [$^{69}$Ga]GaNODAGA-DARPin were administered *via* tail-vein injection (150 µL in sterile PBS, amounts **Table S12**). A competitive blocking study was performed with a lower molar activity [$^{69}$Ga]GaNODAGA-DARPin, produced by supplementing the normal formulation with non-labelled DARPin (blocking group, 150 µL in sterile PBS amounts given in **Table S12**).

**Table S12.** Summary of the decay corrected radiochemical yields (d.c. RCY / %) for the isolated, formulated radiotracers after NAP-5 purification ($n$ = 2) and the amounts (µg, nmol, and MBq), and the molar activity (MBq nmol$^{-1}$) of the administered radiotracers. Values are displayed as geometric mean ± one standard deviation.

| | [$^{68}$Ga]GaNODAGA-DARPin **3** | | [$^{68}$Ga]GaNODAGA-DARPin **6** | | [$^{68}$Ga]GaNODAGA-DARPin **9** | | [$^{68}$Ga]GaNODAGA-DARPin **12** | | [$^{68}$Ga]GaNODAGA-DARPin **13** | |
|---|---|---|---|---|---|---|---|---|---|---|
| D.c. RCY / % ($n$ = 2) | 77.86 ± 1.10 | | 71.03 ± 8.32 | | 56.67 ± 3.90 | | 63.89 ± 2.85 | | 70.60 ± 1.13 | |
| | Normal group ($n$ = 4) | Blocking group ($n$ = 3) | Normal group ($n$ = 3) | Blocking group ($n$ = 3) | Normal group ($n$ = 4) | Blocking group ($n$ = 3) | Normal group ($n$ = 3) | Blocking group ($n$ = 4) | Normal group ($n$ = 4) | - |
| Amount / µg | 3.00 | 598.96 | 3.84 | 601.14 | 2.37 | 322.44 | 3.03 | 392.17 | 3.74 | - |
| Amount / nmol | 0.21 | 31.79 | 0.20 | 31.90 | 0.20 | 27.84 | 0.20 | 20.81 | 0.20 | - |
| Activity / MBq | 2.88 ± 0.29 | 2.66 ± 0.27 | 2.40 ± 0.17 | 2.70 ± 0.17 | 1.56 ± 0.18 | 1.875 ± 0.158 | 1.70 ± 0.18 | 2.40 ± 0.25 | 2.18 ± 0.26 | - |
| Molar. act. / MBq nmol$^{-1}$ | 14.02 ± 1.40 | 0.084 ± 0.009 | 11.79 ± 0.84 | 0.085 ± 0.005 | 7.60 ± 0.86 | 0.067 ± 0.006 | 8.36 ± 0.88 | 0.116 ± 0.012 | 10.64 ± 1.25 | - |

##### Effective half-life measurement

The total internal radioactivity over time was measured using a dose calibrator. With this measurement, the effective half-lives ($t_{1/2}$(eff)) of [$^{68}$Ga]GaNODAGA-DARPins were calculated by using a one-phase decay model in GraphPad Prism.

**Table S13.** Total internal radioactivity relative to internal activity at $t$ = 5, 30, 60 and 120 min. of [$^{68}$Ga]GaNODAGA-DARPins in athymic nude mice bearing s.c. LNCaP xenografts and the calculated $t_{1/2}$ (eff) values obtained from a one-phase decay model. Note: Data were analysed in GraphPad Prism.

| | [$^{68}$Ga]GaNODAGA-DARPin **3** Relative internal activity | | [$^{68}$Ga]GaNODAGA-DARPin **6** Relative internal activity | | [$^{68}$Ga]GaNODAGA-DARPin **9** Relative internal activity | | [$^{68}$Ga]GaNODAGA-DARPin **12** Relative internal activity | | [$^{68}$Ga]GaNODAGA-DARPin **13** Relative internal activity | |
|---|---|---|---|---|---|---|---|---|---|---|
| Time / min. | Normal group ($n$ = 4) | Blocking group ($n$ = 3) | Normal group ($n$ = 3) | Blocking group ($n$ = 3) | Normal group ($n$ = 4) | Blocking group ($n$ = 3) | Normal group ($n$ = 3) | Blocking group ($n$ = 4) | Normal group ($n$ = 4) | - |
| 5 | 1.00 | 1.00 | 1.00 | 1 | 1 | 1 | 1 | 1 | 1 | - |
| 30 | 0.71 ± 0.03 | 0.74 ± 0.13 | 0.69 ± 0.02 | 0.73 ± 0.09 | 0.74 ± 0.02 | 0.65 ± 0.06 | 0.70 ± 0.06 | 0.74 ± 0.11 | 0.83 ± 0.05 | - |
| 60 | 0.57 ± 0.04 | 0.49 ± 0.06 | 0.48 ± 0.04 | 0.51 ± 0.07 | 0.51 ± 0.01 | 0.50 ± 0.03 | 0.42 ± 0.02 | 0.51 ± 0.05 | 0.59 ± 0.02 | - |
| 120 | 0.28 ± 0.02 | 0.27 ± 0.05 | 0.27 ± 0.04 | 0.29 ± 0.05 | 0.27 ± 0.01 | 0.26 ± 0.01 | 0.22 ± 0.01 | 0.24 ± 0.02 | 0.31 ± 0.02 | - |
| $t_{1/2}$ (eff) / min. | 61.26 ± 12.79 | 55.92 ± 22.02 | 39.21 ± 4.75 | 51.98 ± 16.95 | 52.48 ± 3.61 | 38.65 ± 7.38 | 40.04 ± 5.55 | 64.55 ± 20.24 | 112.59 ± 32.29 | - |

##### *Ex vivo* biodistribution studies

Animals were anesthetised by inhalation of 2-3% isoflurane (Baxter Healthcare, Deerfield, IL) / oxygen gas mixture (~5 L min$^{-1}$) and were euthanised by exsanguination under anaesthesia. Tissue samples were removed, rinsed in water, dried in air for ~1 min., weighed and counted on a gamma-counter for accumulation of $^{69}$Ga-radioactivity. Tumour uptake values (% ID g$^{-1}$) are given in **Table S14**. Tumour-to-tissue ratios are given in **Table S15.** The statistical analysis of the comparisons of the different groups is represented in **Table S16** – **Table S18** ($P$-values of unpaired, two-tailed Student's $t$-test).

**Table S14.** Biodistribution data showing the accumulation of [$^{68}$Ga]GaNODAGA-DARPins **3**, **6**, **9**, **12**, and **13** in different tissues at 2 h post-administration in the normal and blocking groups of athymic nude mice bearing s.c. LNCaP xenografts mice. The tissue associated uptake is represented in %ID g$^{-1}$. Note: S.D. = 1 standard deviation. Grey shading indicates that *n* = 3, as one value had to be excluded from the experiment.

| | [$^{68}$Ga]GaNODAGA-DARPin **3** (2 h), %ID g$^{-1}$ ± S.D | | [$^{68}$Ga]GaNODAGA-DARPin **6** (2 h), %ID g$^{-1}$ ± S.D | | [$^{68}$Ga]GaNODAGA-DARPin **9** (2 h), %ID g$^{-1}$ ± S.D | | [$^{68}$Ga]GaNODAGA-DARPin **12** (2 h), %ID g$^{-1}$ ± S.D | | [$^{68}$Ga]GaNODAGA-DARPin **13** (2 h), %ID g$^{-1}$ ± S.D | |
|---|---|---|---|---|---|---|---|---|---|---|
| Tissue | Normal group (*n* = 4) | Blocking group (*n* = 3) | Normal group (*n* = 3) | Blocking group (*n* = 3) | Normal group (*n* = 4) | Blocking group (*n* = 3) | Normal group (*n* = 3) | Blocking group (*n* = 4) | Normal group (*n* = 4) | - |
| Blood | 1.71 ± 0.16 | 1.33 ± 0.25 | 7.13 ± 1.25 | 2.69 ± 0.13 | 6.74 ± 1.14 | 4.56 ± 0.04 | 2.10 ± 0.42 | 0.92 ± 0.11 | 5.53 ± 0.64 | - |
| Tumour | 2.39 ± 0.43 | 1.48 ± 0.19 | 4.16 ± 0.58 | 2.47 ± 0.42 | 4.72 ± 1.07 | 2.9 ± 0.46 | 1.80 ± 0.59 | 1.58 ± 0.38 | 2.93 ± 0.55 | - |
| Heart | 0.91 ± 0.07 | 0.67 ± 0.13 | 2.81 ± 0.81 | 1.94 ± 0.08 | 2.01 ± 1.07 | 1.98 ± 0.23 | 1.20 ± 0.24 | 0.70 ± 0.05 | 1.94 ± 0.11 | - |
| Lungs | 1.28 ± 0.12 | 0.91 ± 0.18 | 3.69 ± 0.87 | 2.24 ± 0.18 | 4.18 ± 0.67 | 2.98 ± 0.20 | 1.59 ± 0.54 | 0.71 ± 0.14 | 3.90 ± 1.10 | - |
| Liver | 21.31 ± 3.24 | 14.25 ± 2.74 | 33.16 ± 7.93 | 29.66 ± 5.56 | 32.65 ± 2.31 | 29.02 ± 4.78 | 20.41 ± 3.99 | 10.67 ± 1.44 | 43.04 ± 3.41 | - |
| Spleen | 6.85 ± 0.99 | 3.58 ± 0.86 | 18.07 ± 7.10 | 8.57 ± 1.13 | 12.56 ± 2.91 | 21.62 ± 1.35 | 12.89 ± 5.24 | 2.68 ± 0.85 | 14.99 ± 2.40 | - |
| Stomach | 0.49 ± 0.04 | 0.51 ± 0.06 | 0.76 ± 0.09 | 0.73 ± 0.06 | 0.54 ± 0.21 | 0.95 ± 0.53 | 0.66 ± 0.14 | 0.67 ± 0.32 | 1.31 ± 0.57 | - |
| Pancreas | 0.85 ± 0.08 | 0.57 ± 0.09 | 1.80 ± 0.34 | 1.64 ± 0.04 | 1.74 ± 0.08 | 1.34 ± 0.18 | 1.39 ± 0.31 | 0.58 ± 0.10 | 1.17 ± 0.66 | - |
| Kidney | 139.28 ± 65.59 | 116.50 ± 19.07 | 4.44 ± 0.66 | 148.80 ± 25.75 | 36.13 ± 4.28 | 80.85 ± 10.27 | 149.63 ± 25.00 | 139.03 ± 23.11 | 35.66 ± 5.02 | - |
| Sm. Int. | 1.03 ± 0.36 | 1.65 ± 1.03 | 2.74 ± 1.09 | 1.47 ± 0.11 | 2.12 ± 0.67 | 2.75 ± 1.68 | 1.31 ± 0.39 | 1.70 ± 1.22 | 3.05 ± 1.85 | - |
| Large Int. | 0.60 ± 0.04 | 0.59 ± 0.15 | 1.08 ± 0.09 | 1.19 ± 0.08 | 1.47 ± 0.43 | 1.16 ± 0.08 | 0.70 ± 0.22 | 0.54 ± 0.18 | 1.55 ± 0.59 | - |
| Fat | 0.65 ± 0.40 | 0.63 ± 0.38 | 0.78 ± 0.14 | 1.48 ± 0.88 | 0.89 ± 0.21 | 1.05 ± 0.13 | 0.46 ± 0.20 | 0.47 ± 0.17 | 0.77 ± 0.06 | - |
| Muscle | 0.37 ± 0.03 | 0.30 ± 0.11 | 0.90 ± 0.10 | 0.88 ± 0.22 | 0.82 ± 0.09 | 0.72 ± 0.19 | 0.49 ± 0.10 | 0.33 ± 0.08 | 0.70 ± 0.17 | - |
| Bone | 1.26 ± 0.17 | 0.83 ± 0.12 | 2.64 ± 0.81 | 2.32 ± 0.73 | 3.15 ± 0.64 | 2.77 ± 1.43 | 2.35 ± 0.39 | 1.11 ± 0.56 | 2.32 ± 0.94 | - |
| Skin | 1.76 ± 0.31 | 1.29 ± 0.05 | 2.35 ± 0.43 | 2.55 ± 0.04 | 2.05 ± 0.27 | 1.80 ± 0.12 | 2.29 ± 0.26 | 1.98 ± 0.63 | 1.74 ± 0.10 | - |

**Table S15.** Tumour-to-tissue contrast data following the biodistribution of [$^{68}$Ga]GaNODAGA-DARPins **3**, **6**, **9**, **12**, and **13** in the normal and blocking groups in athymic nude mice bearing s.c. LNCaP xenografts. Note: S.D. = 1 standard deviation, grey: *n* = 3, as one value had to be excluded.

| | [$^{68}$Ga]GaNODAGA-DARPin **3** (2 h), tumour to tissue contrast ratio ± S.D | | [$^{68}$Ga]GaNODAGA-DARPin **6** (2 h), tumour to tissue contrast ratio ± S.D | | [$^{68}$Ga]GaNODAGA-DARPin **9** (2 h), tumour to tissue contrast ratio ± S.D | | [$^{68}$Ga]GaNODAGA-DARPin **12** (2 h), tumour to tissue contrast ratio ± S.D | | [$^{68}$Ga]GaNODAGA-DARPin **13** (2 h), tumour to tissue contrast ratio ± S.D | |
|---|---|---|---|---|---|---|---|---|---|---|
| Tissue | Normal group (*n* = 4) | Blocking group (*n* = 3) | Normal group (*n* = 3) | Blocking group (*n* = 3) | Normal group (*n* = 4) | Blocking group (*n* = 3) | Normal group (*n* = 3) | Blocking group (*n* = 4) | Normal group (*n* = 4) | - |
| Blood | 1.39 ± 0.15 | 1.12 ± 0.08 | 0.58 ± 0.20 | 0.91 ± 0.11 | 0.70 ± 0.09 | 0.64 ± 0.10 | 0.85 ± 0.18 | 1.72 ± 0.38 | 0.54 ± 0.14 | - |
| Tumour | 1.00 | 1.00 | 1.00 | 1.00 | 1.00 | 1.00 | 1.00 | 1.00 | 1.00 | - |
| Heart | 2.63 ± 0.43 | 2.23 ± 0.18 | 1.48 ± 0.47 | 1.27 ± 0.20 | 3.17 ± 2.28 | 1.47 ± 0.22 | 1.51 ± 0.43 | 2.24 ± 0.44 | 1.50 ± 0.21 | - |
| Lungs | 1.85 ± 0.20 | 1.64 ± 0.17 | 1.13 ± 0.31 | 1.10 ± 0.11 | 1.14 ± 0.22 | 0.97 ± 0.11 | 1.19 ± 0.38 | 2.27 ± 0.59 | 0.78 ± 0.20 | - |
| Liver | 0.11 ± 0.02 | 0.11 ± 0.03 | 0.13 ± 0.03 | 0.08 ± 0.02 | 0.15 ± 0.03 | 0.10 ± 0.02 | 0.09 ± 0.02 | 0.15 ± 0.04 | 0.07 ± 0.01 | - |
| Spleen | 0.35 ± 0.06 | 0.43 ± 0.15 | 0.23 ± 0.10 | 0.29 ± 0.01 | 0.38 ± 0.08 | 0.13 ± 0.01 | 0.15 ± 0.04 | 0.63 ± 0.20 | 0.20 ± 0.05 | - |
| Stomach | 4.87 ± 1.04 | 2.91 ± 0.18 | 5.49 ± 1.01 | 3.38 ± 0.31 | 9.12 ± 2.28 | 3.60 ± 1.68 | 2.82 ± 1.02 | 2.78 ± 1.19 | 2.61 ± 1.33 | - |
| Pancreas | 2.80 ± 0.33 | 2.59 ± 0.07 | 2.31 ± 0.54 | 1.50 ± 0.24 | 2.72 ± 0.60 | 2.21 ± 0.54 | 1.32 ± 0.46 | 2.79 ± 0.71 | 4.15 ± 4.17 | - |
| Kidney | 0.02 ± 0.01 | 0.01 ± 0.00 | 0.94 ± 0.19 | 0.02 ± 0.00 | 0.13 ± 0.04 | 0.04 ± 0.00 | 0.01 ± 0.00 | 0.01 ± 0.00 | 0.08 ± 0.02 | - |
| Sm. Int. | 2.43 ± 0.53 | 1.21 ± 0.79 | 1.52 ± 0.64 | 1.67 ± 0.18 | 2.38 ± 0.84 | 1.26 ± 0.56 | 1.41 ± 0.36 | 1.31 ± 0.79 | 1.28 ± 0.79 | - |
| Large Int. | 3.98 ± 0.67 | 2.58 ± 0.49 | 3.84 ± 0.63 | 2.06 ± 0.24 | 3.30 ± 0.65 | 2.49 ± 0.25 | 2.80 ± 1.40 | 3.22 ± 1.54 | 2.14 ± 0.89 | - |
| Fat | 4.46 ± 2.14 | 2.81 ± 1.20 | 5.33 ± 1.22 | 2.13 ± 1.24 | 5.39 ± 0.91 | 2.82 ± 0.77 | 5.06 ± 4.05 | 3.70 ± 1.62 | 3.68 ± 1.03 | - |
| Muscle | 6.57 ± 1.49 | 5.15 ± 1.38 | 4.64 ± 0.84 | 2.97 ± 0.99 | 5.84 ± 1.76 | 4.29 ± 1.53 | 3.87 ± 1.78 | 4.98 ± 1.12 | 4.39 ± 1.44 | - |
| Bone | 1.90 ± 0.28 | 1.78 ± 0.07 | 1.58 ± 0.53 | 1.10 ± 0.18 | 1.64 ± 0.53 | 1.20 ± 0.50 | 0.76 ± 0.17 | 1.77 ± 0.90 | 1.63 ± 1.22 | - |
| Skin | 1.39 ± 0.39 | 1.14 ± 0.11 | 1.77 ± 0.41 | 0.97 ± 0.15 | 2.31 ± 0.46 | 1.61 ± 0.19 | 0.82 ± 0.37 | 0.86 ± 0.29 | 1.70 ± 0.38 | - |

**Table S16.** Statistical analysis of tumour uptake. Comparison of [$^{68}$Ga]GaNODAGA-DARPin **3**, **6**, **9**, **12**, and **13** *vs.* blocking studies and versus data obtained for the non-binding control DARPin **13**. *P*-values of unpaired, two-tailed Student's *t*-test, light blue *P* <0.05, medium blue *P* <0.01, dark blue *P* <0.001.

| | [$^{68}$Ga]GaNODAGA-DARPin **3** *P*- values | | | [$^{68}$Ga]GaNODAGA-DARPin **6** *P* -values | | | [$^{68}$Ga]GaNODAGA-DARPin **9** *P* -values | | | [$^{68}$Ga]GaNODAGA-DARPin **12** *P* -values | | |
|---|---|---|---|---|---|---|---|---|---|---|---|---|
| Tissue | Normal vs. Blocking Group | Normal group vs. DARPin **13** | Blocking group vs. DARPin **13** | Normal vs. Blocking Group | Normal group vs. DARPin **13** | Blocking group vs. DARPin **13** | Normal vs. Blocking Group | Normal group vs. DARPin **13** | Blocking group vs. DARPin **13** | Normal vs. Blocking Group | Normal group vs. DARPin **13** | Blocking group vs. DARPin **13** |
| Blood | 0.09702 | 0.00078 | 0.00025 | 0.07526 | 0.34196 | 0.00224 | 0.03115 | 0.12729 | 0.05563 | 0.03455 | 0.00038 | 0.00055 |
| Tumour | 0.01709 | 0.17359 | 0.00834 | 0.01770 | 0.04219 | 0.26351 | 0.03494 | 0.03512 | 0.94992 | 0.59791 | 0.05810 | 0.00836 |
| Heart | 0.06954 | 0.00002 | 0.00019 | 0.20224 | 0.20172 | 0.97336 | 0.96178 | 0.90396 | 0.78926 | 0.06482 | 0.02125 | 0.00003 |
| Lungs | 0.04507 | 0.01690 | 0.01076 | 0.09533 | 0.78346 | 0.05382 | 0.03033 | 0.68382 | 0.19072 | 0.10138 | 0.01734 | 0.00957 |
| Liver | 0.02755 | 0.00009 | 0.00007 | 0.56930 | 0.15188 | 0.03244 | 0.32036 | 0.00339 | 0.01682 | 0.04143 | 0.00141 | 0.00006 |
| Spleen | 0.00610 | 0.00333 | 0.00100 | 0.14316 | 0.53572 | 0.00717 | 0.00399 | 0.24686 | 0.00627 | 0.07492 | 0.57054 | 0.00088 |
| Stomach | 0.73876 | 0.06555 | 0.06761 | 0.64946 | 0.15064 | 0.13569 | 0.31674 | 0.07049 | 0.43532 | 0.95300 | 0.10464 | 0.11210 |
| Pancreas | 0.01054 | 0.40687 | 0.16807 | 0.50175 | 0.16451 | 0.24656 | 0.04791 | 0.18370 | 0.65532 | 0.03533 | 0.57747 | 0.17065 |
| Kidney | 0.54947 | 0.05044 | 0.01440 | 0.01040 | 0.00093 | 0.01478 | 0.00991 | 0.89153 | 0.00809 | 0.59568 | 0.01359 | 0.00220 |
| Sm. Int. | 0.40708 | 0.11419 | 0.25938 | 0.17990 | 0.79220 | 0.18442 | 0.58626 | 0.39633 | 0.83095 | 0.57899 | 0.15430 | 0.27558 |
| Large Int. | 0.91093 | 0.04792 | 0.04207 | 0.20419 | 0.21244 | 0.31691 | 0.24881 | 0.83937 | 0.28324 | 0.36956 | 0.05622 | 0.03644 |
| Fat | 0.94203 | 0.59263 | 0.57902 | 0.30036 | 0.95868 | 0.29866 | 0.26168 | 0.35688 | 0.04516 | 0.91593 | 0.09864 | 0.03169 |
| Muscle | 0.42882 | 0.02525 | 0.01294 | 0.89979 | 0.11505 | 0.31192 | 0.44409 | 0.25741 | 0.89627 | 0.07747 | 0.08716 | 0.01294 |
| Bone | 0.01276 | 0.10899 | 0.04993 | 0.64339 | 0.64747 | 0.99076 | 0.70600 | 0.22263 | 0.66190 | 0.03928 | 0.94824 | 0.08940 |
| Skin | 0.05506 | 0.92484 | 0.00084 | 0.51372 | 0.12774 | 0.00012 | 0.15926 | 0.09973 | 0.55495 | 0.42602 | 0.05489 | 0.50551 |

**Table S17.** Statistical analysis of tissue uptake showing the comparison of [$^{68}$Ga]GaNODAGA-DARPin **3**, **6**, **9** for the normal groups *vs.* the other PSA-binding [$^{68}$Ga]GaNODAGA-DARPin. *P*-values of unpaired, two-tailed Student's *t*-test, light blue *P* <0.05, medium blue *P* <0.01, dark blue *P* <0.001.

| | [$^{68}$Ga]GaNODAGA-DARPin **3** *P*- values | | | [$^{68}$Ga]GaNODAGA-DARPin **6** *P* -values | | | [$^{68}$Ga]GaNODAGA-DARPin **9** *P* -values | | |
|---|---|---|---|---|---|---|---|---|---|
| Tissue | vs. DARPin **6** | vs. DARPin **9** | vs. DARPin **12** | vs. DARPin **9** | vs. DARPin **12** | - | vs. DARPin **12** | - | - |
| Blood | 0.01591 | 0.00268 | 0.24203 | 0.79963 | 0.05598 | - | 0.00168 | - | - |
| Tumour | 0.00739 | 0.01587 | 0.22942 | 0.41247 | 0.00778 | - | 0.00659 | - | - |
| Heart | 0.01952 | 0.13031 | 0.16199 | 0.31077 | 0.06445 | - | 0.22822 | - | - |
| Lungs | 0.00666 | 0.00261 | 0.42988 | 0.46117 | 0.03160 | - | 0.00254 | - | - |
| Liver | 0.02570 | 0.00179 | 0.76375 | 0.92265 | 0.08995 | - | 0.01767 | - | - |
| Spleen | 0.04748 | 0.02386 | 0.18095 | 0.30989 | 0.37125 | - | 0.92771 | - | - |
| Stomach | 0.00227 | 0.66281 | 0.17180 | 0.13585 | 0.34894 | - | 0.43481 | - | - |
| Pancreas | 0.02825 | 0.000005 | 0.08375 | 0.77092 | 0.19686 | - | 0.18667 | - | - |
| Kidney | 0.02720 | 0.05113 | 0.78697 | 0.00050 | 0.00970 | - | 0.01424 | - | - |
| Sm. Int. | 0.06255 | 0.03838 | 0.38266 | 0.44223 | 0.14004 | - | 0.10325 | - | - |
| Large Int. | 0.00038 | 0.02622 | 0.52107 | 0.16799 | 0.07363 | - | 0.02947 | - | - |
| Fat | 0.62365 | 0.34850 | 0.43407 | 0.44159 | 0.08939 | - | 0.04145 | - | - |
| Muscle | 0.00036 | 0.00105 | 0.15258 | 0.39092 | 0.00753 | - | 0.00769 | - | - |
| Bone | 0.01973 | 0.03087 | 0.02674 | 0.43982 | 0.61878 | - | 0.15407 | - | - |
| Skin | 0.19665 | 0.20550 | 0.05877 | 0.35818 | 0.83524 | - | 0.29381 | - | - |

**Table S18.** Statistical analysis of tissue uptake showing the comparison of [$^{68}$Ga]GaNODAGA-DARPin **3**, **6**, **9** from the blocking groups *vs*. the other [$^{68}$Ga]GaNODAGA-DARPin (also blocking groups). *P*-values of unpaired, two-tailed Student's *t*-test, light blue *P* <0.05, medium blue *P* <0.01, dark blue *P* <0.001.

| | [$^{68}$Ga]GaNODAGA-DARPin **3** *P*- values | | | [$^{68}$Ga]GaNODAGA-DARPin **6** *P* -values | | | [$^{68}$Ga]GaNODAGA-DARPin **9** *P* -values | | |
|---|---|---|---|---|---|---|---|---|---|
| Tissue | vs. DARPin **6** - Block | vs. DARPin **9** - Block | vs. DARPin **12** - Block | vs. DARPin **9** - Block | vs. DARPin**12** - Block | - | vs. DARPin **12** - Block | - | - |
| Blood | 0.00360 | 0.00161 | 0.08937 | 0.00068 | 0.00003 | - | 0.000001 | - | - |
| Tumour | 0.03824 | 0.02115 | 0.66540 | 0.29570 | 0.04241 | - | 0.01674 | - | - |
| Heart | 0.00196 | 0.00245 | 0.72667 | 0.77004 | 0.0000003 | - | 0.00844 | - | - |
| Lungs | 0.00083 | 0.00021 | 0.18900 | 0.00958 | 0.00047 | - | 0.00025 | - | - |
| Liver | 0.02433 | 0.01653 | 0.13697 | 0.88617 | 0.02286 | - | 0.01679 | - | - |
| Spleen | 0.00464 | 0.00013 | 0.23376 | 0.00025 | 0.00242 | - | 0.00015 | - | - |
| Stomach | 0.01065 | 0.28610 | 0.39659 | 0.54458 | 0.73304 | - | 0.47184 | - | - |
| Pancreas | 0.00054 | 0.00809 | 0.94634 | 0.09375 | 0.00004 | - | 0.00775 | - | - |
| Kidney | 0.16186 | 0.06331 | 0.21852 | 0.03119 | 0.63059 | - | 0.00912 | - | - |
| Sm. Int. | 0.78852 | 0.39668 | 0.95085 | 0.31560 | 0.72504 | - | 0.41825 | - | - |
| Large Int. | 0.00784 | 0.00943 | 0.71762 | 0.66831 | 0.00201 | - | 0.00249 | - | - |
| Fat | 0.23106 | 0.18448 | 0.55098 | 0.48872 | 0.18223 | - | 0.00354 | - | - |
| Muscle | 0.02737 | 0.03935 | 0.79346 | 0.39475 | 0.03775 | - | 0.05193 | - | - |
| Bone | 0.06860 | 0.14282 | 0.48526 | 0.66259 | 0.08927 | - | 0.17179 | - | - |
| Skin | 0.00001 | 0.00924 | 0.11534 | 0.00459 | 0.16907 | - | 0.60288 | - | - |
